# Affinity-Based Interactome Mapping of Inositol Pyrophosphates Reveals 4/6-PP-InsP_5_–Binding Proteins in Plants

**DOI:** 10.1101/2025.10.20.683386

**Authors:** Kevin Ritter, Verena Gaugler, Sara Christina Stolze, Riya Ghosh, Akhila Jayamon, Felix Wollensack, Debabrata Laha, Hirofumi Nakagami, Gabriel Schaaf, Henning Jacob Jessen

## Abstract

Inositol pyrophosphates (PP-InsPs) are central regulators of eukaryotic signaling events. While certain PP-InsP isomers have been conclusively linked to the regulation of phosphate homeostasis through interaction with SPX domain containing proteins in plants, the functions of the recently discovered isomer 4/6-PP-InsP_5_ remain largely unknown. Here, we employed two complementary affinity-based strategies – a matrix approach and a photoaffinity probe – to systematically identify 4/6-PP-InsP_5_-binding proteins in *Arabidopsis thaliana*. The two methods yielded partially overlapping protein sets, with photoaffinity enrichment likely capturing additional transient and/or weak interactions. Moreover, competition experiments with different isomers were applied to obtain information about potential isomer-specific interactions. As a proof-of-concept, one candidate interactor (FHA domain-containing protein AtFHA2) was shown to bind 4-PP-InsP_5_ *in vitro* with substantially higher affinity than InsP_6_. Thus, besides the SPX domain, FHA domain containing proteins, of which 18 exist in Arabidopsis, are potentially regulated by inositol pyrophosphates. More generally, our findings reveal a diverse protein network associated with 4/6-PP-InsP_5_ and establish a versatile platform for dissecting its biological roles in plants and other organisms.

## 1 Introduction

Inositol phosphates (InsPs) are a diverse class of highly charged intracellular signaling molecules derived from *myo*-inositol (**1**), a cyclohexane hexol with a distinct stereochemistry including an internal mirror plane, which classifies it as a *meso*-compound.^[1–3]^ The addition of phosphate and diphosphate groups at different hydroxyl positions generates a vast array of regioisomers and enantiomers through desymmetrization.^[4,5]^ Among those, inositol pyrophosphates (PP-InsPs) represent a densely phosphorylated subset that plays a critical role in cellular regulation, influencing processes such as phosphate homeostasis, energy metabolism, and stress responses across organisms.^[2,3,6,7]^

Research has primarily focused on 5-PP-InsP_5_ (**2**) and 1,5-(PP)_2_-InsP_4_ (**3**) in mammals and plants, but recent studies demonstrated that other PP-InsP isomers (see Figure 1) are widespread and more abundant than previously thought. 6-PP-InsP_5_ (**5**) was initially believed to be unique to *Dictyostelium discoideum*, where it is the predominant PP-InsP_5_ isomer.^[8,9]^ However, recent studies have identified 4/6-PP-InsP_5_ (**4**/**5**) in various eukaryotic systems, including plants, patient-derived peripheral blood mononuclear cells (PBMCs), and mouse colon and heart tissues^[9–12]^. These findings were obtained using capillary electrophoresis–mass spectrometry (CE-MS) with heavy isotope labeled internal references, confirming its occurrence across diverse biological systems ^[10,11]^. Notably, in all studied land plants and PBMCs, 4/6-PP-InsP_5_(**4/5**) was detected at levels comparable to or exceeding those of 5-PP-InsP_5_ (**2**), suggesting a more prominent role than previously assumed.^[9,11,12]^ Since CE-MS does not discriminate between enantiomers, the signal could arise from 4-PP-InsP_5_, 6-PP-InsP_5_ (**4** or **5**), or both. For clarity, we refer to them collectively as 4/6-PP-InsP_5_ (**4/5**). The detection of this isomer beyond *D. discoideum* challenges long-standing assumptions about PP-InsP metabolism and highlights the need to reassess the functional significance of 4/6-PP-InsP_5_ (**4/5**) in eukaryotic signaling pathways.

**Figure 1:**
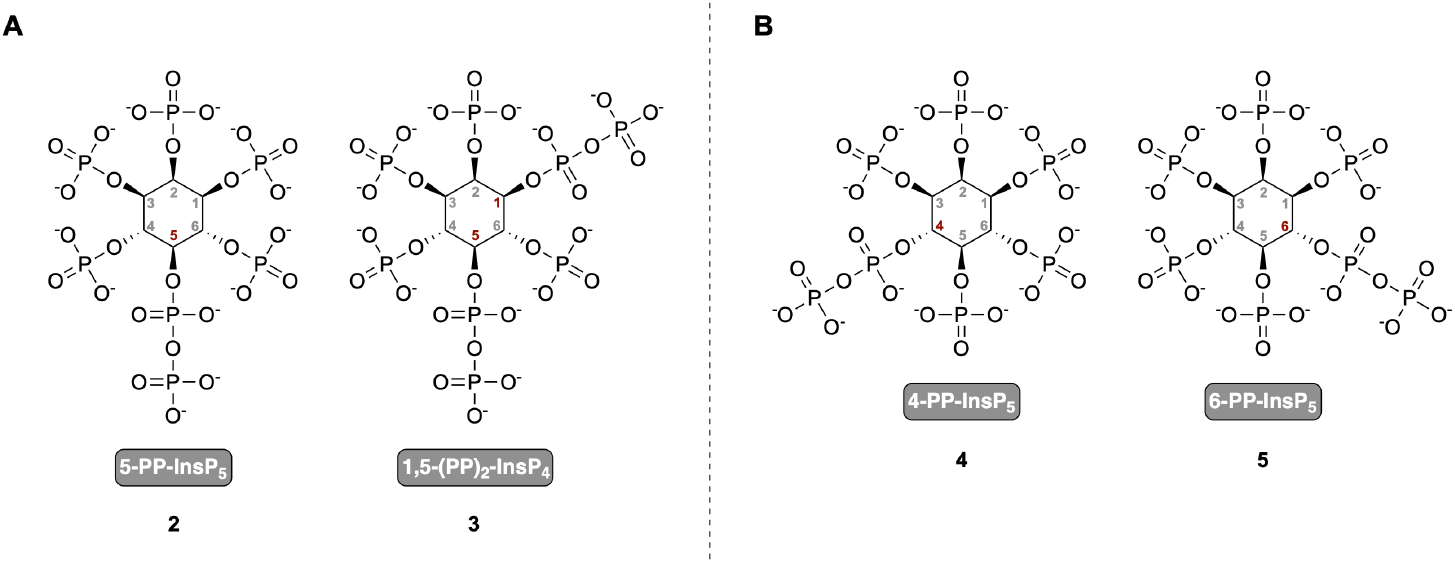
Chemical structures of selected inositol pyrophosphates. a) 5-PP-InsP_5_ (**2**) and 1,5-(PP)_2_-InsP_4_ (**3**), the two most extensively studied PP-InsPs in mammalian and plant systems. b) 4-PP-InsP_5_ (**4**) and 6-PP-InsP_5_ (**5**) are enantiomers and were recently identified in plants, mammalian cells, and other eukaryotic species, while the exact configuration remains unknown.

In *Arabidopsis thaliana*, the inositol polyphosphate multikinase AtIPK2α and AtIPK2β phosphorylate InsP_6_ (**6**) to generate 4/6-PP-InsP_5_ (**4/5**) *in vitro*.^[12]^ Together they regulate the cellular levels of 4/6-PP-InsP_5_ (**4/5**) *in planta*. Notably, these kinases play a critical role in heat stress acclimation, as their disruption leads to impaired expression of heat shock proteins and reduced thermo-tolerance.^[12]^ The evolutionary conservation of this function is supported by findings in *Marchantia polymorpha*, where an IPMK homolog contributes to heat stress responses, suggesting an ancient role of 4/6-PP-InsP_5_ (**4/5**) in environmental adaptation.^[12]^ Beyond heat stress, PP-InsPs regulate phosphate homeostasis via SPX-domain-containing proteins, which mediate phosphate starvation responses.^[13]^ Additionally, certain NUDIX hydrolases selectively degrade 4-PP-InsP_5_ (**4**), suggesting an isomer-specific regulatory mechanism.^[14,15]^ Taken together, these insights highlight the need to further investigate the specific roles of 4/6-PP-InsP_5_ (**4/5**) in plant signaling networks, particularly its potential impact on stress adaptation.

Despite the growing recognition of 4/6-PP-InsP_5_ (**4/5**) as a functional signaling molecule, its protein interactome remains largely unexplored. Previous affinity enrichment studies have been limited to 5-PP-InsP_5_ (**2**) and 1,5-(PP)_2_-InsP_4_ (**3**) in non-plant systems, expanding our understanding of PP-InsP interactomes and providing new candidates for functional studies in yeast and mammalian cells.^[16–18]^ In Arabidopsis, affinity enrichment experiments have so far been conducted exclusively for 5-PP-InsP_5_ (**2**), using an Affi-Gel method, in which a nonhydrolyzable 5-PCP-InsP_5_ (**7**) analog is immobilized on a resin matrix to enable selective protein binding.^[14,15]^ Given the structural differences between PP-InsP isomers and their potential for distinct protein interactions,^[6,19]^ a targeted approach to characterize the interactors of 4/6-PP-InsP_5_ (**4/5**) is necessary.

To address this gap, we applied complementary affinity-based enrichment strategies to identify 4/6-PP-InsP_5_-binding proteins in the flowering plant *Arabidopsis thaliana*. By combining a matrix-based approach with a photoaffinity labeling method, we systematically mapped the ligand’s interactome. Our study provides a robust methodological framework for investigating 4/6-PP-InsP_5_ signaling in plants and beyond.

## 2 Results & Discussion

### Synthesis of Amino-PEG-4/6-PCP-InsP_5_

To enable the selective enrichment of 4/6-InsP_5_-binding proteins, we developed a modular inositol pyrophosphate analog bearing a terminal amine suitable for covalent modification or resin attachment. The resulting compound, Amino-PEG-4/6-PCP-InsP_5_ (**14**), consists of a methylene bisphosphonate (PCP) at either the 4- or 6-position of *myo*-inositol (**1**), and a polyethylene glycol (PEG) linker with a terminal amine installed on the opposite phosphate. That is, when the PCP group is located at position 4, the PEG linker is attached at position 6, and *vice versa*. As the 4- and 6-positions are enantiotopic, and it is not clear whether 4- or 6-PP-InsP_5_ (**4** or **5**) is the biologically relevant isomer, the compound was synthesized and used as a racemic mixture of 4- and 6-PCP-isomers.

Our compound design builds on an affinity enrichment strategy established by Wu et al., who immobilized a nonhydrolyzable methylene bisphosphonate (PCP) analog of 5-PP-InsP_5_ (**2**) on Affi-Gel resin for pull-down experiments.^[16]^ PCP analogs are chemically stabilized diphosphate mimics that preserve the geometry and charge of native PP-groups while resisting hydrolysis and eliminating phosphoryl transfer, making them powerful tools for probing PP-InsP signaling.^[20,21]^ Other stabilized diphosphate mimics, including α-phosphonoacetic acid (PA) esters and difluoro-substituted analogues, have also been developed to retain non-covalent recognition while blocking phosphoryl transfer, but they have not yet been used in enrichment workflows.^[22,23]^ To address this gap, we synthesized a PCP-containing analog of 4/6-PP-InsP_5_ (**4/5**) designed for both resin coupling and photoaffinity labeling.

The starting point for this synthesis was a previously established strategy for the regioselective functionalization of the 4- and 6-positions of *myo*-inositol (**1**) (see Scheme 1).^[24]^ In the first step, *myo*-inositol (**1**) was protected as its orthoformate using triethyl orthoformate under acidic conditions. Selective silylation at position 2 with TBSCl and a sterically hindered base (2,6-lutidine) enabled differentiation between axial and equatorial hydroxyl groups.^[24]^ Allyl groups were then introduced at positions 4 and 6 to allow for orthogonal deprotection in later steps, affording the bis-allylated inositol derivative **8**.

Acid treatment removed both the orthoformate and the TBS group, releasing hydroxyl groups at positions 1, 2, 3, and 5. These were phosphorylated using a standard phosphoramidite protocol with dibenzyl phosphoramidite and 4,5-dicyanoimidazole (DCI) as activator, followed by oxidation with *meta*-chloroperbenzoic acid (*m*CPBA).^[4,25,26]^ The resulting tetraphosphate intermediate **10** was then subjected to PdCl_2_-mediated deallylation, releasing hydroxyl groups at positions 4 and 6.^[26]^ As prolonged reaction times led to phosphate migration, the reaction was carefully timed and monitored via ^31^P-NMR. Following phosphate migration, the resulting regioisomeric mixtures cannot be resolved, making it essential to prevent their formation.

The PCP group was introduced using PCP-phosphoramidite (**S2**) synthesized according to Hostachy et al.^[27]^ As the substitution occurs at either the 4- or 6-position of the inositol, and each phosphorylation step creates a stereogenic center at phosphorus upon oxidation, a total of four stereoisomers are formed – two diastereomers, each present as a pair of enantiomers. Two distinct species were observed both by ^31^P-NMR spectroscopy, with signals in a ratio of approximately 2:3, and by LC-MS, which showed closely eluting peaks of identical mass. Separation of the diastereomers by chromatography was not attempted, as the stereogenic centers at phosphorus collapse upon global deprotection. Their transient formation nevertheless confirmed the successful and selective incorporation of the PCP group.

The final phosphate diester, bearing a PEG linker with a terminal primary amine, was installed at the remaining free hydroxyl group using the phosphoramidite approach. The PEG-phosphoramidite was synthesized following the procedure reported by Wu et al.^[16]^ Upon oxidation, the phosphorus atom of the newly introduced group becomes stereogenic, adding an additional layer of stereochemical complexity to the molecule. Consequently, prior to global deprotection, up to eight stereoisomers – corresponding to four pairs of enantiomers – are theoretically possible. However, because of signal overlap and the unequal formation of individual species, the resulting diastereomers could not be fully resolved by NMR or LC-MS. Subsequent catalytic hydrogenation over Pd/C effected global debenzylation in a single step, eliminating the complex stereochemical mixture generated earlier. The target compound, Amino-PEG-4/6-PCP-InsP_5_ (**14**), was obtained as a mixture of the two enantiomers with substitution at either the 4- or 6-position of *myo*-inositol. This molecule served as a precursor for both affinity enrichment strategies described in this study. The free amine allowed direct coupling to NHS-activated Affi-Gel resin, as previously shown for the PCP analog of 5-PP-InsP_5_ (**2**),^[16]^ and was also compatible with trifunctional photoaffinity linkers. Applying this strategy to 4/6-PP-InsP_5_ (**4/5**) provided the basis for identifying stereoisomer-specific protein interactors in plants.

**Scheme 1:**
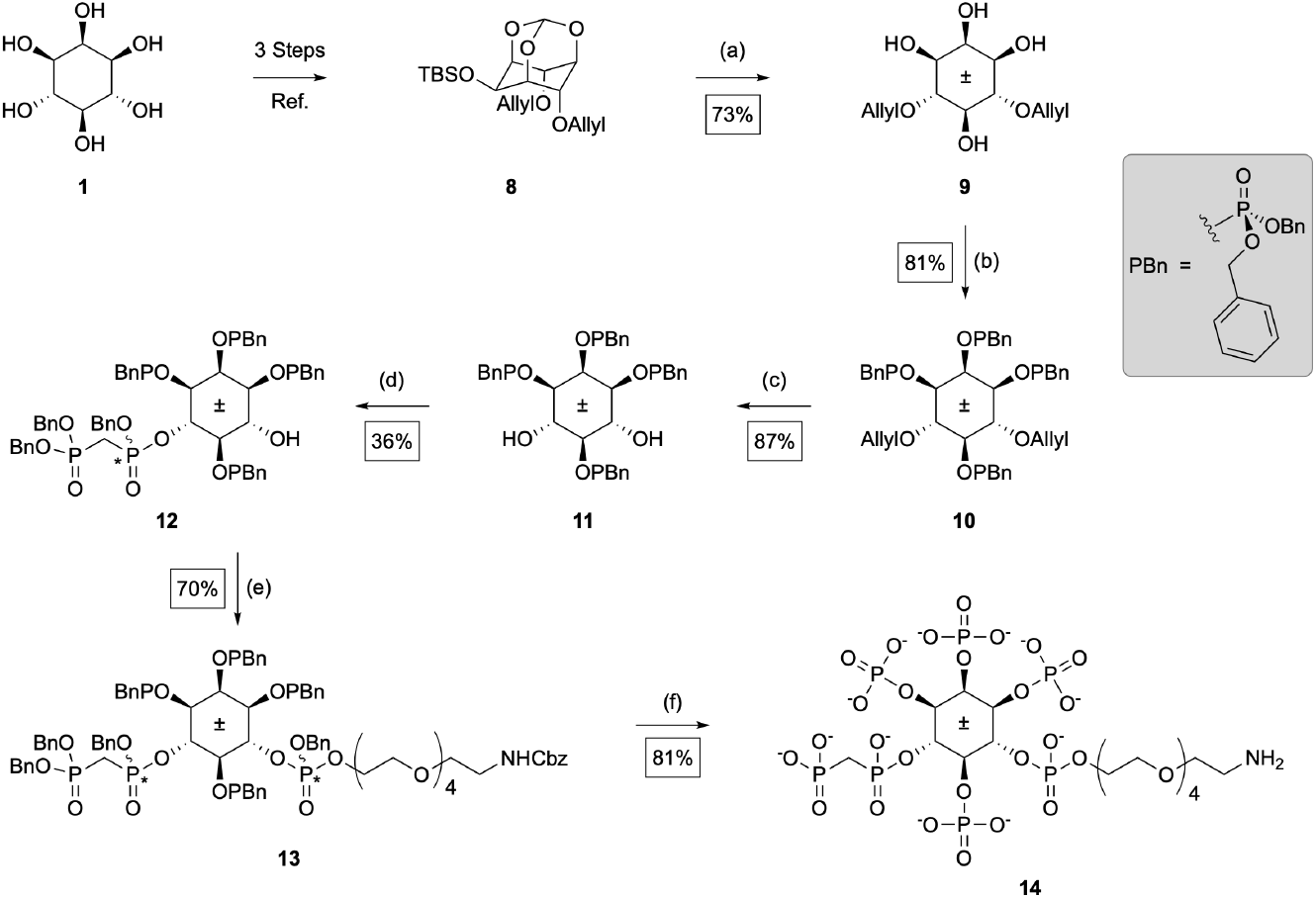
Synthesis of Amino-PEG-4/6-PCP-InsP_5_ (**14**). Reagents and conditions: (**a**) NaH (2.6 eq), allyl bromide (2.6 eq), NaI (cat.), DMF, 0°C to rt, overnight. (**b**) Bn-PA (7.0 eq), DCI (7.0 eq), DMF, rt, 2h; *then m*CPBA (7.0 eq), 0°C to rt, 10 min. (**c**) PdCl_2_ (2.0 eq), MeOH, rt, 2h. (**d**) PCP-PA (1.2 eq), DCI (2.0 eq), CH_2_Cl_2_ rt, 1.5h; *then m*CPBA (2.0 eq), 0°C to rt, 10 min. (**e**) Fm-DiPA (6.2 eq), PEG-linker-alcohol (6.2 eq), ETT (6.2 eq); *then* **12** (1.0 eq), ETT (3.0 eq), CH_2_Cl_2_ rt; *then m*CPBA (3.0 eq), 0°C to rt, 10 min. (**f**) H_2_ (30 bar), Pd/C (3.0 eq), NaHCO_3_ (13.0 eq), *t*BuOH/H_2_O (40:7), rt, 21 h. **Abbreviations:** Bn-PA = Bis-benzyl-*N,N*-diisopropylamino phosphoramidite; DCI = 4,5-Dicyanoimidazol; ETT = 5-Ethylthio-1*H*-tetrazole; Fm-DiPA = 9*H*-fluoren-9-ylmethyl-bis(*N,N*-diisopropylamino) phosphordiamidite. *m*CPBA = meta-chloroperoxybenzoic acid; PA= phosphoramidite; PCP = methylenebisphosphonate.

### Synthesis of a Trifunctional-Photoaffinity Compound

Photoaffinity capture enables covalent crosslinking of ligand-binding proteins upon UV activation, allowing detection of transient or low-affinity interactions that are not readily covered by conventional pull-down approaches.^[28,29]^ To apply this strategy to 4/6-PP-InsP_5_ (**4/5**), a suitable linker must combine three essential features: a photoreactive group for UV-induced crosslinking, a biotin-based tag for streptavidin-mediated enrichment, and an activated ester for coupling to the amino-functionalized probe.

Biotin and desthiobiotin both form strong non-covalent interactions with streptavidin, enabling efficient recovery of labeled protein complexes via immobilized streptavidin matrices, including agarose or magnetic beads.^[28,30]^ A commonly used linker that fulfills these requirements is Sulfo-SBED (Thermo Fisher Scientific), which integrates an aryl azide, biotin, and a cleavable disulfide bridge. However, this reagent is costly and incompatible with reducing agents such as DTT, which are often used in lysate preparations.

To overcome these limitations, we designed a custom photoaffinity linker that retained the essential functional elements of Sulfo-SBED but replaced the disulfide bridge with a stable backbone and featured desthiobiotin instead of biotin. The custom linker (**19**, see Scheme 2A) was synthesized in seven steps, starting from a commercially available Boc-protected lysine methyl ester (**15**). In the first step, the photoreactive aryl azide was introduced via coupling with 4-azidobenzoic acid under standard peptide coupling conditions (HOBt, EDCI, NEt_3_), affording intermediate **16** in 94% yield. After basic hydrolysis of the methyl ester, the resulting carboxylic acid was coupled with aminohexanoic methyl ester (**S8**) to give compound **17** in 70% yield. Aminohexanoic methyl ester (**S8**) was synthesized separately following a reported procedure.^[31]^ Subsequent Boc deprotection using trifluoroacetic acid (TFA) afforded the free amine, which was immediately subjected to a third peptide coupling with desthiobiotin, yielding intermediate **18** (68% yield). Final hydrolysis of the methyl ester and *in situ* activation with DCC in DMF furnished the sulfonated NHS ester **19**. The sulfonated NHS ester ensured aqueous solubility, as the final coupling to the Amino-PEG-4/6-PCP-InsP_5_ probe (**14**) had to be carried out in water.

Conjugation of the photoaffinity linkers to the Amino-PEG-4/6-PCP-InsP_5_ probe (**14**) was performed in aqueous sodium bicarbonate buffer under mild conditions, yielding the two fully functionalized capture reagents depicted in Scheme 2B. Conjugation with the custom linker afforded compound **20**, while coupling with the commercially available Sulfo-SBED linker provided compound **21**, both isolated in different protonation degrees with TEAA as counterions after purification. Both reagents were obtained as racemates and were applied in photoaffinity pulldown experiments to identify 4/6-PP-InsP_5_-binding proteins.

**Scheme 2:**
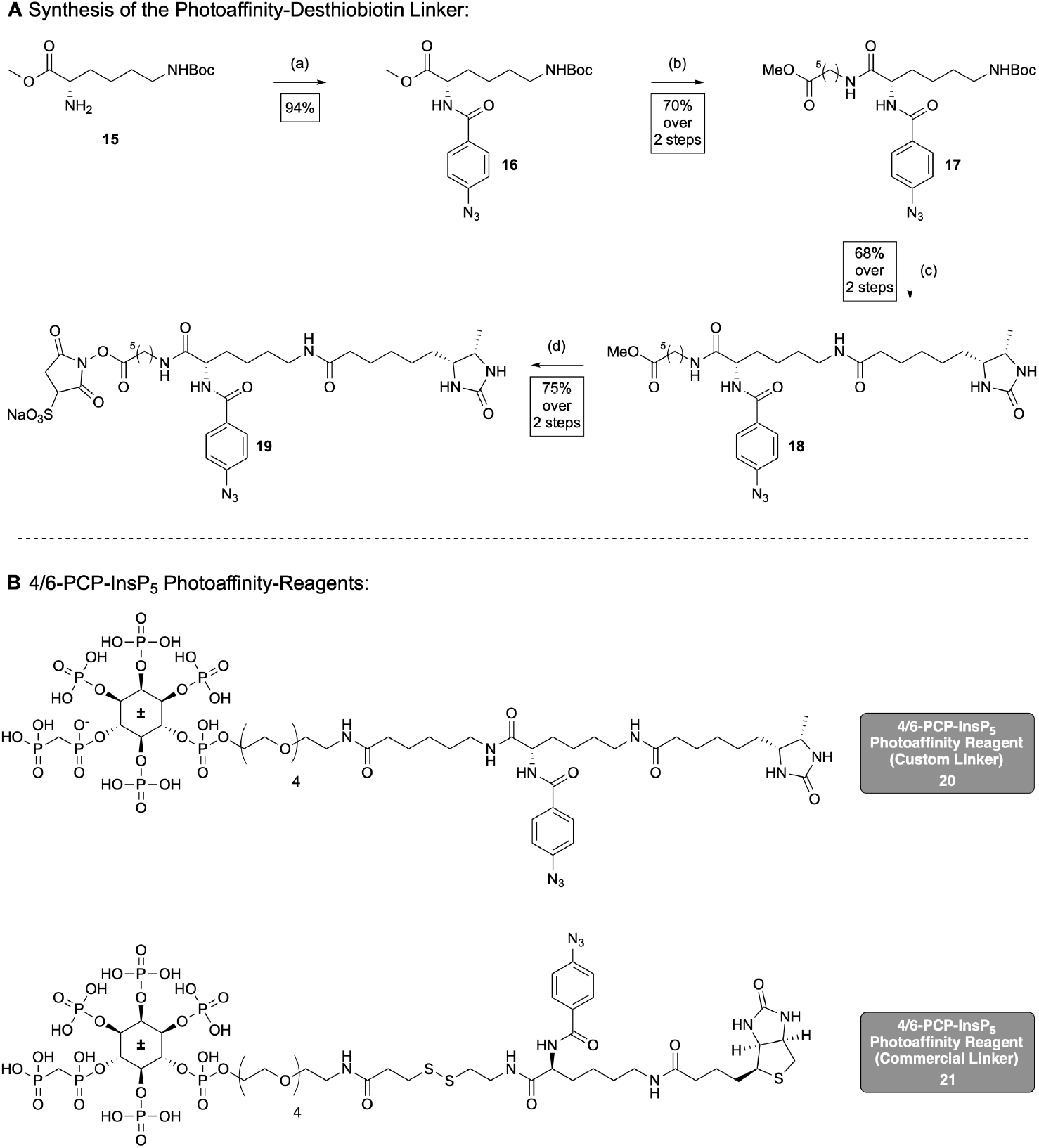
(**A**) Synthesis of custom-photoaffinity linker (**17**). (**B**) Structures of the racemic photoaffinity reagents with custom (**20**)and commercial linker (**21**). Reagents and conditions: (**a**) 4-azodibenzoic acid (1.1 eq), HOBt (1.2 eq), EDCI (1.2), NEt_3_ (3.5 eq), CH_2_Cl_2_, rt, overnight. (**b**) aq. NaOH (2.0 eq), MeOH, 0°C, 30 min; *then* **S8** (1.1 eq), HOBt (1.2 eq), EDCI (1.2 eq), NEt_3_ (3.5 eq), rt, overnight. (**c**) TFA (33 % v/v), CH_2_Cl_2_, rt, 90 min; *then* desthiobiotin (1.1 eq), HOBt (1.5 eq), EDCI (1.5 eq), NEt_3_ (3.0 eq), DMF, rt, overnight. (**d**) aq. NaOH (1M), MeOH, 0°C, 90 min; *then* sulfo-NHS (1.0 eq), DCC (3.0 eq), rt, 48h. **Abbreviations:** HOBt = *N*-Hydroxybenzotriazole; EDCI = 1-Ethyl-3-(3-dimethylaminopropyl)carbodiimide; DCC = *N,N*′-Dicyclohexylcarbodiimid.

### Affinity Enrichment and Proteomic Analysis

Root and shoot material of Arabidopsis was prepared as previously described.^[14]^ Tissue was ground in liquid nitrogen and extracted with magnesium-containing lysis buffer. DTT was included for Affi-Gel and custom linker (**20**) experiments but omitted for commercial linker (**21**)to prevent disulfide reduction. Lysates were clarified by centrifugation and directly used for enrichment.

To probe non-covalent interactions, amino-functionalized PCP analogs of 4/6-PP-InsP_5_ (**4/5**) were immobilized on NHS-activated agarose beads, according to Furkert et al.^[32]^ Negative control matrices contained immobilized inorganic phosphate coupled to the same linker. Beads were incubated with clarified lysates to allow equilibrium binding, then extensively washed. Bound proteins were eluted with 20 mM InsP_6_ (**6**) (elution fraction), while remaining proteins were subjected to on-bead trypsin digestion (on-bead fraction). Both fractions were analyzed by LC-MS/MS. The workflow was adapted from Wu et al. and Schneider et al.^[14,16]^ Comparative enrichment versus control matrices revealed candidate interactors.

In parallel, a photoaffinity-based approach was used to covalently capture protein interactions (see Figure 2), adapted from Haas et al.^[33]^ Two biotinylated capture compounds were applied, both comprising a non-hydrolyzable analog of 4/6-PP-InsP_5_ (**4/5**), a photoreactive group, and a sorting handle. The design and synthesis of both linker systems are described in detail above (see Scheme 2).

**Figure 2:**
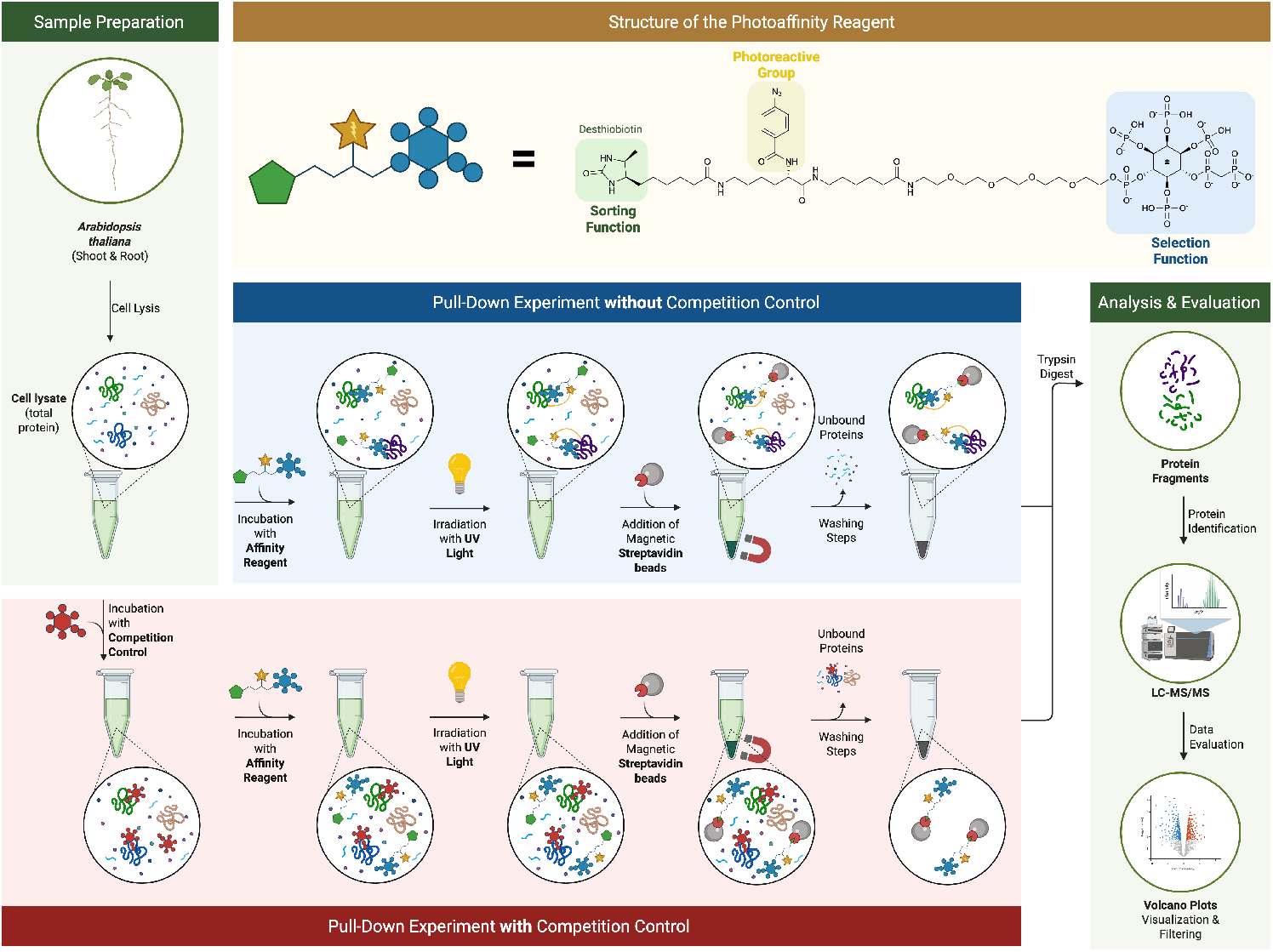
Schematic overview of the photoaffinity-based enrichment workflow used to identify 4/6-PP-InsP_5_-binding proteins in *Arabidopsis thaliana*. Top left: Sample preparation from root and shoot tissue. Top center: Structure of the photoaffinity reagent featuring a photoreactive group, a sorting function, and a selection function. Center panels: Parallel workflows with (red) and without (blue) competition control, including incubation with the photoaffinity reagent, UV crosslinking, and enrichment with streptavidin magnetic beads. Right: Analysis and evaluation steps involving on-bead digestion, LC-MS/MS, and data interpretation using volcano plots. Figure was created with Biorender.com.

Prior to probe addition, lysates were either left untreated or pre-incubated with a 300-fold excess of soluble competitors – enantiopure 4-PP-InsP_5_ (**4**) or 6-PP-InsP_5_ (**5**) – to assess binding specificity and potential enantioselectivity in binding. Capture compounds were added and incubated at 4 °C, followed by UV irradiation (365 nm, 30 min) to induce covalent crosslinking. Resulting complexes were isolated using streptavidin magnetic beads. DTT was included with the custom linker to stabilize interactions, but omitted with the commercial linker to preserve the disulfide bridge. After washing, beads were stored at −80 °C. Candidate interactors were defined as proteins enriched in non-competed samples compared to competitor-treated controls, indicating specific or stereoselective binding.

Proteins from both enrichment approaches were processed using a unified proteomics workflow. Trypsin digestion was performed either on-bead (photoaffinity and Affi-Gel beads) or in-solution (Affi-Gel eluates), followed by desalting and LC-MS/MS analysis on Orbitrap mass spectrometers. Protein identification and label-free quantification were performed with MaxQuant, and statistical analysis of protein enrichment was performed in Perseus using a consistent threshold (S0 = 1, FDR = 0.05).^[34,35]^ Proteins with a log_2_ fold change > 2 were considered candidate interactors. Quantitative filtering was set to a minimum of three valid values per condition for Affi-Gel datasets and two for photoaffinity experiments.

Gene Ontology (GO) enrichment analysis was performed using g:Profiler (default parameters), focusing on categories related to phosphatidylinositol metabolism and inositol phosphate signaling. Shown terms represent Driver Terms as defined by g:Profiler.^[36]^ A condition-specific matrix listing all proteins with log_2_ enrichment > 2 in at least one condition, including gene IDs, annotations, and fold changes, is provided in the Supporting Information. Data processing and visualization were supported by custom R scripts (partly generated via ChatGPT-4o) used to structure the matrix, apply filtering, and generate volcano plots, UpSet diagrams, and GO-term charts.

### Affinity Enrichment: Comparative Results and Key Insights

Comparative analysis of the enrichment datasets provides an integrated view of the 4/6-PP-InsP_5_ interactome in Arabidopsis, revealing how tissue type, enrichment strategy, and competition isomer influence the captured protein subsets.

Affinity-based enrichment using Affi-Gel matrices yielded a moderate but specific set of 4/6-PP-InsP_5_-binding proteins, with about 40–100 proteins enriched per condition (see Figure 3). In addition to direct on-bead digestion, an elution fraction was generated by releasing proteins with InsP_6_ (**6**). The enrichment profiles showed a combination of proteins consistently detected across all conditions and others that appeared only in specific tissues or fractions. Eight proteins were reproducibly enriched in every condition – AT1G07310.1 (CaLB-domain protein), AT1G10900.1 (phosphatidylinositol-5-kinase), AT1G12380.1 (uncharacterized protein), AT1G31440.1 (SH3-domain protein), AT1G47550.1 (SEC3A), AT3G03790.1 (ankyrin/RCC1 repeat protein), AT3G22170.2 (FHY3), and AT4G25550.1 (cleavage and polyadenylation factor) – suggesting that they represent stable, high-affinity interactors.

**Figure 3:**
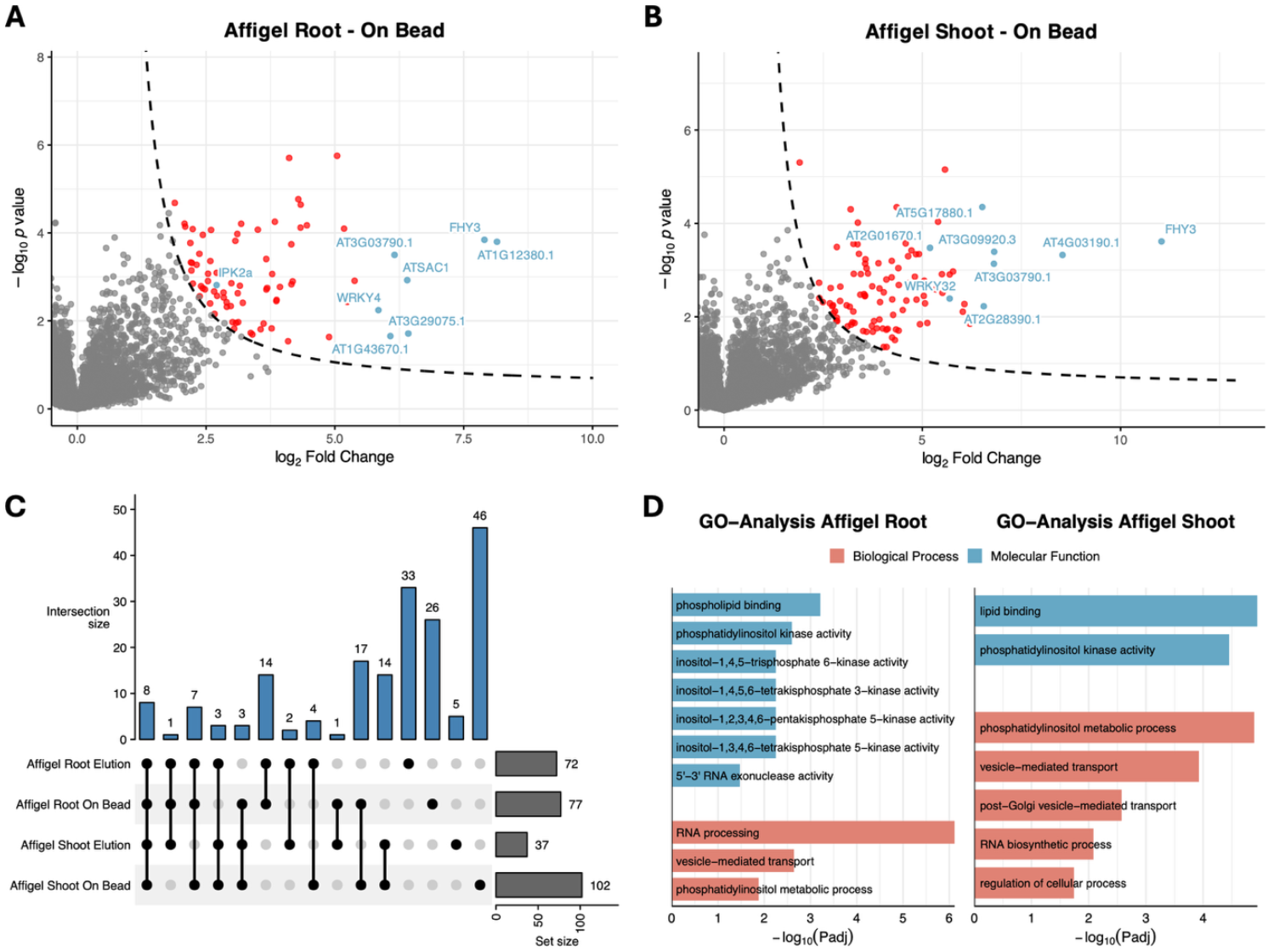
Analysis of Affi-Gel datasets from *Arabidopsis thaliana* samples. a,b) Volcano plots showing significantly enriched proteins (FDR < 0.05, S_0_ = 1) from Affi-Gel on-bead fractions of root (panel a) and shoot (panel b) samples. Highlighted proteins (blue) were selected based on high enrichment scores and/or their potential relevance to inositol phosphate signaling and phosphoinositide metabolism, including: a) AT5G07370.4 (IPK2α; inositol polyphosphate kinase 2 alpha), AT1G13960.2 (WRKY4; WRKY DNA-binding protein 4), AT1G22620.1 (ATSAC1; phosphoinositide phosphatase family protein), AT1G43670.1 (inositol monophosphatase family protein), and AT3G22170.2 (FHY3; far-red elongated hypocotyls 3). b) AT3G22170.2 (FHY3; far-red elongated hypocotyls 3), AT1G47550.2 (SEC3A; exocyst complex component), AT2G01670.1 (NUDT17; nudix hydrolase homolog 17), AT4G30935.1 (WRKY32; WRKY DNA-binding protein 32), and AT1G31440.1 (SH3 domain-containing protein). c) UpSet plot illustrating the overlap of enriched proteins across root and shoot samples. d) Gene Ontology (GO) analysis of enriched proteins from root and shoot samples, showing Driver Terms identified by g:Profiler. For this analysis, protein lists from on-bead and elution fractions were combined. Categories are grouped into molecular function (blue) and biological process (red).

Overlap analysis showed that on-bead fractions contained a broader range of interactors than eluates, with 17 proteins shared between root and shoot on-bead samples but only two in eluates. This indicates that on-bead fractions mainly enrich stronger or more stable binders, whereas eluates capture weaker or more transient associations. Several proteins were unique to individual conditions, with the shoot on-bead fraction containing the largest number of exclusive hits. GO-term analysis revealed significant enrichment of categories related to phosphatidylinositol metabolism and inositol phosphate signaling, consistent with the expected biological roles of 4/6-PP-InsP_5_ (**4**/**5**). Collectively, these results demonstrate that the Affi-Gel approach captures a selective set of high-affinity interactors with low background, providing a reliable platform for validation and future mechanistic studies.

Photoaffinity-based enrichment revealed a broader and more variable interactome than the Affi-Gel approach, reflecting the ability of covalent crosslinking to stabilize transient associations and retain weak binders that would otherwise be lost during washing. Photoaffinity probes may also label protein complexes associated with direct interactors, further expanding the apparent interactome. In roots, 180–380 proteins were enriched per condition (see Figure 4) and in shoots 90–490 (see Figure 5). Across all experiments, the custom-designed linker (compound **20**) consistently yielded more enriched proteins than the commercial Sulfo-SBED reagent (compound **21**). This difference likely reflects the inclusion of DTT in the custom linker experiments, which helped preserve protein integrity, whereas DTT was omitted with the commercial linker to avoid cleavage of its disulfide bridge. Additionally, linker design features such as crosslinking efficiency, spatial arrangement of reactive groups, and target accessibility may also have contributed to the broader interactome coverage observed with the custom linker. These probe-specific properties should be considered when interpreting the data and when planning future validation experiments.

**Figure 4:**
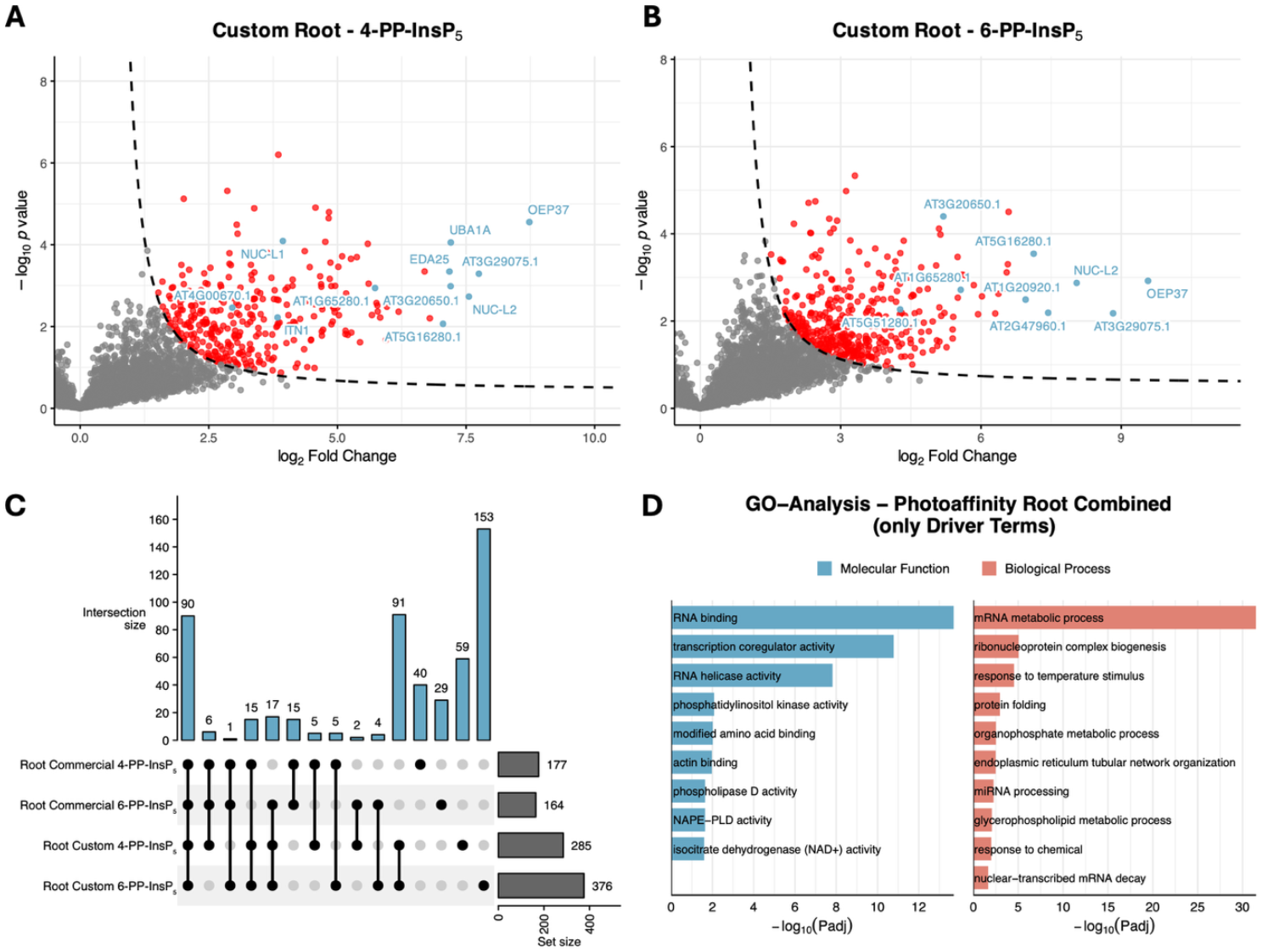
Analysis of photoaffinity-enriched datasets from *Arabidopsis thaliana* root samples. a,b) Volcano plots of significantly enriched proteins (FDR < 0.05, S0 = 1) from competition experiments with free 4-PP-InsP_5_ (panel A) or free 6-PP-InsP_5_ (panel B) as competitors. Highlighted proteins (blue) were selected based on high enrichment scores and/or their potential relevance to inositol phosphate signaling and phosphoinositide metabolism, including: a) AT4G00670.1 (Remorin family protein), AT1G48920.1 (NUC-L1; nucleolin like 1), AT3G12360.1 (ITN1; ankyrin repeat family protein), AT1G65280.1 (DNAJ heat shock N-terminal domain-containing protein), and AT3G20650.1 (mRNA capping enzyme family protein). b) AT5G51280.1 (DEAD-box protein, putative), AT3G20650.1 (mRNA capping enzyme family protein), AT1G65280.1 (DNAJ heat shock N-terminal domain-containing protein), AT1G20920.1 (P-loop containing nucleoside triphosphate hydrolases superfamily protein), and AT3G18610.1 (NUC-L2; nucleolin like 2). c) UpSet plot illustrating the overlap of enriched proteins across all root samples from photoaffinity enrichment experiments, including datasets from both custom and commercial linker experiments with 4-PP-InsP_5_ and 6-PP-InsP_5_ as competitors. d) Gene Ontology (GO) analysis of enriched proteins from root samples, showing Driver Terms identified by g:Profiler. For this analysis, all root sample datasets were combined. Categories are grouped into molecular function (blue) and biological process (red).

**Figure 5:**
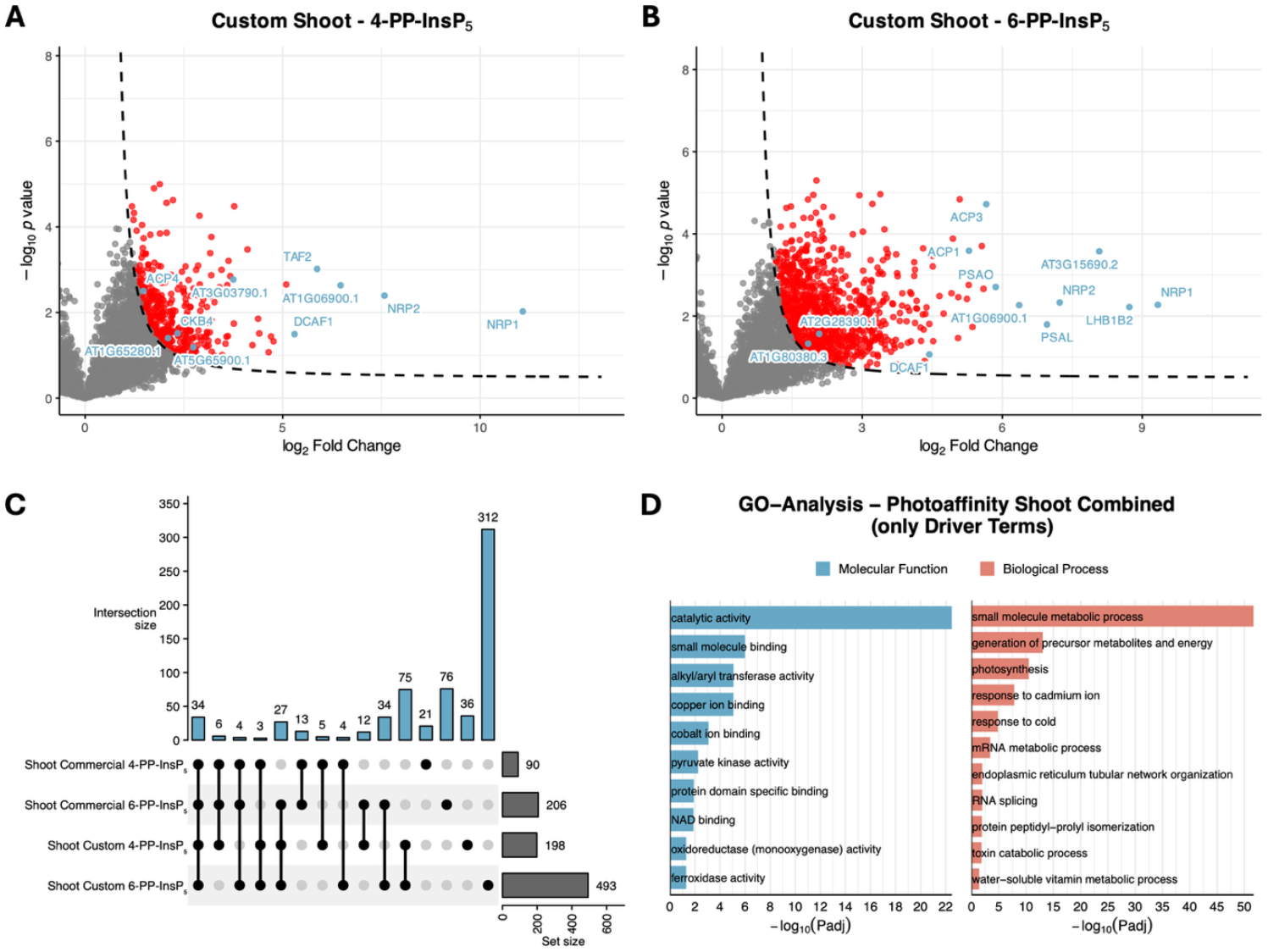
Analysis of photoaffinity-enriched datasets from *Arabidopsis thaliana* shoot samples. (a,b) Volcano plots of significantly enriched proteins (FDR < 0.05, S0 = 1) from competition experiments with free 4-PP-InsP_5_ (panel A) or free 6-PP-InsP_5_ (panel B) as competitors. Highlighted proteins (blue) were selected based on high enrichment scores and/or their potential relevance to inositol phosphate signaling and phosphoinositide metabolism, including: a) AT4G25050.1 (ACP4; acyl carrier protein 4), AT1G65280.1 (DNAJ heat shock N-terminal domain-containing protein), AT2G44680.2 (CKB4; casein kinase II beta subunit 4), AT3G03790.1 (ankyrin repeat/RCC1 family protein), and AT5G65900.1 (DEA(D/H)-box RNA helicase family protein). b) AT1G80380.3 (P-loop containing nucleoside triphosphate hydrolases superfamily protein), AT2G28390.1 (SAND family protein), AT4G31160.1 (DCAF1; DDB1-CUL4 associated factor 1), AT1G54630.1 (ACP3; acyl carrier protein 3), and AT3G05020.1 (ACP1; acyl carrier protein 1). c) UpSet plot illustrating the overlap of enriched proteins across all shoot samples from photoaffinity enrichment experiments, including datasets from both custom and commercial linker experiments with free 4-PP-InsP_5_ and 6-PP-InsP_5_ as competitors. d) Gene Ontology (GO) analysis of enriched proteins from shoot samples, showing Driver Terms identified by g:Profiler. For this analysis, protein lists from on-bead and elution fractions were combined. Categories are grouped into molecular function (blue) and biological process (red).

**Figure 6:**
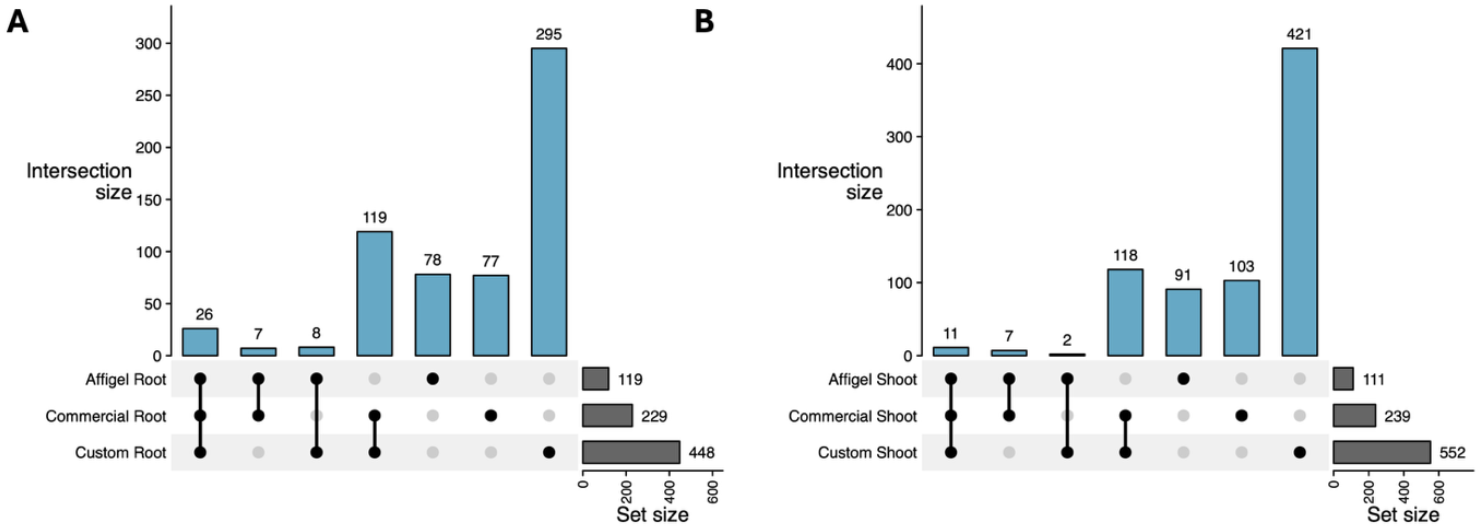
UpSet plots illustrating the overlap of enriched proteins across different enrichment strategies in *Arabidopsis thaliana* root (panel a) and shoot (panel b) samples. Data include Affi-Gel, commercial, and custom photoaffinity enrichment experiments. For Affi-Gel, on-bead and elution fractions were combined. For photoaffinity enrichment experiments, protein lists from both competition experiments were combined prior to analysis.

Competition experiments consistently showed that free 6-PP-InsP_5_ (**5**) displaced a broader set of proteins than 4-PP-InsP_5_ (**4**), suggesting a more diverse interactome. In root samples, the custom linker yielded 376 proteins displaced by 6-PP-InsP_5_ (**5**) compared to 285 by 4-PP-InsP_5_ (**4**), and in shoots the difference was even greater (493 vs. 198). With the commercial linker, the same trend was seen in shoots (206 vs. 90), whereas in roots 4-PP-InsP_5_ (**3**) displaced slightly more proteins than 6-PP-InsP_5_ (**4**) (177 vs. 164). These results support the often-contested view that PP-InsP_5_ isomers display distinct binding profiles, consistent with their having potentially different biological roles. Despite substantial overlap, each isomer also recruited unique subsets of interactors, indicating that 4-PP-InsP_5_ (**4**) and 6-PP-InsP_5_ (**5**) engage both with shared and specific binding partners under the tested conditions.

Notably, both root and shoot samples showed substantial but incomplete overlap of proteins across enrichment conditions, indicating that linker design, competition strategy, and tissue context shape the captured interactome. Unique subsets were detected in individual competition setups, with the largest proportion in custom linker 6-PP-InsP_5_ (**5**) experiments for both tissues. These findings underline that candidate interactors must be interpreted with caution and require thorough follow-up analysis. Importantly, proteins identified by affinity-based enrichment should not be assumed to always reflect physiologically relevant interactions. The loss of cellular compartmentalization during extraction may give rise to artefactual associations, and proteins that are part of multi-protein assemblies may co-purify without directly binding to the ligand.

GO-term enrichment, analyzed separately for root and shoot samples, revealed pronounced tissue-specific differences. In roots, categories linked to RNA processing and transcription, protein folding, and phosphatidylinositol metabolism were strongly enriched. Shoots instead showed enrichment of small-molecule metabolic processes, ion- and metabolite-binding activities, and photosynthesis-related terms, consistent with their distinct physiological roles. Notably, the term “response to temperature stimulus” was enriched in root samples, while “response to cold” appeared in shoots, suggesting that 4/6-PP-InsP_5_ (**4/5**) may contribute to temperature-related stress responses. This observation complements recent findings implicating 4/6-PP-InsP_5_ (**4/5**) in thermal signaling pathways in plants.^[12]^ Collectively, these data indicate 4/6-PP-InsP_5_ (**4/5**) may modulate diverse biological pathways, requiring detailed functional validation in follow-up studies.

These results further demonstrate the usefulness of photoaffinity-based approaches to reveal a broad spectrum of potential 4/6-PP-InsP_5_ interactors, while showing that probe design, competition strategy, and tissue context critically shape the captured interactome. Although the custom linker provided higher sensitivity, it may also increase non-specific or transient binding, underscoring the need for cautious interpretation and validation of candidate proteins. Together, these findings offer a refined perspective on the application of photoaffinity strategies to dissect complex plant interactomes of inositol pyrophosphates, potentially with information about isomer specific responses based on the competition isomer used.

Together, the Affi-Gel and photoaffinity enrichment strategies provided complementary perspectives on the Arabidopsis 4/6-PP-InsP_5_ interactome. The Affi-Gel method reproducibly captured a focused, tissue-independent subset of high-affinity interactors, while photoaffinity enrichment expanded coverage to include weaker and transient interactions stabilized through covalent crosslinking. Both approaches consistently enriched proteins associated with phosphatidylinositol metabolism and inositol phosphate signaling. Across datasets, more proteins were displaced by 6-PP-InsP_5_ (**5**) than by 4-PP-InsP5 (**4**), suggesting a broader interactome and potentially a more pronounced signaling role for this isomer. This observation aligns with prior findings that implicate 6-PP-InsP_5_ (**5**) as the biologically relevant isomer in *Dictyostelium discoideum*.^[37]^ Yet, of course the situation in plants might be different and future studies will have to provide clarity about the enantiomer identity of these PP-InsPs.

Several proteins with established inositol phosphate- or lipid-binding domains, such as PH, SPX, or C2, were reproducibly detected across datasets – including AT4G14740.2 (PH-like domain), AT3G59660.1 (C2 domain), and AT1G35350.1 (EXS family; SPX-related) – further supporting the specificity and reliability of the enrichment approaches, as these domains have previously been implied in PP-InsP binding.^[3,6]^

To evaluate whether the enrichment approach yields proteins capable of direct ligand interaction, we selected candidate interactors for biophysical validation based on favorable expression characteristics. Specifically, we prioritized small, predicted soluble proteins and in particular abundant domains lacking predicted transmembrane regions to facilitate recombinant production and downstream analysis. Among the considered candidates, AtFHA2 (AT3G07220.1), a FHA domain-containing protein, was the first to yield sufficient amounts of properly folded protein and was therefore selected for follow-up experiments. FHAs are domains present in at least 18 genes encoded by the Arabidopsis genome and they also occur in other eukaryotes and eubacteria.^[38]^ In our screen, it was identified in the root photoaffinity pulldown using the commercial linker and 4-PP-InsP_5_ (**4**) as competitor. Given its phosphothreonine-binding FHA domain,^[39]^ AtFHA2 might be involved in phosphorylation-dependent signaling processes that intersect with inositol pyrophosphate pathways.

Ligand binding was confirmed by isothermal titration calorimetry (ITC), which demonstrated direct interaction of AtFHA2 with InsP_6_ (**6**) and 4-PP-InsP_5_ (**4**) (see Figure 7A and 7B). While InsP_6_ (**6**) showed only weak binding with a dissociation constant of ca. 34 µM, 4-PP-InsP_5_ (**4**) bound substantially more tightly, with a K_d_ of ca. 4 µM. These results confirm a markedly higher binding affinity of 4-PP-InsP_5_ (**4**) under the tested conditions. These two ligands were selected based on their relevance to the experimental design: InsP_6_ (**6**) served as a broadly established reference, while 4-PP-InsP_5_ (**4**) corresponded to the competition ligand used during the enrichment. A broader comparison with additional PP-InsP_5_ isomers was not pursued, as protein yield was limited and 4-PP-InsP_5_ (**4**) was directly linked to the dataset in which AtFHA2 was identified. Circular dichroism (CD) spectroscopy further confirmed a well-folded secondary structure and revealed minor conformational changes upon ligand addition in a dose-dependent manner (see Figure 7C). Taken together, the direct *in vitro* interaction between AtFHA2 and 4-PP-InsP_5_ (**4**) exemplifies the ability of the enrichment strategy to uncover relevant binding partners for further *in vivo* validation.

**Figure 7:**
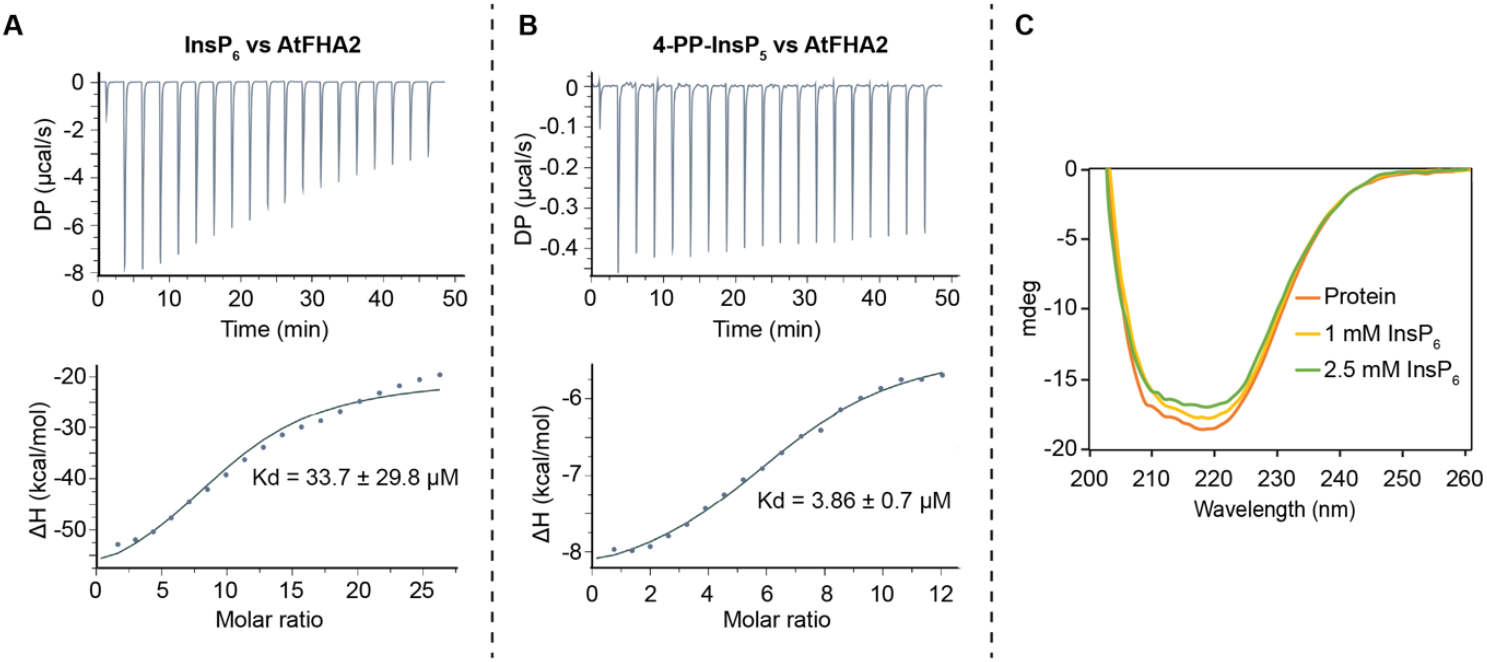
Biophysical validation of AtFHA2 as a ligand-binding protein. A) ITC analysis of InsP_6_ (**6**) binding to recombinant AtFHA2: raw titration data and corresponding binding isotherm. B) ITC analysis of 4-PP-InsP_5_ (**4**) binding to AtFHA2, displayed as raw data and fitted isotherm. C) Circular dichroism (CD) spectra of AtFHA2 in the absence and presence of InsP_6_, confirming a folded structure and no major conformational changes upon ligand binding.

## 3 Conclusion

In this study, we established complementary Affi-Gel- and photoaffinity-based enrichment strategies to systematically map the protein interactome of 4/6-PP-InsP_5_ (**4/5**) in *Arabidopsis thaliana*. Affi-Gel preferentially enriched a consistent core of high-affinity interactors, whereas photoaffinity labeling, applied for the first time to inositol pyrophosphate pull-downs, revealed a broader, context-dependent spectrum of proteins, reflecting both stable and transient interactions. The custom linker (compound **20**) outperformed the commercial reagent (compound **21**), demonstrating higher capture efficiency. Competition experiments further showed that 6-PP-InsP_5_ (**5**) displaced a more diverse set of proteins than 4-PP-InsP_5_ (**4**), suggesting isomer-specific roles in plant signaling. Functional annotation highlighted strong links to phosphatidylinositol metabolism and inositol phosphate signaling, as well as temperature stress related proteins. Recombinant validation of AtFHA2 provided direct evidence for specific ligand binding. Since multiple FHA domain-containing proteins exist in Arabidopsis and other organisms, it might be established as another general PP-InsP binding domain, much like SPX, PH, and C2. Together, these findings establish a versatile approach for PP-InsP interactome mapping in plants and provide a basis for future studies to clarify the cellular roles and signaling functions of this underexplored isomer.

## Supporting information

Supporting Info data and procedures

## Supporting Information

The authors have cited additional references within the Supporting Information.^[11,14,16,24,27,31,32,34,35,40–43]^

## Acknowledgement

We thank Dr. Stefan Braukmüller and Dr. Manfred Keller from MagRes at the University of Freiburg for their support and for providing a significant amount of NMR measurement time. We also thank Christoph Warth for HRMS measurements and Guizhen Lui, Mengsi Lu and Isabel Prucker for CE-MS measurements. We also thank Anne Harzen for performing in proteomics sample preparation and Brigitte Ueberbach for technical assistance with Arabidopsis cultivation and plant lysate preparation. This study was supported by the Deutsche Forschungsgemeinschaft (DFG) (Project ID 560443421, JE 572/11-1, to H.J.J.; SCHA 1274/5-1, to G.S.) and under Germany’s Excellence Strategy (CIBSS, EXC-2189, Project ID 390939984, to H.J.J.; PhenoRob, EXC-2070-390732324, to G.S.). H.J.J. acknowledges funding from the Volkswagen Foundation (VW Momentum Grant 98604). D.L. is grateful to the funding from Ministry of Education, MoE-STARS/STARS-2/2023-0162. ChatGPT (OpenAI, GPT-4o) was used for language refinement and support in generating R scripts for data analysis and visualization. The Table of Contents Figure and Figure 2 were created using BioRender.com.

