## Supporting Info data and procedures for "Affinity-Based Interactome Mapping of Inositol Pyrophosphates Reveals 4/6-PP-InsP_5_–Binding Proteins in Plants"

### Table of Contents


| | | |
|---|---|---|
| 1 | General Remarks | 2 |
| 2 | Chemical Synthesis | 4 |
| 3 | Affinity Enrichment Experiments | 19 |
| 4 | Proteomics | 21 |
| 5 | Biophysical Validation of AtFHA2–Ligand Interactions | 24 |
| 6 | References | 25 |
| 7 | Gene Ontology Term Enrichment Analysis by g:Profiler | 26 |
| 8 | NMR – Spectra | 34 |
| 9 | Mass Spectra, CE Electropherograms & HPLC | 68 |

### 1 General Remarks

**Reactions** were carried out using oven-dried glassware under an atmosphere of dry Argon and magnetically stirred, unless noted otherwise. Air- and moisture-sensitive liquids and solutions were transferred via syringe or stainless steel canula.

**Reagents** were purchased from commercial suppliers (Acros, Sigma-Aldrich, Fluka, TCI, BLDpharm, ChemPur, Alfa Aesar, VWR) and used without further purification, unless noted otherwise.

**Solvents** were obtained in analytical grade and used as received for extractions, precipitation and solid washings.

**Dry solvents** for reactions were purchased in a dry form from commercial suppliers (Sigma-Aldrich, Acros, Thermo Scientific) and stored over molecular sieves as well as under the atmosphere of dry Argon.

**Deuterated solvents** for NMR and reactions were obtained from commercial suppliers (Eurisotope and Deutero) in the indicated purity grade and used as received for NMR spectroscopy.

**Thin layer chromatography (TLC)** was performed with Merck silica gel 60 $F_{254}$ . Compounds were visually analysed by UV light (λ = 254 and 365 nm) or stained. Staining solutions: $KMnO_4$ stain (1.5 g $KMnO_4$, 10 g $Na_2CO_3$, 1.25 mL 10% aq. NaOH, 200 mL $H_2O$), phosphomolybdic acid stain (PMA, 3-4 g $H_3PMo_{12}O_{40}$ in 200 mL EtOH).

**Silica column chromatography** was carried out using silica gel 60 (0.04 – 0.063 mm, 230 – 400 mesh) from Macherery-Nagel as stationary phase and with non-dry solvents.

**Preparative RP-MPLC** was performed using a PuriFlash 5.125 by Advion-Interchim. The stationary phase consists of $C_{18}$ reversed-phase (15 µm or 30 µm) silica, prepacked in different column sizes and supplied by Advion-Interchim.

**Lyophilizations** were done with Christ Freeze Dryer Alpha 1-4 LDplus and Christ Freeze Dryer Alpha 1-2 LDplus.

**$^{1}$H-NMR** spectra were recorded on a Bruker 400 MHz (with Prodigy CryoProbe) spectrometer in the indicated deuterated solvent. Data are reported as follows: chemical shift (δ, ppm), multiplicity (s, singlet; d, doublet; t, triplet; q, quartet; m, multiplet; br. s, broad singlet), coupling constant(s) (J, Hz), integration. All signals were referenced to the internal solvent signal as standard ($D_2O$: δ = 4.79 ppm; MeCN-$d_3$: δ = 1.94 ppm, $CDCl_3$: δ = 7.26 ppm).

**$^{13}$C{$^{1}$H}-NMR** spectra were recorded with $^{1}$H-broadband decoupling on a Bruker 101 MHz (with Prodigy CryoProbe) spectrometers at 298K in the indicated deuterated solvent. All signals were referenced to an internal standard.

**$^{31}$P{$^{1}$H}-NMR** spectra and **$^{31}$P-NMR** spectra were recorded with $^{1}$H-broadband decoupling or $^{1}$H-coupling on a 162 MHz spectrometer (with Prodigy CryoProbe) in the indicated deuterated solvent. All signals were referenced to an internal standard (PPP).

**Mass spectra** were recorded by C. Warth (Mass spectrometry service of the University of Freiburg) on a Thermo LCQ Advantage [spray voltage: 2.5 – 4.0 kV, spray current: 5 µA, ion transfer tube: 250 (150) °C, evaporation temperature: 50 – 400°C].

### 2 Chemical Synthesis

#### 2.1 Synthesis of Phosphoramidites & Other Reagents

##### 2.1.1 Synthesis of Bn-Phosphoramidite **S1**

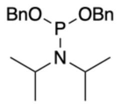

**S1**

$C_{20}H_{28}NO_2P$
MW: 345,42

The compound was synthesized according to Hofer et al. Analytical data were in accordance with literature.[1]

##### 2.1.2 Synthesis of PCP-Phosphoramidite **S2**

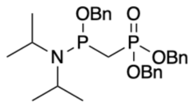

**S2**

$C_{28}H_{37}NO_4P_2$
MW: 513,5545

The compound was synthesized according to Hostachy et al. Analytical data were in accordance with literature.[2]

##### 2.1.3 Synthesis of PEG-Linker Alcohol **S3**

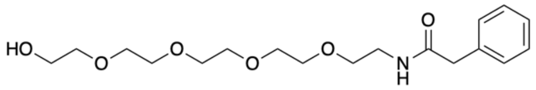

**S3**

$C_{18}H_{29}NO_6$
MW: 355,4310

The compound was synthesized according to Wu et al. Analytical data were in accordance with literature.[3]

#### 2.1.4 Synthesis of Fm-Phosphordiamidite S4

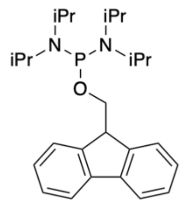

**S4**

$C_{26}H_{39}N_2OP$
MW: 426,5848

The compound was synthesized according to Ma et al. Analytical data were in accordance with literature.[4]

#### 2.1.5 Synthesis of PEG-Linker-Phosphates S5

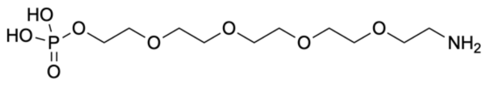

**S5**

$C_{10}H_{24}NO_8P$
MW: 317,2748

The compound was synthesized according to Wu et al. Analytical data were in accordance with literature.[3]

#### 2.1.6 Synthesis of 4-PP-InsP$_5$ (4) & 6-PP-InsP$_5$ (5)

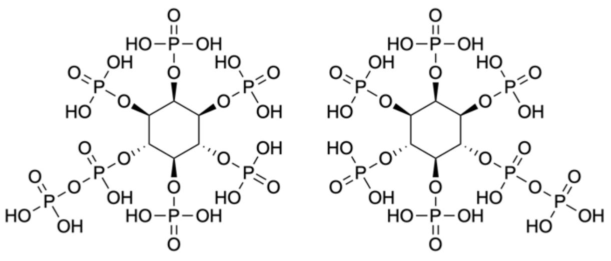

**4** **5**

$C_6H_{19}O_{27}P_7$
MW: 740,0073

The compounds were synthesized according to Qiu et al. Analytical data were in accordance with the literature.[5]

#### 2.2 Synthesis of Amino-PEG-4/6-PCP-InsP$_5$

##### 2.2.1 Synthesis of TBS-protected Ortho Ester **S6**

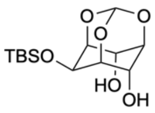


**S6**

$C_{13}H_{24}O_6Si$
MW: 304,4140

The compound was synthesized according to Baudin et al. Analytical data were in accordance with literature.[6]

##### 2.2.2 Synthesis of Diallyl-Protected Ortho Ester **8**

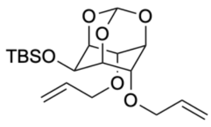


**8**

$C_{19}H_{32}O_6Si$
MW: 384,5440

TBS-protected ortho ester **S6** (1.00 g, 3.29 mmol) was dissolved in DMF (25 mL) and cooled to 0°C. Sodium hydride (342 mg, 8.54 mmol, 2.6 eq) was added portion-wise, and the mixture was stirred for 20 min at 0°C. Allyl bromide (738 µL, 1.03 g, 8.54 mmol, 2.6 eq) was added dropwise, along with a catalytic amount of sodium iodide (~50 mg). The reaction mixture was stirred overnight at room temperature, then diluted with EtOAc (200 mL). The solution was washed with water (100 mL) and brine (3 × 100 mL), and the layers were separated. The organic layer was dried over $MgSO_4$, and the solvent was removed under reduced pressure. The crude product was purified by silica column chromatography (cyclohexane/$Et_2O$ 5:1), and the product **8** (1.25 g, 3.24 mmol, 99%) was obtained as a colorless oil.

**$^1$H-NMR** (400 MHz, $CDCl_3$): $\delta$ = 5.98 – 5.82 (m, 2H), 5.57 – 5.52 (m, 1H), 5.31 (dq, *J* = 17.2, 1.6 Hz, 2H), 5.21 (dp, *J* = 10.5, 1.5 Hz, 2H), 4.40 (tt, *J* = 3.4, 1.6 Hz, 1H), 4.34 – 4.29 (m, 1H), 4.29 – 4.24 (m, 2H), 4.17 – 4.10 (m, 4H), 4.06 (ddt, *J* = 12.9, 5.4, 1.5 Hz, 2H), 0.97 (s, 9H), 0.17 (s, 6H) ppm.
**$^{13}$C-NMR** (101 MHz, $CDCl_3$): $\delta$ = 134.17, 117.02, 103.18, 74.08, 73.36, 70.45, 68.16, 61.71, 26.00, 18.46, -4.65 ppm.
**HRMS** (APCI) $[M+H]^+$ calculated for $C_{19}H_{33}O_6Si$: 385.2041, found 385.2043.

##### **2.2.3** Synthesis of Diallyl-protected Inositol **9**

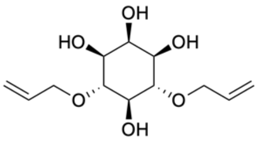

**9**

$C_{12}H_{20}O_6$
MW: 260,2860

Diallyl-protected ortho ester **8** (400 mg, 1.04 mmol) was dissolved in THF (10 mL), and aqueous HCl (1M, 10 mL) was added. The reaction mixture was stirred overnight at 40°C. After completion of the reaction cyclohexane (50 mL) and water (50 mL) were added, and the layers were separated. The organic layer was discarded, and the aqueous phase was concentrated under reduced pressure. The crude product was purified by silica column chromatography ($CH_2Cl_2$/MeOH 90:10), and the product **9** (197 mg, 0.757 mmol, 73%) was obtained as a colorless solid.

**$^1$H-NMR** (400 MHz, $CD_3CN$): δ = 5.98 (ddt, *J* = 17.4, 10.5, 5.7 Hz, 2H), 5.30 – 5.20 (m, 2H), 5.11 (ddt, *J* = 10.5, 2.1, 1.3 Hz, 2H), 4.33 – 4.19 (m, 4H), 3.85 (s, 1H), 3.40 – 3.22 (m, 6H), 3.19 (d, *J* = 3.4 Hz, 1H), 3.07 (d, *J* = 5.2 Hz, 2H) ppm.

**$^{13}$C-NMR** (101 MHz, $CD_3CN$): δ = 137.24, 116.44, 82.44, 75.43, 74.37, 73.71, 72.54 ppm.

**HRMS** (APCI) $[M-H]^-$ calculated for $C_{12}H_{19}O_6$: 259.1187, found 259.1182.

##### **2.2.4** Synthesis of Protected Inositol Tetraphosphate **10**

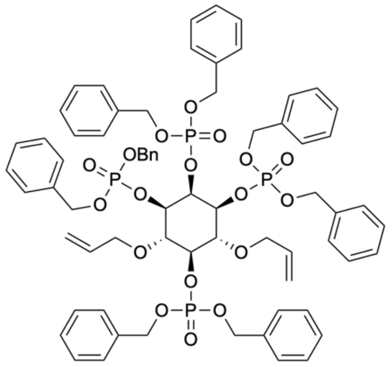

**10**

$C_{68}H_{72}O_{18}P_4$
MW: 1301,2010

Diallyl-protected Inositol **9** (179 mg, 688 µmol) was coevaporated twice with MeCN (3 mL) and dissolved in DMF (3 mL). Bn-phosphoramidite **S1** (1.85 g, 4.81 mmol, 7.0 eq) and DCI (948 mg, 4.81 mmol, 7.0 eq) were added, and the reaction mixture was stirred for 2 h. Reaction progress was monitored by $^{31}$P-NMR. After the phosphitylation was completed, oxidation was done by *m*CPBA (70%, 1.19 g, 4.81 mmol, 7.0 eq) at 0°C, and the reaction mixture was stirred for 10 min at room temperature. EtOAc (100 ml) and water (100 ml) were added, and the layers were separated. The organic layer was washed with water (100 ml) and brine (3 × 100 mL) and dried over $MgSO_4$. Celite was added, and the solvent was removed under reduced pressure to prepare the crude product as dry load for purification by automated reversed-phase MPLC (Interchim $C_{18}$-HP-Column, $H_2O$/MeCN, gradient: 20 - 100% MeCN). The product **10** (722 mg, 555 µmol, 81%) was obtained as a colorless oil.

**$^{1}$H-NMR** (400 MHz, $CDCl_3$): δ = 7.28 – 7.12 (m, 40H), 5.74 (ddt, *J* = 17.3, 10.4, 5.7 Hz, 2H), 5.27 (dt, *J* = 8.5, 2.4 Hz, 1H), 5.11 – 4.92 (m, 18H), 4.92 – 4.83 (m, 2H), 4.28 – 4.13 (m, 3H), 4.08 (dt, *J* = 5.8, 1.4 Hz, 4H), 3.71 (t, *J* = 9.6 Hz, 2H) ppm.

**$^{31}$P-NMR** (162 MHz, $CDCl_3$): δ = -1.38 (s, 2P), -1.59 (s, 1P), -1.93 (s, 1P) ppm.

**$^{13}$C-NMR** (101 MHz, $CDCl_3$): δ = 136.12 – 135.57 (m)*, 134.59, 128.67 – 127.51 (m)*, 117.16, 79.37 (d, *J* = 7.3 Hz), 77.21 (d, *J* = 5.0 Hz), 76.89 (dd, *J* = 5.6, 3.4 Hz), 75.34 (d, *J* = 7.0 Hz), 74.02, 70.57 – 68.37 (m)* ppm.

**Due to significant signal overlap, not all expected carbon resonances could be identified. Overlapping signals are listed as multiplets (m) and marked with an asterisk. Missing inositol signals are located below the $CDCl_3$ signal and were observed in HSQC.*

**HRMS** (ESI) $[M+Na]^+$ calculated for $C_{68}H_{72}O_{18}P_4Na$: 1323.3561, found 1323.3570.

#### **2.2.5** Synthesis of Protected Inositol Tetraphosphate **11**

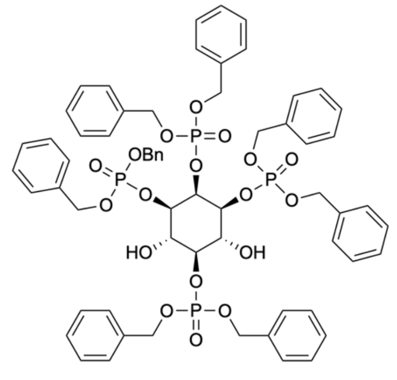


**11**

$C_{62}H_{64}O_{18}P_4$
MW: 1221,0710

Protected Inositol Tetraphosphate **10** (250 mg, 192 µmol) was dissolved in MeOH (10 mL) and $PdCl_2$ (75.7 mg, 384 mmol, 2.0 eq) was added. The reaction was stirred for 2h at room temperature. The reaction progress was monitored by $^{31}$P-NMR. After completion of the reaction the mixture was diluted with EtOAc (100 mL) and washed with a saturated aqueous $NaHCO_3$-solution (2 × 100 mL) and brine (100 mL). The organic phase was dried over $MgSO_4$ and the solvent was removed under reduced pressure. The product **11** (189 mg, 169 µmol, 88%) was obtained as a colorless solid was used without further purification.

**$^{1}$H-NMR** (400 MHz, $CDCl_3$): δ = 7.28 – 7.08 (m, 40H), 5.11 (dt, *J* = 8.9, 2.5 Hz, 1H), 5.07 – 4.89 (m, 16H), 4.39 – 4.32 (m, 4H), 4.25 (td, *J* = 9.0, 7.9 Hz, 1H), 3.91 (td, *J* = 9.4, 4.3 Hz, 2H) ppm.

**$^{31}$P-NMR** (162 MHz, $CDCl_3$): δ = -0.32 (s, 1P), -0.77 (s, 2P), -2.01 (s, 1P) ppm.

**$^{13}$C-NMR** (101 MHz, $CDCl_3$): δ = 135.99 – 135.36 (m)*, 128.75 – 127.60 (m)*, 81.62 (d, *J* = 5.9 Hz), 77.30 (d, *J* = 4.7 Hz), 75.94 (d, *J* = 4.6 Hz), 70.70 (t, *J* = 3.4 Hz), 70.20 – 69.22 (m)* ppm.

**Due to significant signal overlap, not all expected carbon resonances could be identified. Overlapping signals are listed as multiplets (m) and marked with an asterisk. Missing inositol signals are located below the $CDCl_3$ signal and were observed in HSQC.*

**HRMS** (ESI) $[M+NH_4]^+$ calculated for $C_{62}H_{68}NO_{18}P_4$: 1238.3381, found 1238.3385.

#### 2.2.6 Synthesis of Protected Inositol Pentaphosphate 12

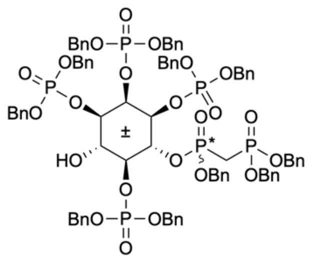


**12**

$C_{84}H_{86}O_{23}P_6$
MW: 1649,4316

Protected inositol tetraphosphate **11** (271 mg, 222 µmol) was coevaporated twice with MeCN (3 mL) and dissolved in $CH_2Cl_2$ (3 mL). PCP-phosphoramidite **S2** (137 mg, 266 mmol, 1.2 eq) and DCI (52.4 mg, 444 µmol, 2.0 eq) were added, and the reaction mixture was stirred for 1.5 h. Reaction progress was monitored by $^{31}$P-NMR. After the phosphitylation was completed, oxidation was done by *m*CPBA (70%, 109 mg, 444 mmol, 2.0 eq) at 0°C, and the reaction mixture was stirred for 10 min at room temperature. Celite was added, and the solvent was removed under reduced pressure to prepare the crude product as dry load for purification by automated reversed-phase MPLC (Interchim $C_{18}$-HP-Column, $H_2O$/MeCN, gradient: 20 - 100% MeCN). The product **12** (186 mg, 113 µmol, 51%) was obtained as a colorless oil.

**$^{1}$H-NMR** (400 MHz, $CDCl_3$): δ = 7.29 – 7.02 (m, 55H), 5.40 – 5.26 (m, 1H), 5.23 – 4.68 (m, 24H), 4.19 (qt, *J* = 9.3, 2.2 Hz, 1H), 4.10 – 4.00 (m, 1H), 3.98 – 3.84 (m, 2H), 2.89 – 2.45 (m, 2H) ppm.

**$^{31}$P-NMR** (162 MHz, $CDCl_3$): δ = 21.11 (d, 2P), 20.52 – 19.40 (m, 2P), 0.66 and 0.41 (s, 1P), -1.06 and -1.23 (s, 1P), -1.34 and -1.36 (s, 1P), -2.25 (s, 1P) ppm.

*The compound was obtained as a mixture of diastereomers arising from the formation of a stereogenic phosphorus center during phosphorylation. Duplication of some signals is observed in both the $^{1}H$ and $^{31}P$ NMR spectra, with summed integrals remaining consistent with the expected structure.*

**$^{13}$C-NMR** (101 MHz, $CDCl_3$): δ = 136.41 – 134.62 (m)*, 129.44 – 127.07 (m)*, 80.52 – 79.92 (m), 76.54 – 76.15 (m), 75.57 – 74.93 (m), 73.93 – 73.59 (m), 73.37 (d, *J* = 22.2 Hz), 70.97 – 69.22 (m)*, 68.43 – 67.33 (m)*, 27.82 – 24.76 (m) ppm.

**Due to significant signal overlap, not all expected carbon resonances could be identified. Overlapping signals are listed as multiplets (m) and marked with an asterisk. Missing inositol signal is located below the benzyl signals and was observed in HSQC.*

**HRMS** (ESI) $[M+Na]^+$ calculated for $C_{84}H_{86}O_{23}NaP_6$: 1671.3878, found 1671.3861.

#### **2.2.7** Synthesis of Protected Inositol Phosphate **13**

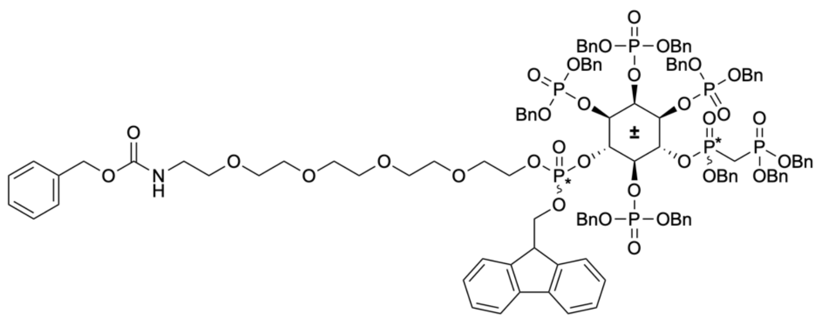

**13**

$C_{116}H_{124}NO_{32}P_7$
MW: 2261,0593

Fm-Phosphordiamidite **S4** (300 mg, 703 µmol, 6.2 eq) and PEG-Linker alcohol **S3** (261 mg, 703 µmol, 6.2 eq) were dissolved in MeCN (2 ml) and coevaporated twice with MeCN (2 ml). The mixture was dissolved in $CH_2Cl_2$ (1.3 mL) and a solution of ETT in $CH_2Cl_2$ (2.6 mL, 91.5 mg, 703 µmol, 6.2 eq) was added dropwise at 0°C. The reaction mixture was stirred for 30 min at room temperature. Reaction progress was monitored by $^{31}$P-NMR. The resulting phosphoramidite was directly used without purification to phosphorylate protected inositol pentaphosphate **12.**

Protected inositol pentaphosphate **12** (186 mg, 113 µmol) was coevaporated twice with MeCN (2 ml) and dissolved in $CH_2Cl_2$ (3 ml). The solution was added dropwise to the phosphoramidite solution, and a solution of ETT in MeCN (1 M, 338 µl, 44.0 mg, 338 µmol, 3.0 eq) was added dropwise at 0°C. Reaction progress was monitored by $^{31}$P-NMR. After the phosphitylation was completed, oxidation was done by *m*CPBA (70%, 83.4 mg, 338 µmol, 3.0 eq) at 0°C, and the reaction mixture was stirred for 10 min at room temperature. Celite was added, and the solvent was removed under reduced pressure to prepare the crude product as dry load for purification by automated reversed-phase MPLC (Interchim $C_{18}$-HP-Column, $H_2O$/MeCN, gradient: 20 - 100% MeCN). The product **13** (179 mg, 79.2 µmol, 70%) was obtained as a colorless oil.

*The compound was isolated as a mixture of four diastereomers in an undetermined ratio. Since the configuration at the stereocenters is removed in a subsequent deprotection step, the diastereomeric composition is not relevant for the final compound. Due to the overlap of signals arising from the diastereomeric mixture, the expected splitting patterns in the $^1H$, 13C, and $^{31}P$ NMR spectra were not fully resolved. Nevertheless, integration confirmed that the relative intensities of the overlapping $^1H$ and $^{31}P$ signals are consistent with the expected structure. The $^{13}C$ NMR spectrum is included for completeness but no peak assignments are provided due to significant signal overlap from the PEG and benzyl protecting groups as well as from the diastereomeric mixture.*

**$^1$H-NMR** (400 MHz, $CDCl_3$): δ = 7.77 – 7.02 (m, 68H), 5.70 – 5.48 (m, 1H), 5.36 – 4.79 (m, 27H), 4.59 – 4.34 (m, 1H), 4.34 – 3.86 (m, 7H), 3.62 – 3.01 (m, 18H), 3.00 – 2.60 (m, 2H) ppm.
**$^{31}$P-NMR** (162 MHz, $CDCl_3$): δ = 22.28 – 21.42 (1P, m), 20.63 – 19.95 (1P, m), -0.23 – -3.03 (5P, m) ppm.
**HRMS** (ESI) $[M+H]^+$ calculated for $C_{116}H_{125}O_{32}NP_7$: 2260.6343, found 2260.6401.

#### 2.2.8 Synthesis of Amino-PEG-4/6-PCP-InsP$_5$ (**14**)

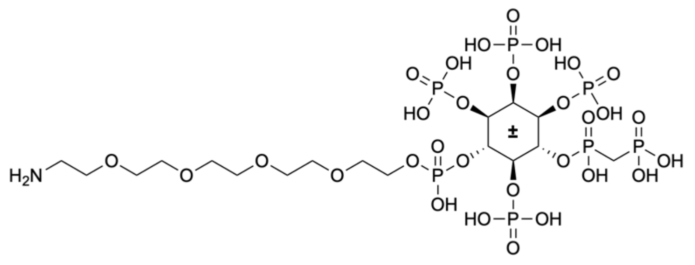


**14**

$C_{17}H_{42}NO_{30}P_7$
MW: 957,3163

Protected inositol phosphate **13** (20.0 mg, 8.85 µmol) was dissolved in a mixture of *t*BuOH and $H_2O$ (3.6 mL, 40:7). $NaHCO_3$ (28.2 mg, 26.5 µmol, 13.0 eq) and Pd/C (10% on activated charcoal, 28.2 mg, 26.5 µmol, 3.0 eq) were added and the reaction was hydrogenated for 21h at 30 bar (autoclave). The reaction was interrupted and water (4 mL) was added to the mixture to ensure dissolution. The reaction was further hydrogenated for an additional 3h at 30 bar (autoclave). The catalyst was removed by filtration through a syringe filter. The filter was rinsed with water (2 mL). The solvent was removed by lyophilization and the product **14** (8.00 mg, 7.20 µmol, 81%) was obtained as a colorless solid without any further purification.

*NMR samples contained diisopropylamine to improve resolution. Due to the low amount of compound, a direct $^{13}$C-NMR measurement was not practical; therefore, $^{13}$C-NMR resonances were assigned via the edHSQC spectrum. Two inositol protons overlapped with the water peak, as confirmed by edHSQC.*

**$^{1}$H-NMR** (400 MHz, $CDCl_3$): δ = 4.58 – 4.47 (m, 3H), 4.39 (d, *J* = 12.0 Hz, 1H), 4.25 – 4.15 (m, 2H), 3.83 (t, *J* = 4.5 Hz, 2H), 3.81 – 3.71 (m, 12H), 3.66 (t, *J* = 5.3 Hz, 2H), 2.94 (t, *J* = 5.3 Hz, 2H), 2.24 – 2.11 (m, 2H) ppm.

**$^{31}$P-NMR** (162 MHz, $CDCl_3$): δ = 22.04 (d, *J* = 5.9 Hz, 1P), 13.27 (d, *J* = 6.0 Hz, 1P), 5.20 (s, 1P), 4.43 (s, 1P), 4.25 (s, 1P), 3.85 (s, 1P), -0.94 (s, 1P) ppm.

**$^{13}$C-NMR** (101 MHz, $CDCl_3$): δ = 74.76, 73.79, 73.47, 70.89, 70.57, 70.57, 69.60, 69.60, 65.73, 65.41, 39.62, 30.60, 29.31, 28.02 ppm.

**HRMS** (ESI) $[M-2H]^{2-}$ calculated for $C_{17}H_{40}NO_{30}P_7$: 477.4904, found 477.4892.

#### 2.3 Synthesis of the Photoaffinity Linker

##### 2.3.1 Synthesis of Aminohexanoic methyl ester (S7)

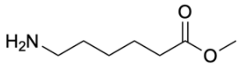

**S7**

$C_7H_{15}NO_2$
MW: 145,2020

The compound was synthesized according to Tantisuwanno et al. Analytical data were in accordance with literature.[7]

##### 2.3.2 Synthesis of Protected Lysine Derivative 16

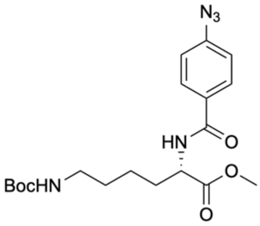

**16**

$C_{19}H_{27}N_5O_5$
MW: 405,4550

Boc-Lys-OMe · HCl (800 mg, 3.07 mmol) and 4-azidobenzoic acid (551 mg, 3.38 mmol, 1.1 eq) were dissolved in $CH_2Cl_2$ (25 ml). HOBt (498 mg, 3.69 mmol, 1.2 eq), EDCI · HCl (707 mg, 3.69 mmol, 1.2 eq) and triethyl amine (1.49 ml, 1.09 g, 10.8 mmol, 3.5 eq) were added, and the reaction mixture was stirred overnight at room temperature. The reaction mixture was dilluted with $CH_2Cl_2$ (25 ml), and washed with $KHSO_4$-solution (1 M, 2 × 50 mL) and saturated $NaHCO_3$-

solution (3 × 50 mL). The mixture was dried over $MgSO_4$ and the solvent was removed under reduced pressure. The crude product was purified by silica column chromatography (cyclohexane/EtOAc 1:1), and the product **16** (1.18 g, 2.90 mmol, 94%) was obtained as a beige solid.

**$^1$H-NMR** (400 MHz, $CDCl_3$): $\delta$ = 7.80 – 7.70 (m, 2H), 7.06 – 6.95 (m, 2H), 6.68 (d, *J* = 7.6 Hz, 1H), 4.71 (td, *J* = 7.6, 5.0 Hz, 1H), 4.53 (s, 1H), 3.71 (s, 3H), 3.04 (q, *J* = 6.5 Hz, 2H), 1.96 – 1.84 (m, 1H), 1.80 – 1.69 (m, 1H), 1.53 – 1.25 (m, 13H) ppm.
**$^{13}$C-NMR** (101 MHz, $CDCl_3$): $\delta$ = 173.07, 166.16, 156.13, 143.63, 130.34, 128.98, 119.02, 79.14, 52.53, 52.50, 40.02, 32.18, 29.69, 28.39, 22.50 ppm.
**HRMS** (ESI) $[M+Na]^+$ calculated for $C_{19}H_{27}N_5NaO_5$: 428.1904, found 428.1899.

##### 2.3.3 Synthesis of Protected Peptide Derivative 17

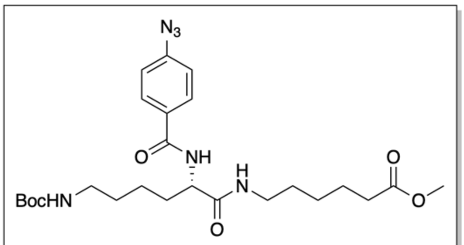


**17**

$C_{25}H_{38}N_6O_6$
MW: 518,6150

Protected lysine derivative **16** (400 mg, 987 mmol) was dissolved in MeOH (16 mL), and an aqueous solution of sodium hydroxide (1 M, 1.97 mL, 79.0 mg, 1.97 mmol, 2.0 eq) was added at 0°C. The reaction mixture was stirred for 30 min at 0°C. When the reaction was completed, $CHCl_3$ (100 mL) and water (50 mL) were added, and the pH was adjusted to 2-3 (HCl, 1 M). The layers were separated, and the aqueous layer was extracted with $CHCl_3$ (5 × 50 mL). The combined organic layers were dried over $MgSO_4$, and the solvent was removed under reduced pressure. The crude product was directly subjected to the following peptide coupling without further purification.
The crude product was dissolved in $CH_2Cl_2$ (8 mL) and aminohexanoic methyl ester (**S7**) (158 mg, 1.08 mmol, 1.1 eq), HOBt (160 mg, 1.18 mmol, 1.2 eq), EDCI · HCl (227 mg, 1.18 mmol, 1.2 eq) and triethylamine (478 µL, 349 mg, 3.45 mmol, 3.5 eq) were added. The reaction mixture was stirred overnight at room temperature. After completion of the reaction, the solvent was removed under reduced pressure. The residue was dissolved in EtOAc (50 mL), and the mixture was washed with an aqueous $KHSO_4$-solution (1 M, 2 × 50 mL), saturated aqueous $NaHCO_3$-solution (3 × 50 mL) and brine (50 mL). The mixture was dried over $Na_2SO_4$, and the solvent was removed under reduced pressure. The crude product was purified by silica column chromatography

(cyclohexane/EtOAc 1:2 to 1:4), and the product **17** (342 mg, 659 µmol, 67%) was obtained as a beige solid.

**$^1$H-NMR** (400 MHz, $CDCl_3$): δ = 7.81 – 7.70 (m, 2H), 7.05 (d, *J* = 7.8 Hz, 1H), 7.02 – 6.93 (m, 2H), 6.65 (q, *J* = 6.2 Hz, 1H), 4.67 (t, *J* = 6.0 Hz, 1H), 4.55 (td, *J* = 7.8, 5.8 Hz, 1H), 3.59 (s, 3H), 3.27 – 3.09 (m, 2H), 3.03 (dhept, *J* = 13.5, 6.8 Hz, 2H), 2.27 – 2.16 (m, 2H), 1.88 (dtd, *J* = 13.5, 7.7, 5.7 Hz, 1H), 1.80 – 1.64 (m, 1H), 1.61 – 1.16 (m, 19H) ppm.
**$^{13}$C-NMR** (101 MHz, $CDCl_3$): δ = 174.01, 171.65, 166.41, 156.17, 143.65, 130.26, 128.99, 118.98, 79.10, 53.47, 51.52, 40.04, 39.34, 33.79, 32.38, 29.66, 29.04, 28.41, 26.30, 24.38, 22.73 ppm.
**HRMS** (ESI) $[M-H]^-$ calculated for $C_{25}H_{37}N_6O_6$: 517.2780, found 517.2776.

#### **2.3.4** Synthesis of Desthiobiotin Derivative **18**

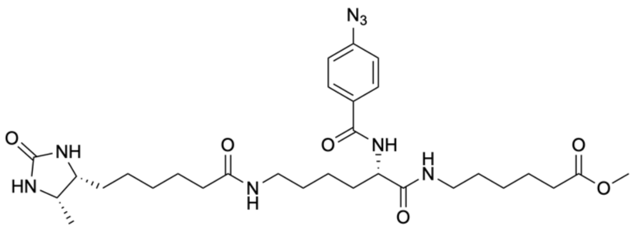


**18**
$C_{30}H_{46}N_8O_6$
MW: 614,7480

Protected peptide derivative **17** (227 mg, 438 µmol) was dissolved in $CH_2Cl_2$ (22 mL), and trifluoroacetic acid (11 mL) was added. The reaction mixture was stirred for 90 min at room temperature. When the reaction was completed, toluene (15 mL) was added. The solvent was removed under reduced pressure and the residue was coevaporated with toluene (10 mL). The crude product was used directly in the subsequent peptide coupling without further purification. The crude product was dissolved in DMF (11 mL) and desthiobiotin (103 mg, 481 µmol, 1.1 eq), HOBt (88.7 mg, 657 µmol, 1.5 eq), EDCI · HCl (126 mg, 657 µmol, 1.5 eq) and triethylamine (182 µL, 133 mg, 1.31 µmol, 3.0 eq) were added. The reaction mixture was stirred overnight at room temperature. After completion of the reaction, the reaction mixture was diluted with $CH_2Cl_2$ (6 mL) and washed with an aqueous $KHSO_4$ solution (1 M, 2 × 10 mL), saturated aqueous $NaHCO_3$ solution (3 × 10 mL) and brine (2 × 10 mL). The organic layer was dried over $Na_2SO_4$, and the solvent was removed under reduced pressure. The crude product was purified by silica column chromatography ($CH_2Cl_2$/MeOH 95:5 to 9:1), and the product **18** (182 mg, 296 µmol, 68%) was obtained as a beige solid.

**$^1$H-NMR** (400 MHz, $CDCl_3$): δ = 7.94 – 7.86 (m, 2H), 7.73 (d, *J* = 8.1 Hz, 1H), 7.26 (t, *J* = 5.7 Hz, 1H), 7.12 – 7.04 (m, 2H), 6.48 (t, *J* = 5.4 Hz, 1H), 6.02 (s, 1H), 4.79 – 4.66 (m, 2H), 3.90 – 3.74 (m, 2H),

3.68 (s, 3H), 3.35 – 3.17 (m, 4H), 2.31 (t, *J* = 7.4 Hz, 2H), 2.28 – 2.11 (m, 2H), 1.96 – 1.71 (m, 3H), 1.70 – 1.27 (m, 17H), 1.13 (d, *J* = 6.1 Hz, 3H) ppm.

**$^{13}$C-NMR** (101 MHz, $CDCl_3$): δ = 174.14, 173.31, 172.24, 166.87, 163.59, 143.67, 130.09, 129.20, 118.96, 55.10, 53.04, 51.53*, 39.30, 38.44, 35.57, 33.88, 31.61, 29.24, 28.97, 28.13, 27.35, 26.34, 25.25, 24.74, 24.49, 22.52, 15.82 ppm.

** The HSQC spectrum reveals that this signal corresponds to two overlapping $^{13}$C resonances.*

**HRMS** (ESI) $[M+Na]^+$ calculated for $C_{30}H_{46}N_8NaO_6$: 637.3433, found 637.3440.

#### 2.3.5 Synthesis of Sulfo-NHS-Ester 19

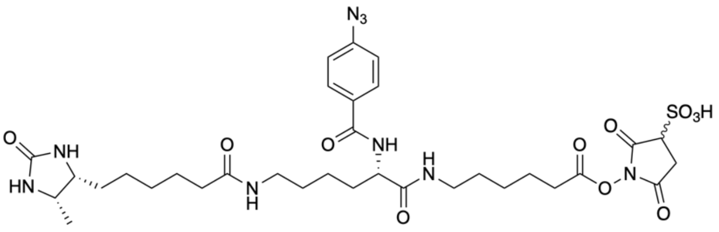


**19**

$C_{33}H_{47}N_9O_{11}S$
MW: 777,8510

Desthiobiotin derivative **18** (50.0 mg, 81.3 µmol) was dissolved in MeOH (1.6 mL) and an aqueous solution of NaOH (1M, 5 mL) was added at 0°C. The reaction mixture was stirred for 90 min at 0°C. When Reaction was completed $CHCl_3$ (30 mL) and water (30 mL) were added, and the pH was adjusted to 2-3 (HCl, 1M). The layers were separated, and the aqueous layer was extracted with $CHCl_3$ (5 × 30 mL). The combined organic layers were dried over $MgSO_4$, and the solvent was removed under reduced pressure. The crude product was used directly in the subsequent reaction without further purification.

The crude product (35.0 mg, 58.3 µmol) was dissolved in DMF (1.6 mL), and *N*-hydroxysulfosuccinimide sodium salt (12.7 mg, 58.3 µmol, 1.0 eq) and DCC (36.1 mg, 175 µmol, 3.0 eq) were added. The reaction mixture was stirred for 48 h at room temperature. Reaction progress was monitored by HPLC. After completion of the reaction the solution was cooled to 4°C and the resulting solid was filtered off. A mixture of EtOAc and cyclohexane (1:1, 30 mL) was added to precipitate the product **19** (49.0 mg, 61.3 µmol, 75%) as a colorless solid.

**$^{1}$H-NMR** (400 MHz, DMSO-$d_6$): δ = 8.35 (d, *J* = 7.9 Hz, 1H), 8.02 – 7.85 (m, 3H), 7.77 – 7.64 (m, 1H), 7.29 – 7.14 (m, 2H), 6.27 (s, 1H), 6.10 (s, 1H), 4.39 – 4.29 (m, 1H), 3.69 (dd, *J* = 8.8, 2.4 Hz, 1H), 3.64 – 3.55 (m, 1H), 3.46 (td, *J* = 8.0, 5.7 Hz, 1H), 3.12 – 2.92 (m, 4H), 2.92 – 2.80 (m, 1H), 2.75 – 2.58 (m, 1H), 2.17 (t, *J* = 7.4 Hz, 1H), 2.01 (t, *J* = 7.4 Hz, 2H), 1.75 – 1.58 (m, 4H), 1.57 – 1.12 (m, 18H), 0.95 (d, *J* = 6.4 Hz, 3H).

**$^{13}$C-NMR** (101 MHz, DMSO-$d_6$): $\delta$ = 174.42, 171.88, 171.58, 171.33, 168.79, 167.99, 165.33, 165.30, 162.77, 142.27, 130.74, 129.46, 118.71, 56.27, 56.18, 54.94, 53.46, 50.19, 38.33, 38.15, 35.35, 33.65, 31.36, 30.92, 30.72, 30.11, 29.47, 28.87, 28.75, 28.71, 28.46, 25.87, 25.52, 25.30, 25.19, 24.17, 23.87, 15.46 ppm.

*Additional $^{13}$C NMR signals arise from the formation of diastereomeric sulfo-NHS esters. The use of racemic sulfo-NHS introduces a stereocenter, resulting in two species with distinct chemical environments.*

**HRMS** (ESI) [M-H]$^-$ calculated for $C_{33}H_{46}N_9O_{11}S$: 776.3043, found 776.3048.

#### 2.4 Synthesis of the Photoaffinity Reagents

##### 2.4.1 Synthesis of Photoaffinity Reagent 20 (Custom Linker)

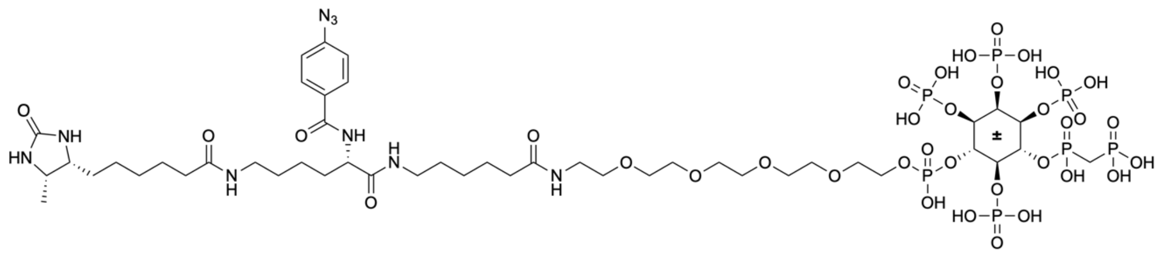

**20**

$C_{46}H_{84}N_9O_{35}P_7$
MW: 1540,0223

Amino-PEG-4/6-PCP-InsP$_5$ (**14**) (5.00 mg, 4.50 µmol) was dissolved in a $NaHCO_3$ buffer (0.2 M, 500 µL). A solution of Sulfo-NHS-Ester **19** (10 mg/mL, 1.0 mL, 10.0 mg, 12.5 µmol, 2.8 eq) was added and the reaction mixture was stirred for 2h at room temperature. The mixture was directly subjected to purification by automated reversed-phase MPLC (Interchim $C_{18}$-HP-Column, $H_2O$/MeCN, gradient: 20 - 90% MeCN, 10% TEAA buffer). The product **20** was obtained as a colorless oil (7.00 mg, 3.11 µmol, 69%) in form of the triethylammonium salt.

*Due to extensive signal overlap with solvent (MeOD-$d_4$, $H_2O$), triethylammonium counterions, and the PEG linker region, only selected resonances could be confidently assigned. Characteristic signals, such as aromatic protons, the desthiobiotin methyl group, and triethylammonium peaks, were observed as expected. Resonances associated with the PEG linker and the inositol core overlap with solvent and counterion signals and could not be individually resolved. Full spectra are provided. Compound identity was confirmed by $^{31}$P NMR, HRMS, and analytical HPLC. The inositol $^{13}$C signals are only visible in HSQC spectra, as $^{31}$P – $^{13}$C coupling causes broadening in direct $^{13}$C NMR. $^{13}$C signals marked with m indicate overlapping resonances.*

**$^1$H-NMR** (400 MHz, MeOD-$d_4$): $\delta$ = 7.98 – 7.90 (m, 1H), 7.21 – 7.13 (m, 1H), 5.34 (ddd, $J$ = 5.6, 4.4, 1.1 Hz, 0H), 5.16 (d, $J$ = 10.2 Hz, 0H), 4.71 – 4.62 (m, 0H), 4.57 (q, $J$ = 9.6 Hz, 0H), 4.50 – 4.41 (m, 0H), 4.35 – 4.09 (m, 2H), 3.87 – 3.75 (m, 0H), 3.75 – 3.46 (m, 8H), 2.53 – 2.10 (m, 5H), 2.09 – 2.00 (m, 1H), 1.10 (d, $J$ = 6.5 Hz, 2H) ppm.

**$^{31}$P-NMR** (162 MHz, MeOD-$d_4$): δ = 19.81 (d, 1P, $J$ = 11.2 Hz), 13.99 (d, 1P, $J$ = 11.3 Hz), 0.87 (s, 2P), 0.76 (s, 1P) , 0.36 (s, 1P), 0.06 (s, 1P) ppm.

**$^{13}$C-NMR** (101 MHz, MeOD-$d_4$): δ = 177.90, 174.86, 174.75, 173.01, 167.62, 164.77, 143.65, 130.38, 129.19, 118.59, 70.22 – 68.90 (m), 55.98, 54.13, 51.29, 38.93, 38.74, 38.58, 35.61, 35.51, 35.13, 34.80, 31.65, 31.48, 29.66 – 28.48 (m), 26.72, 26.14, 25.72, 25.56 – 25.41 (m), 25.24, 24.60, 23.10, 22.32, 14.24, 13.02, 7.85 ppm.

**HRMS** (ESI) $[M-2H]^{2-}$ calculated for $C_{46}H_{82}N_9O_{35}P_7$: 768.6544, found 768.6544.

#### **2.4.2** Synthesis of Photoaffinity Reagent **21** (Commercial Linker)

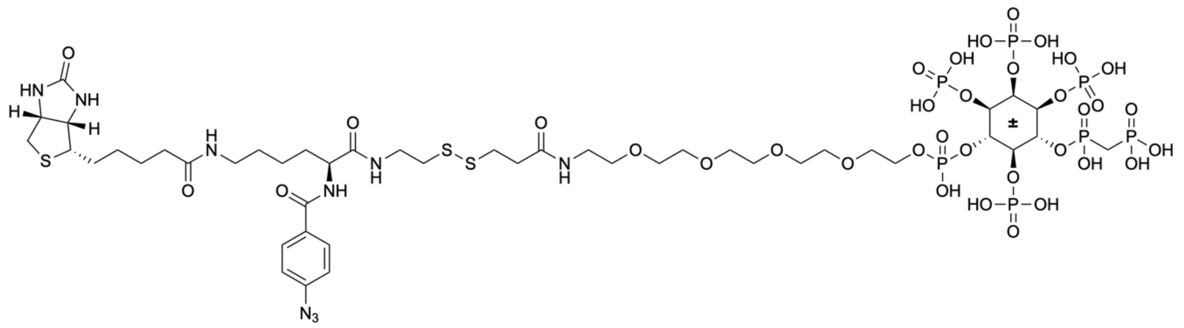

**21**

$C_{45}H_{80}N_9O_{35}P_7S_3$
MW: 1620,1593

Amino-PEG-4/6-PCP-InsP$_5$ (**14**) (4.00 mg, 3.60 µmol) was dissolved in a $NaHCO_3$ buffer (0.4 M, 400 µL). A solution of commercial Sulfo-SBED linker (10 mg/mL, 500 µL, 5.00 mg, 5.68 µmol, 1.6 eq) was added and the reaction mixture was stirred for 30 min at room temperature. The mixture was directly subjected to purification by automated reversed-phase MPLC (Interchim $C_{18}$-HP-Column, $H_2O$/MeCN, gradient: 20 - 90% MeCN, 10% TEAA buffer). The product **21** was obtained as a colorless oil (5.00 mg, 2.58 µmol, 72%) in form of the triethylammonium salt.

*Due to extensive signal overlap with solvent (MeOD-d$_4$, $H_2O$), triethylammonium counterions, and the PEG linker region, only selected resonances could be confidently assigned. Characteristic signals, such as aromatic protons and triethylammonium peaks, were observed as expected. Resonances associated with the PEG linker and the inositol core overlap with solvent and counterion signals and could not be individually resolved. Full spectra are provided. Compound identity was confirmed by $^{31}$P NMR, HRMS, and analytical HPLC. The inositol $^{13}$C signals are only visible in HSQC spectra, as $^{31}$P – $^{13}$C coupling causes broadening in direct $^{13}$C NMR. $^{13}$C signals marked with m indicate overlapping resonances.*

**$^{1}$H-NMR** (400 MHz, MeOD-$d_4$): δ = 8.00 – 7.90 (m, 1H), 7.20 – 7.14 (m, 1H), 5.19 (d, $J$ = 11.0 Hz, 1H), 4.72 – 4.56 (m, 1H), 4.53 – 4.44 (m, 1H), 4.39 – 4.18 (m, 3H), 3.77 – 3.42 (m, 9H), 3.31 (dt, $J$ = 3.3, 1.6 Hz, 19H), 3.00 – 2.83 (m, 2H), 2.78 – 2.67 (m, 1H), 2.67 – 2.60 (m, 1H), 2.45 – 2.30 (m, 1H), 2.18 (t, $J$ = 7.2 Hz, 1H), 1.99 – 1.78 (m, 1H), 1.78 – 1.36 (m, 4H) ppm.

**$^{31}$P-NMR** (162 MHz, MeOD-$d_4$): $\delta$ = 20.18 (s, 1P), 13.37 (s, 1P), 1.13 (s, 1P), 0.94 (s, 1P), 0.59 (s, 1P), 0.05 (s, 1P), -0.40 (s, 1P) ppm.

**$^{13}$C-NMR** (101 MHz, MeOD-$d_4$): $\delta$ = 174.65, 173.33, 172.41, 167.71, 164.71, 143.67, 130.35, 129.22, 118.60, 70.21 – 69.76 (m), 69.10, 61.96, 60.24, 55.85 – 55.30 (m), 54.20, 39.67, 39.08, 38.61, 38.23, 37.03, 35.42, 35.21, 33.87, 31.36, 28.70, 28.05, 25.51, 25.48, 23.12 ppm.

**HRMS** (ESI) $[M-2H]^{2-}$ calculated for $C_{45}H_{78}N_9O_{35}P_7S_3$: 808.5968, found 808.5968.

### 3 Affinity Enrichment Experiments

#### 3.1 Buffer Compositions

Wash Buffer:

- 50 mM Tris pH 7.5
- 100 mM NaCl
- 10 % glycerol (v/v)
- 1 mM $MgCl_2$

Extraction Buffer:

- 50 mM Tris pH 7.5
- 100 mM NaCl
- 10 % Glycerol
- 1 mM $MgCl_2$
- 1 % Triton X-100 (v/v)
- 5 mM DTT (only Affi-Gel and photoaffinity reagent custom)
- 10 µL/mL protease inhibitor cocktail (Sigma #P9599)
- 20 µL/mL phosphatase inhibitor cocktail (Sigma #5726)
- 20 µL/mL phosphatase inhibitor cocktail (Sigma #P0044)

#### 3.2 Affi-Gel Based Affinity Enrichment Experiments

Plants were grown as described by Schneider et al.[8] Root and shoot tissues (~6 g) were harvested and pulverized in liquid nitrogen. The powdered material was extracted with 8–10 mL of the extraction buffer (composition provided in chapter 3.1). The suspension was incubated for 30 minutes on ice. Lysates were clarified by three consecutive centrifugation steps (15 minutes at 20.000 g, 4°C), transferring the supernatant to a new tube after each step. The final lysate was kept on ice. Protein concentrations were quantified using the Pierce™ 660 nm Protein Assay Reagent (Thermo Scientific) according to the manufacturer's protocol. Samples were diluted 1:10 in milliQ water prior to analysis. Average protein concentrations were ~0.5 mg/mL for root lysates and 1–2 mg/mL for shoot lysates.

Affinity reagents were prepared by coupling either Amino-PEG-4/6-PCP-$InsP_5$ (**12**) or PEG-Linker-Phosphates **S5** to Affi-Gel beads as described by Furkert et al.[9] and stored at 4°C until use. For pull-down experiments, Affi-Gel beads (200 µL) were washed twice with 1 mL of wash buffer (centrifuged for 3 minutes at 2000 g, supernatant discarded). The beads were then incubated with

1 mL of plant lysate at 4°C for 1 hour on an overhead shaker. After incubation, the beads were washed three times with 1 mL of wash buffer (each wash: 10 minutes on an overhead shaker, centrifuged for 3 minutes at 2000 g, supernatant discarded). Following the final wash, the beads were incubated with 100 µL of wash buffer supplemented with 20 mM $InsP_6$ for 1 hour at 4°C on an overhead shaker. Beads and supernatant were then separated, frozen in liquid nitrogen, and stored at -80°C.

Experiments were conducted with both the 4-/6-PP-$InsP_5$ affinity reagent and the phosphate control reagent under identical conditions.

#### 3.3 Photoaffinity Based Affinity Enrichment Experiments

Plants were grown as described by Schneider et al.[8] Root and shoot tissues (~6 g) were harvested and pulverized in liquid nitrogen. The powdered material was extracted with 8–10 mL of extraction buffer (composition provided in chapter 7.3.3.1) and incubated for 30 minutes on ice. Lysates were clarified by three consecutive centrifugation steps (15 minutes at 20,000 g, 4°C), transferring the supernatant to a new tube after each step. The final lysate was kept on ice. Protein concentrations were quantified using the Pierce™ 660 nm Protein Assay Reagent (Thermo Scientific) according to the manufacturer's protocol. Samples were diluted 1:10 in milliQ water prior to analysis. Average protein concentrations were ~0.5 mg/mL for root lysates and 1–2 mg/mL for shoot lysates.

Photoaffinity experiments were conducted using two different reagents: Photoaffinity Reagent **18** (Custom Linker) and Photoaffinity Reagent **19** (Commercial Linker). For all experiments with Photoaffinity Reagent **19** (Commercial Linker), extraction buffers without DTT were used. Streptavidin-coated magnetic beads (90 µL) were washed twice with 600 µL of wash buffer (composition provided in Chapter 3.1) using a magnetic rack to remove the supernatant after each wash. For competition experiments, 700 µL of lysate were pre-incubated with 50 µL of competitor stock solution (10 mM, either 4-PP-$InsP_5$ or 6-PP-$InsP_5$) for 45 minutes at 4°C under rotation. For control experiments without competitor, 50 µL of water were added instead and incubated under identical conditions. Subsequently, 5 µL of the capture compound stock solution (400 µM) were added and the mixture incubated for 1 hour at 4°C under rotation. Crosslinking was performed by irradiating the mixture under a 365 nm UV lamp for 30 minutes at 4°C. The crosslinked mixture was then added to the prepared beads and incubated for 1 hour at 4°C under rotation. After binding, the beads were washed twice with 600 µL of wash buffer (each wash: magnetic rack, supernatant discarded). Following the final wash, the beads were frozen in liquid nitrogen and stored at -80°C until further processing.

### 4 Proteomics

#### 4.1 Sample preparation

##### 4.1.1 On-bead digest for Affi-Gel and Photoaffinity enrichments

Enriched proteins were submitted to an on-bead digestion using trypsin. In brief, dry beads were re-dissolved in 25 µL digestion buffer 1 (50 mM Tris, pH 7.5, 2M urea, 1mM DTT, 5 ng/µL trypsin) and incubated for 30 min at 30 °C in a Thermomixer with 400 rpm. Next, beads were pelleted and the supernatant was transferred to a fresh tube. Digestion buffer 2 (50 mM Tris, pH 7.5, 2M urea, 5 mM CAA) was added to the beads, after mixing the beads were pelleted, the supernatant was collected and combined with the previous one. The combined supernatants were then incubated overnight at 32 °C in a Thermomixer with 400 rpm; samples were protected from light during incubation. The digestion was stopped by adding 5 µL 20% TFA and desalted with $C_{18}$ Empore disk membranes according to the StageTip protocol[10] and dried in a vacuum evaporator.

##### 4.1.2 Sample preparation (in-solution digest for Affigel elutions)

The eluted protein mixtures were reduced with dithiothreitol, alkylated with chloroacetamide, and digested with trypsin (0.125 µg). These digested samples were desalted using with $C_{18}$ Empore disk membranes according to the StageTip protocol[10] and dried in a vacuum evaporator.

#### 4.2 LC-MS/MS data acquisition

Dried peptides were re-dissolved in 2% MeCN, 0.1% TFA (10 µL) for analysis. Streptavidin pulldown samples were analyzed using an EASY-nLC 1200 (Thermo Fisher) coupled to a Q Exactive Plus mass spectrometer (Thermo Fisher). Peptides were separated on 16 cm frit-less silica emitters (MSWil, 75 µm inner diameter), packed in-house with reversed-phase ReproSil-Pur $C_{18}$ AQ 1.9 µm resin (Dr. Maisch). Peptides were loaded on the column and eluted for 115 min using a segmented linear gradient of 5% to 95% solvent B (0 min : 5%B; 0-5 min -> 5%B; 5-65 min -> 20%B; 65-90 min ->35%B; 90-100 min -> 55%; 100-105 min ->95%, 105-115 min ->95%) (solvent A 0% MeCN, 0.1% FA; solvent B 80% MeCN, 0.1%FA) at a flow rate of 300 nL/min. Mass spectra were acquired in data-dependent acquisition mode with a TOP15 method. MS spectra were acquired in the Orbitrap analyzer with a mass range of 300–1750 m/z at a resolution of 70,000 FWHM and a target value of $3\times10^6$ ions. Precursors were selected with an isolation window of 1.3 m/z. HCD fragmentation was performed at a normalized collision energy of 25. MS/MS spectra were acquired with a target value of $10^5$ ions at a resolution of 17,500 FWHM, a maximum injection time (max.) of 55 ms and a fixed first mass of m/z 100. Peptides with a charge of +1, greater than

6, or with unassigned charge state were excluded from fragmentation for $MS^2$, dynamic exclusion for 30s prevented repeated selection of precursors.

Affi-Gel samples were analyzed using an Ultimate 3000 RSLC nano (Thermo Fisher) coupled to an Orbitrap Exploris 480 mass spectrometer equipped with a FAIMS Pro interface for Field asymmetric ion mobility separation (Thermo Fisher). Peptides were pre-concentrated on an Acclaim PepMap 100 pre-column (75 μM x 2 cm, $C_{18}$, 3 μM, 100 Å, Thermo Fisher) using the loading pump and buffer A** (water, 0.1% TFA) with a flow of 7 µl/min for 5 min. Peptides were separated on 16 cm frit-less silica emitters (MSWil, 75 µm inner diameter), packed in-house with reversed-phase ReproSil-Pur C18 AQ 1.9 µm resin (Dr. Maisch). Peptides were loaded on the column and eluted for 130 min using a segmented linear gradient of 5% to 95% solvent B (0 min : 5%B; 0-5 min -> 5%B; 5-65 min -> 20%B; 65-90 min ->35%B; 90-100 min -> 55%; 100-105 min ->95%, 105-115 min ->95%, 115-115.1 min -> 5%, 115.1-130 min ->5%) (solvent A 0% MeCN, 0.1% FA; solvent B 80% MeCN, 0.1%FA) at a flow rate of 300 nL/min. Mass spectra were acquired in data-dependent acquisition mode with a TOP_S method using a cycle time of 2 seconds. For Field asymmetric ion mobility separation (FAIMS) two compensation voltages (-45 and -60) were applied, the cycle time for each experiment was set to 1 second. MS spectra were acquired in the Orbitrap analyzer with a mass range of 320–1200 m/z at a resolution of 60,000 FWHM and a normalized AGC target of 300%. Precursors were filtered using the MIPS option (MIPS mode = peptide), the intensity threshold was set to 5000, Precursors were selected with an isolation window of 1.6 m/z. HCD fragmentation was performed at a normalized collision energy of 30%. MS/MS spectra were acquired with a target value of 75% ions at a resolution of 15,000 FWHM, at an automated injection time and a fixed first mass of m/z 100. Peptides with a charge of +1, greater than 6, or with unassigned charge state were excluded from fragmentation for $MS^2$, dynamic exclusion for 40s prevented repeated selection of precursors.

#### 4.3 Data analysis

Raw data were processed using MaxQuant software (version 1.6.3.4, http://www.maxquant.org/)[11] with label-free quantification (LFQ) and iBAQ enabled.[12]

MS/MS spectra were searched by the Andromeda search engine against a combined database containing the sequences from *A. thaliana* (TAIR10_pep_20101214*) and sequences of 248 common contaminant proteins and decoy sequences. Trypsin specificity was required and a maximum of two missed cleavages allowed. Minimal peptide length was set to seven amino acids. Carbamidomethylation of cysteine residues was set as fixed, oxidation of methionine and protein N-terminal acetylation as variable modifications. The match between runs option was enabled. Peptide-spectrum-matches and proteins were retained if they were below a false discovery rate of 1% in both cases.

Statistical analysis of the MaxLFQ values was carried out using Perseus (version 1.5.8.5, http://www.maxquant.org/). Quantified proteins were filtered for reverse hits and hits "identified by site" and MaxLFQ values were log2 transformed. After grouping samples by condition only those proteins were retained for the subsequent analysis that had two valid values in one of the conditions. Two-sample t-tests were performed using a permutation-based FDR of 5%. Alternatively, quantified proteins were grouped by condition and only those hits were retained that had three valid values in one of the conditions. Missing values were imputed from a normal distribution (1.8 downshift, separately for each column). Volcano plots were generated in Perseus using an FDR of 5% and an S0=1. The Perseus output was exported and further processed using Excel.

* ftp://ftp.arabidopsis.org/home/tair/Proteins/TAIR10_protein_lists/

### 5 Biophysical Validation of AtFHA2–Ligand Interactions

#### 5.1 Protein Purification

The coding sequence of AtFHA2 was cloned into an N-terminal His–MBP–tagged pET28b expression vector. *E. coli* RIL cells were transformed with the AtFHA2 construct. The bacterial transformants were cultured at 37 °C overnight, and the secondary culture was grown at 16 °C until the optical density at 600 nm ($OD_{600}$) reached ~0.8. For purification, cells were lysed in Tris–HCl buffer (500 mM NaCl, 50 mM Tris–HCl, pH 8.0, 5 % glycerol, 0.2 mM PMSF, 2 mM β-ME, and 0.1 % Tween-20). The lysates were centrifuged at 14 000 rpm for 45 min at 4 °C, and the supernatant was collected. The supernatant was then incubated with pre-equilibrated Ni–NTA beads (Qiagen, 30230) at 4 °C for 12 h. Bound protein was eluted using lysis buffer supplemented with 250 mM imidazole. For size-exclusion chromatography, the eluted fractions were injected into an FPLC system and passed through a HiLoad 16/600 Superdex 200 pg column pre-equilibrated with 1× PBS buffer. The eluted fractions were analyzed by SDS-PAGE and Coomassie blue staining. The final protein fractions were pooled and concentrated to 0.4–0.5 mg $mL^{-1}$. The purified protein was dialyzed against 1× PBS buffer prior to binding assays.

#### 5.2 Circular dichroism (CD)

CD measurements were carried out using a Chirascan-plus CD spectrometer. Protein samples were prepared in 1× PBS (pH 7.4). AtFHA2–His–MBP protein was dialyzed and concentrated to 13 μM. Experiments were performed at room temperature. The UV CD spectra were recorded in the presence or absence of $InsP_6$ (500 μM and 1 mM), in the range of 190–320 nm using a 0.1 cm quartz cuvette.

#### 5.3 Isothermal Titration Calorimetry

Isothermal titration calorimetry (ITC) experiments were performed using a MicroCal PEAQ-ITC instrument at 25 °C as described previously.[13] The recombinant AtFHA2 protein and ligand ($InsP_6$, 4-PP-$InsP_5$) solutions were dialysed against 1× PBS buffer (pH 7.4). Samples were degassed under vacuum for 10 min before loading into the instrument. A series of 19 injections was given as command. The initial injection (0.4 μL) was later excluded during analysis to account for artefacts. The binding affinity of AtFHA2 for $InsP_6$ and 4-PP-$InsP_5$ was estimated by injecting $InsP_6$ (750 μM) and 4-PP-$InsP_5$ (1 mM) into the AtFHA2 protein solution inside the cell. Control titrations of ligand and buffer were performed under similar conditions to check the heat change. The data were processed using the built-in software of MicroCal instruments (version 1.21).

### 7 Gene Ontology Term Enrichment Analysis by g:Profiler

**Table 1: GO-Term Enrichment Results of the Combined Affi-Gel Root Dataset (g:Profiler Output).**
The table summarizes the Affi-Gel root enrichment results, including both the on-bead and elution fractions.The results were obtained using g:Profiler.

| Source | Term Name | Term ID | $p_{adj}$ | $-log_{10}$ ($p_{adj}$) | Term Size | Intersection Size |
|---|---|---|---|---|---|---|
| GO:MF | phospholipid binding | GO:0005543 | 0,00061188 | 3,213333878 | 135 | 7 |
| GO:MF | phosphatidylinositol kinase activity | GO:0052742 | 0,002512644 | 2,599869077 | 30 | 4 |
| GO:MF | 1-phosphatidylinositol-4-phosphate 5-kinase activity | GO:0016308 | 0,003832301 | 2,416540376 | 11 | 3 |
| GO:MF | inositol-1,4,5-trisphosphate 6-kinase activity | GO:0000823 | 0,005617333 | 2,250469797 | 2 | 2 |
| GO:MF | inositol-1,4,5,6-tetrakisphosphate 3-kinase activity | GO:0000824 | 0,005617333 | 2,250469797 | 2 | 2 |
| GO:MF | inositol-1,2,3,4,6-pentakisphosphate 5-kinase activity | GO:0102732 | 0,005617333 | 2,250469797 | 2 | 2 |
| GO:MF | inositol-1,3,4,6-tetrakisphosphate 5-kinase activity | GO:0047326 | 0,005617333 | 2,250469797 | 2 | 2 |
| GO:MF | lipid kinase activity | GO:0001727 | 0,009861894 | 2,006039655 | 42 | 4 |
| GO:MF | lipid binding | GO:0008289 | 0,018509837 | 1,732597404 | 310 | 8 |
| GO:MF | binding | GO:0005488 | 0,032995384 | 1,481546816 | 15005 | 82 |
| GO:MF | 5'-3' RNA exonuclease activity | GO:0004534 | 0,033513749 | 1,474776985 | 4 | 2 |
| GO:BP | RNA processing | GO:0006396 | 7,63E-07 | 6,117268116 | 1086 | 22 |
| GO:BP | mRNA metabolic process | GO:0016071 | 2,14704E-06 | 5,66815932 | 569 | 16 |
| GO:BP | mRNA processing | GO:0006397 | 1,06837E-05 | 4,971278665 | 468 | 14 |
| GO:BP | ribosome biogenesis | GO:0042254 | 0,000117291 | 3,930734377 | 254 | 10 |
| GO:BP | nucleic acid metabolic process | GO:0090304 | 0,000213307 | 3,670994737 | 4239 | 40 |
| GO:BP | RNA metabolic process | GO:0016070 | 0,000242699 | 3,614931285 | 3742 | 37 |
| GO:BP | RNA biosynthetic process | GO:0032774 | 0,000710247 | 3,148590784 | 3396 | 34 |
| GO:BP | nucleic acid biosynthetic process | GO:0141187 | 0,001183156 | 2,926958144 | 3471 | 34 |
| GO:BP | rRNA processing | GO:0006364 | 0,001546271 | 2,810714302 | 194 | 8 |
| GO:BP | rRNA metabolic process | GO:0016072 | 0,001869174 | 2,72835023 | 199 | 8 |
| GO:BP | vesicle-mediated transport | GO:0016192 | 0,002278804 | 2,642293021 | 437 | 11 |
| GO:BP | nucleobase-containing compound metabolic process | GO:0006139 | 0,002328473 | 2,632928784 | 4634 | 40 |
| GO:BP | RNA splicing | GO:0008380 | 0,002456317 | 2,609715611 | 355 | 10 |
| GO:BP | ribonucleoprotein complex biogenesis | GO:0022613 | 0,003141115 | 2,502916231 | 365 | 10 |
| GO:BP | endocytosis | GO:0006897 | 0,003637191 | 2,439233844 | 57 | 5 |
| GO:BP | nucleobase-containing compound biosynthetic process | GO:0034654 | 0,004957258 | 2,304758461 | 3696 | 34 |
| GO:BP | clathrin-dependent endocytosis | GO:0072583 | 0,008195186 | 2,086441172 | 32 | 4 |
| GO:BP | receptor-mediated endocytosis | GO:0006898 | 0,009294689 | 2,031765119 | 33 | 4 |
| GO:BP | phosphatidylinositol metabolic process | GO:0046488 | 0,013204288 | 1,879284998 | 74 | 5 |
| GO:BP | negative regulation of clathrin-dependent endocytosis | GO:1900186 | 0,014061189 | 1,85197795 | 2 | 2 |
| GO:BP | negative regulation of receptor-mediated endocytosis | GO:0048261 | 0,014061189 | 1,85197795 | 2 | 2 |
| GO:BP | negative regulation of endocytosis | GO:0045806 | 0,014061189 | 1,85197795 | 2 | 2 |
| GO:BP | gene expression | GO:0010467 | 0,026040601 | 1,584348996 | 5292 | 41 |
| GO:BP | import into cell | GO:0098657 | 0,027402432 | 1,562210895 | 86 | 5 |
| GO:BP | mRNA splicing, via spliceosome | GO:0000398 | 0,040063977 | 1,397245945 | 223 | 7 |
| GO:BP | regulation of receptor-mediated endocytosis | GO:0048259 | 0,042064323 | 1,376086094 | 3 | 2 |
| GO:BP | regulation of clathrin-dependent endocytosis | GO:2000369 | 0,042064323 | 1,376086094 | 3 | 2 |
| GO:CC | nucleus | GO:0005634 | 4,31E-10 | 9,36598358 | 5696 | 61 |
| GO:CC | intracellular membrane-bounded organelle | GO:0043231 | 0,000561056 | 3,250993526 | 11808 | 79 |
| GO:CC | membrane-bounded organelle | GO:0043227 | 0,00060015 | 3,221740532 | 11823 | 79 |
| GO:CC | nuclear protein-containing complex | GO:0140513 | 0,002326996 | 2,63320443 | 651 | 13 |
| GO:CC | intracellular anatomical structure | GO:0005622 | 0,004042179 | 2,39338444 | 13505 | 84 |
| GO:CC | organelle lumen | GO:0043233 | 0,004122886 | 2,384798645 | 1245 | 18 |
| GO:CC | intracellular organelle lumen | GO:0070013 | 0,004122886 | 2,384798645 | 1245 | 18 |
| GO:CC | membrane-enclosed lumen | GO:0031974 | 0,004122886 | 2,384798645 | 1245 | 18 |
| GO:CC | intracellular organelle | GO:0043229 | 0,004878838 | 2,311683608 | 12312 | 79 |
| GO:CC | organelle | GO:0043226 | 0,005165761 | 2,286865705 | 12326 | 79 |
| GO:CC | cell plate | GO:0009504 | 0,006486978 | 2,1879576 | 41 | 4 |
| GO:CC | nuclear lumen | GO:0031981 | 0,006813004 | 2,16666136 | 1054 | 16 |
| GO:CC | clathrin-coated vesicle | GO:0030136 | 0,039645122 | 1,401810244 | 65 | 4 |
| KEGG | Inositol phosphate metabolism | KEGG:00562 | 1,64817E-05 | 4,782997511 | 76 | 7 |
| KEGG | Ribosome biogenesis in eukaryotes | KEGG:03008 | 8,15701E-05 | 4,088469182 | 96 | 7 |
| KEGG | Phosphatidylinositol signaling system | KEGG:04070 | 0,000246659 | 3,60790327 | 74 | 6 |
| KEGG | Spliceosome | KEGG:03040 | 0,001518138 | 2,818688798 | 205 | 8 |
| KEGG | Endocytosis | KEGG:04144 | 0,017318747 | 1,76148353 | 158 | 6 |

**Table 2: GO-Term Enrichment Results of the Combined Affi-Gel Shoot Dataset (g:Profiler Output).**
The table summarizes the Affi-Gel shoot enrichment results, including both the on-bead and elution fractions.The results were obtained using g:Profiler.

| *Source* | *Term Name* | *Term ID* | $p_{adj}$ | $-log_{10}(p_{adj})$ | *Term Size* | *Intersection Size* |
|---|---|---|---|---|---|---|
| GO:MF | lipid binding | GO:0008289 | 1,13524E-05 | 4,944913233 | 310 | 11 |
| GO:MF | phosphatidylinositol kinase activity | GO:0052742 | 3,52182E-05 | 4,453232499 | 30 | 5 |
| GO:MF | lipid kinase activity | GO:0001727 | 0,00020265 | 3,693254243 | 42 | 5 |
| GO:MF | binding | GO:0005488 | 0,000655415 | 3,183483876 | 15005 | 78 |
| GO:MF | protein binding | GO:0005515 | 0,002910719 | 2,535999729 | 7655 | 50 |
| GO:MF | 1-phosphatidylinositol-4-phosphate 5-kinase activity | GO:0016308 | 0,003080756 | 2,511342695 | 11 | 3 |
| GO:MF | phospholipid binding | GO:0005543 | 0,004928159 | 2,307315254 | 135 | 6 |
| GO:MF | phosphatidylinositol binding | GO:0035091 | 0,005098986 | 2,292516198 | 80 | 5 |
| GO:BP | phosphatidylinositol metabolic process | GO:0046488 | 1,28318E-05 | 4,8917124 | 74 | 7 |
| GO:BP | vesicle-mediated transport | GO:0016192 | 0,000118903 | 3,924806722 | 437 | 12 |
| GO:BP | phosphatidylinositol phosphate biosynthetic process | GO:0046854 | 0,00047896 | 3,31970079 | 18 | 4 |
| GO:BP | glycerophospholipid metabolic process | GO:0006650 | 0,000810394 | 3,091303762 | 135 | 7 |
| GO:BP | post-Golgi vesicle-mediated transport | GO:0006892 | 0,002672265 | 2,573120513 | 27 | 4 |
| GO:BP | glycerolipid metabolic process | GO:0046486 | 0,004056063 | 2,391895323 | 172 | 7 |
| GO:BP | RNA biosynthetic process | GO:0032774 | 0,008285983 | 2,081655941 | 3396 | 30 |
| GO:BP | nucleic acid biosynthetic process | GO:0141187 | 0,012807843 | 1,892524013 | 3471 | 30 |
| GO:BP | phosphatidylinositol biosynthetic process | GO:0006661 | 0,016278415 | 1,788387894 | 42 | 4 |
| GO:BP | phospholipid metabolic process | GO:0006644 | 0,017259674 | 1,762967413 | 215 | 7 |
| GO:BP | regulation of cellular process | GO:0050794 | 0,018150899 | 1,74110187 | 4692 | 36 |
| GO:BP | nucleobase-containing compound biosynthetic process | GO:0034654 | 0,04337145 | 1,36279606 | 3696 | 30 |
| GO:BP | regulation of biological process | GO:0050789 | 0,047079995 | 1,327163593 | 5102 | 37 |
| GO:CC | clathrin-coated vesicle | GO:0030136 | 8,35013E-05 | 4,078306864 | 65 | 6 |
| GO:CC | coated vesicle | GO:0030135 | 0,000995281 | 3,00205429 | 99 | 6 |
| GO:CC | preribosome, large subunit precursor | GO:0030687 | 0,04305172 | 1,366009491 | 6 | 2 |
| GO:CC | nucleus | GO:0005634 | 0,044952348 | 1,347247622 | 5696 | 43 |
| KEGG | Endocytosis | KEGG:04144 | 1,82042E-06 | 5,73982948 | 158 | 9 |
| KEGG | Inositol phosphate metabolism | KEGG:00562 | 2,53274E-06 | 5,596409471 | 76 | 7 |
| KEGG | Phosphatidylinositol signaling system | KEGG:04070 | 0,001023764 | 2,989800318 | 74 | 5 |

**Table 3: GO-Term Enrichment Results of the Combined Photoaffinity Root Dataset (g:Profiler Output).**
The table summarizes the results from all photoaffinity shoot conditions, combining both the custom and commercial linker reagents as well as the different competition controls. The results were obtained using g:Profiler.

| Source | *Term Name* | *Term ID* | $p_{adj}$ | $-log_{10}(p_{adj})$ | *Term Size* | *Intersection Size* |
|---|---|---|---|---|---|---|
| GO:MF | RNA binding | GO:0003723 | 2,02E-14 | 13,69512889 | 2517 | 114 |
| GO:MF | transcription coregulator activity | GO:0003712 | 1,65E-11 | 10,78305308 | 108 | 20 |
| GO:MF | protein-macromolecule adaptor activity | GO:0030674 | 1,70E-10 | 9,769365657 | 202 | 25 |
| GO:MF | binding | GO:0005488 | 6,42E-09 | 8,192677769 | 15005 | 369 |
| GO:MF | RNA helicase activity | GO:0003724 | 1,53E-08 | 7,81520633 | 92 | 16 |
| GO:MF | ATP-dependent activity, acting on RNA | GO:0008186 | 1,53E-08 | 7,81520633 | 92 | 16 |
| GO:MF | mRNA binding | GO:0003729 | 2,89E-08 | 7,538641065 | 1619 | 73 |
| GO:MF | protein binding | GO:0005515 | 2,40021E-06 | 5,619749888 | 7655 | 214 |
| GO:MF | helicase activity | GO:0004386 | 2,68991E-05 | 4,570262555 | 214 | 19 |
| GO:MF | ATP-dependent activity | GO:0140657 | 4,59422E-05 | 4,337788245 | 858 | 42 |
| GO:MF | organic cyclic compound binding | GO:0097159 | 4,75589E-05 | 4,322767878 | 8337 | 223 |
| GO:MF | hydrolase activity, acting on acid anhydrides | GO:0016817 | 5,28932E-05 | 4,276600465 | 831 | 41 |
| GO:MF | hydrolase activity, acting on acid anhydrides, in phosphorus-containing anhydrides | GO:0016818 | 0,000121754 | 3,914518533 | 826 | 40 |
| GO:MF | ribonucleoside triphosphate phosphatase activity | GO:0017111 | 0,000239319 | 3,621023083 | 753 | 37 |
| GO:MF | pyrophosphatase activity | GO:0016462 | 0,000244522 | 3,611682106 | 817 | 39 |
| GO:MF | nucleic acid binding | GO:0003676 | 0,000464365 | 3,333140224 | 5097 | 147 |
| GO:MF | ATP hydrolysis activity | GO:0016887 | 0,00107481 | 2,96866823 | 552 | 29 |
| GO:MF | phosphatidylinositol kinase activity | GO:0052742 | 0,008322077 | 2,079768258 | 30 | 6 |
| GO:MF | modified amino acid binding | GO:0072341 | 0,009708388 | 2,012852895 | 19 | 5 |
| GO:MF | actin binding | GO:0003779 | 0,010637848 | 1,973146219 | 120 | 11 |
| GO:MF | small molecule binding | GO:0036094 | 0,011458522 | 1,940871387 | 6758 | 177 |
| GO:MF | ion binding | GO:0043167 | 0,013331164 | 1,875131916 | 6637 | 174 |
| GO:MF | 1-phosphatidylinositol-4-phosphate 5-kinase activity | GO:0016308 | 0,01547963 | 1,810239433 | 11 | 4 |
| GO:MF | anion binding | GO:0043168 | 0,016576497 | 1,780507233 | 3218 | 96 |
| GO:MF | carbohydrate derivative binding | GO:0097367 | 0,019448459 | 1,711114795 | 2856 | 87 |

| | | | | | | |
|---|---|---|---|---|---|---|
| GO:MF | phospholipase D activity | GO:0004630 | 0,022847593 | 1,641159541 | 12 | 4 |
| GO:MF | N-acylphosphatidylethanolamine-specific phospholipase D activity | GO:0070290 | 0,022847593 | 1,641159541 | 12 | 4 |
| GO:MF | nucleotide binding | GO:0000166 | 0,024196841 | 1,616241322 | 3124 | 93 |
| GO:MF | isocitrate dehydrogenase (NAD+) activity | GO:0004449 | 0,025286834 | 1,597105538 | 5 | 3 |
| GO:MF | nucleoside phosphate binding | GO:1901265 | 0,025969866 | 1,585530285 | 3130 | 93 |
| GO:MF | purine ribonucleoside triphosphate binding | GO:0035639 | 0,027790238 | 1,556107727 | 2678 | 82 |
| GO:MF | heterocyclic compound binding | GO:1901363 | 0,042814577 | 1,368408342 | 3258 | 95 |
| GO:BP | mRNA metabolic process | GO:0016071 | 3,17E-32 | 31,49881886 | 569 | 69 |
| GO:BP | RNA splicing | GO:0008380 | 1,00E-31 | 30,99984409 | 355 | 56 |
| GO:BP | mRNA processing | GO:0006397 | 6,91E-31 | 30,16053375 | 468 | 62 |
| GO:BP | RNA processing | GO:0006396 | 2,17E-28 | 27,66362373 | 1086 | 88 |
| GO:BP | mRNA splicing, via spliceosome | GO:0000398 | 5,32E-20 | 19,27404242 | 223 | 36 |
| GO:BP | RNA splicing, via transesterification reactions | GO:0000375 | 2,99E-19 | 18,52439684 | 250 | 37 |
| GO:BP | RNA splicing, via transesterification reactions with bulged adenosine as nucleophile | GO:0000377 | 2,99E-19 | 18,52439684 | 250 | 37 |
| GO:BP | nucleobase-containing compound biosynthetic process | GO:0034654 | 6,31E-15 | 14,19998472 | 3696 | 143 |
| GO:BP | RNA biosynthetic process | GO:0032774 | 5,97E-14 | 13,22374447 | 3396 | 133 |
| GO:BP | RNA metabolic process | GO:0016070 | 3,04E-13 | 12,516669 | 3742 | 140 |
| GO:BP | nucleic acid biosynthetic process | GO:0141187 | 3,59E-13 | 12,44430754 | 3471 | 133 |
| GO:BP | nucleobase-containing compound metabolic process | GO:0006139 | 6,72E-12 | 11,17281248 | 4634 | 158 |
| GO:BP | nucleic acid metabolic process | GO:0090304 | 2,99E-10 | 9,524939431 | 4239 | 144 |
| GO:BP | biosynthetic process | GO:0009058 | 1,73E-07 | 6,762698536 | 7320 | 203 |
| GO:BP | gene expression | GO:0010467 | 3,09086E-06 | 5,509920284 | 5292 | 155 |
| GO:BP | macromolecule biosynthetic process | GO:0009059 | 4,79102E-06 | 5,319571655 | 5736 | 164 |
| GO:BP | ribonucleoprotein complex biogenesis | GO:0022613 | 9,00489E-06 | 5,045521383 | 365 | 26 |
| GO:BP | response to temperature stimulus | GO:0009266 | 2,97416E-05 | 4,526635761 | 644 | 35 |
| GO:BP | rRNA processing | GO:0006364 | 0,000877378 | 3,056813351 | 194 | 16 |
| GO:BP | protein folding | GO:0006457 | 0,001101049 | 2,958193368 | 152 | 14 |
| GO:BP | response to cold | GO:0009409 | 0,001174159 | 2,930272905 | 410 | 24 |
| GO:BP | rRNA metabolic process | GO:0016072 | 0,001227632 | 2,910931686 | 199 | 16 |
| GO:BP | negative regulation of biosynthetic process | GO:0009890 | 0,001305056 | 2,884370903 | 658 | 32 |
| GO:BP | cellular component biogenesis | GO:0044085 | 0,001778875 | 2,749854539 | 1115 | 45 |
| GO:BP | negative regulation of macromolecule biosynthetic process | GO:0010558 | 0,001872111 | 2,727668363 | 637 | 31 |
| GO:BP | negative regulation of macromolecule metabolic process | GO:0010605 | 0,00293305 | 2,532680538 | 684 | 32 |
| GO:BP | organophosphate metabolic process | GO:0019637 | 0,003050961 | 2,515563354 | 620 | 30 |
| GO:BP | protein-RNA complex assembly | GO:0022618 | 0,003103596 | 2,508134809 | 122 | 12 |
| GO:BP | endoplasmic reticulum tubular network organization | GO:0071786 | 0,003255647 | 2,487362626 | 7 | 4 |
| GO:BP | protein-RNA complex organization | GO:0071826 | 0,003678064 | 2,43438071 | 124 | 12 |
| GO:BP | negative regulation of metabolic process | GO:0009892 | 0,004770901 | 2,321399583 | 734 | 33 |
| GO:BP | miRNA processing | GO:0035196 | 0,005885259 | 2,230234407 | 26 | 6 |
| GO:BP | primary metabolic process | GO:0044238 | 0,006921356 | 2,159808782 | 10896 | 254 |
| GO:BP | ribosome biogenesis | GO:0042254 | 0,00699659 | 2,155113589 | 254 | 17 |
| GO:BP | glycerophospholipid metabolic process | GO:0006650 | 0,008830282 | 2,054025423 | 135 | 12 |
| GO:BP | response to chemical | GO:0042221 | 0,010235903 | 1,989873845 | 2940 | 88 |
| GO:BP | response to stimulus | GO:0050896 | 0,011708611 | 1,931494604 | 6002 | 155 |
| GO:BP | cellular process | GO:0009987 | 0,014707193 | 1,832470216 | 16600 | 355 |
| GO:BP | glycerolipid metabolic process | GO:0046486 | 0,02207413 | 1,65611641 | 172 | 13 |
| GO:BP | nuclear-transcribed mRNA catabolic process, nonsense-mediated decay | GO:0000184 | 0,022710923 | 1,643765216 | 20 | 5 |
| GO:BP | regulation of RNA splicing | GO:0043484 | 0,037367565 | 1,427505196 | 22 | 5 |
| GO:BP | endoplasmic reticulum organization | GO:0007029 | 0,037367565 | 1,427505196 | 22 | 5 |
| GO:BP | phosphatidylcholine metabolic process | GO:0046470 | 0,046994638 | 1,327951688 | 23 | 5 |
| GO:CC | intracellular anatomical structure | GO:0005622 | 3,61E-30 | 29,44254694 | 13505 | 414 |
| GO:CC | nuclear protein-containing complex | GO:0140513 | 5,44E-28 | 27,2643184 | 651 | 73 |
| GO:CC | spliceosomal complex | GO:0005681 | 5,99E-27 | 26,22233118 | 130 | 36 |
| GO:CC | intracellular organelle | GO:0043229 | 2,06E-23 | 22,68603522 | 12312 | 382 |
| GO:CC | organelle | GO:0043226 | 2,80E-23 | 22,55337807 | 12326 | 382 |
| GO:CC | intracellular membrane-bounded organelle | GO:0043231 | 3,66E-22 | 21,43637355 | 11808 | 370 |
| GO:CC | membrane-bounded organelle | GO:0043227 | 5,04E-22 | 21,29781857 | 11823 | 370 |
| GO:CC | mediator complex | GO:0016592 | 6,11E-20 | 19,21417321 | 49 | 21 |
| GO:CC | nuclear speck | GO:0016607 | 2,01E-19 | 18,69748867 | 83 | 25 |
| GO:CC | organelle lumen | GO:0043233 | 1,55E-18 | 17,81023149 | 1245 | 86 |
| GO:CC | intracellular organelle lumen | GO:0070013 | 1,55E-18 | 17,81023149 | 1245 | 86 |
| GO:CC | membrane-enclosed lumen | GO:0031974 | 1,55E-18 | 17,81023149 | 1245 | 86 |
| GO:CC | nuclear lumen | GO:0031981 | 8,78E-18 | 17,05629516 | 1054 | 77 |
| GO:CC | nuclear body | GO:0016604 | 1,74E-17 | 16,75853551 | 119 | 27 |

| | | | | | | |
|---|---|---|---|---|---|---|
| GO:CC | nucleoplasm | GO:0005654 | 4,97E-17 | 16,30393192 | 357 | 43 |
| GO:CC | nucleus | GO:0005634 | 1,25E-16 | 15,90309291 | 5696 | 218 |
| GO:CC | cytosol | GO:0005829 | 1,22E-11 | 10,91447295 | 2420 | 112 |
| GO:CC | protein-containing complex | GO:0032991 | 2,19E-11 | 10,65882475 | 2604 | 117 |
| GO:CC | cytoplasm | GO:0005737 | 6,16E-09 | 8,210168928 | 9524 | 286 |
| GO:CC | cellular anatomical structure | GO:0110165 | 1,13E-08 | 7,945883095 | 18498 | 452 |
| GO:CC | nucleolus | GO:0005730 | 5,90E-08 | 7,228940902 | 701 | 45 |
| GO:CC | ribonucleoprotein complex | GO:1990904 | 4,86E-07 | 6,313104503 | 831 | 48 |
| GO:CC | cell periphery | GO:0071944 | 0,000200638 | 3,697586241 | 2917 | 104 |
| GO:CC | plasma membrane | GO:0005886 | 0,000212532 | 3,672575208 | 2524 | 93 |
| GO:CC | plasmodesma | GO:0009506 | 0,000235819 | 3,627421307 | 900 | 44 |
| GO:CC | cell-cell junction | GO:0005911 | 0,000235819 | 3,627421307 | 900 | 44 |
| GO:CC | cell junction | GO:0030054 | 0,000235819 | 3,627421307 | 900 | 44 |
| GO:CC | anchoring junction | GO:0070161 | 0,000235819 | 3,627421307 | 900 | 44 |
| GO:CC | symplast | GO:0055044 | 0,000242777 | 3,614792872 | 901 | 44 |
| GO:CC | endoplasmic reticulum lumen | GO:0005788 | 0,003982869 | 2,399803965 | 40 | 7 |
| GO:CC | nuclear inner membrane | GO:0005637 | 0,006714494 | 2,172986727 | 10 | 4 |
| GO:CC | membraneless organelle | GO:0043228 | 0,007100135 | 2,148733369 | 2568 | 88 |
| GO:CC | intracellular membraneless organelle | GO:0043232 | 0,007100135 | 2,148733369 | 2568 | 88 |
| GO:CC | cytoplasmic side of endoplasmic reticulum membrane | GO:0098554 | 0,015204319 | 1,818033033 | 5 | 3 |
| KEGG | Spliceosome | KEGG:03040 | 4,60E-30 | 29,33749708 | 205 | 52 |
| KEGG | mRNA surveillance pathway | KEGG:03015 | 6,65509E-06 | 5,176846028 | 119 | 18 |
| KEGG | Phosphatidylinositol signaling system | KEGG:04070 | 0,048016644 | 1,318608199 | 74 | 9 |
| KEGG | Nucleocytoplasmic transport | KEGG:03013 | 0,049714598 | 1,303516071 | 105 | 11 |
| WP | TCA cycle Krebs cycle | WP:WP2624 | 0,012633814 | 1,898465536 | 29 | 4 |

**Table 4: GO-Term Enrichment Results of the Combined Photoaffinity Shoot Dataset (g:Profiler Output).** The table summarizes the results from all photoaffinity shoot conditions, combining both the custom and commercial linker reagents as well as the different competition controls. The results were obtained using g:Profiler.

| Source | Term Name | Term ID | $p_{adj}$ | $-\log_{10}(p_{adj})$ | Term Size | Intersection Size |
|---|---|---|---|---|---|---|
| GO:MF | catalytic activity | GO:0003824 | 3,20E-23 | 22,49543984 | 9210 | 362 |
| GO:MF | oxidoreductase activity | GO:0016491 | 6,69E-13 | 12,17477969 | 1475 | 92 |
| GO:MF | lyase activity | GO:0016829 | 6,65E-10 | 9,177256729 | 446 | 41 |
| GO:MF | carbon-carbon lyase activity | GO:0016830 | 3,10E-08 | 7,508720875 | 114 | 19 |
| GO:MF | intramolecular oxidoreductase activity | GO:0016860 | 1,15E-07 | 6,939051248 | 51 | 13 |
| GO:MF | isomerase activity | GO:0016853 | 6,62E-07 | 6,179080343 | 270 | 27 |
| GO:MF | small molecule binding | GO:0036094 | 1,01653E-06 | 5,992879808 | 6758 | 240 |
| GO:MF | oxidoreductase activity, acting on the CH-OH group of donors, NAD or NADP as acceptor | GO:0016616 | 2,86614E-06 | 5,542703185 | 148 | 19 |
| GO:MF | modified amino acid binding | GO:0072341 | 3,32693E-06 | 5,477955742 | 19 | 8 |
| GO:MF | ligase activity, forming carbon-carbon bonds | GO:0016885 | 6,83533E-06 | 5,165240779 | 9 | 6 |
| GO:MF | CoA carboxylase activity | GO:0016421 | 6,83533E-06 | 5,165240779 | 9 | 6 |
| GO:MF | transferase activity, transferring alkyl or aryl (other than methyl) groups | GO:0016765 | 8,46267E-06 | 5,072492828 | 158 | 19 |
| GO:MF | copper ion binding | GO:0005507 | 8,58318E-06 | 5,066351883 | 192 | 21 |
| GO:MF | oxidoreductase activity, acting on CH-OH group of donors | GO:0016614 | 2,66354E-05 | 4,574541365 | 187 | 20 |
| GO:MF | ligase activity | GO:0016874 | 3,04256E-05 | 4,516760938 | 283 | 25 |
| GO:MF | magnesium ion binding | GO:0000287 | 4,36546E-05 | 4,359969567 | 141 | 17 |
| GO:MF | carboxylic acid binding | GO:0031406 | 5,1285E-05 | 4,290009857 | 96 | 14 |
| GO:MF | organic acid binding | GO:0043177 | 5,1285E-05 | 4,290009857 | 96 | 14 |
| GO:MF | hydro-lyase activity | GO:0016836 | 5,18682E-05 | 4,285098767 | 82 | 13 |
| GO:MF | cation binding | GO:0043169 | 5,91847E-05 | 4,227790313 | 3961 | 151 |
| GO:MF | metal ion binding | GO:0046872 | 8,14554E-05 | 4,08907998 | 3948 | 150 |
| GO:MF | carboxy-lyase activity | GO:0016831 | 0,0001199 | 3,921181338 | 74 | 12 |
| GO:MF | ion binding | GO:0043167 | 0,000166943 | 3,777432604 | 6637 | 226 |
| GO:MF | amide binding | GO:0033218 | 0,000344326 | 3,46303037 | 32 | 8 |
| GO:MF | alkali metal ion binding | GO:0031420 | 0,000558352 | 3,253091932 | 16 | 6 |
| GO:MF | cobalt ion binding | GO:0050897 | 0,000817688 | 3,087412585 | 47 | 9 |
| GO:MF | glutathione transferase activity | GO:0004364 | 0,000863537 | 3,063719232 | 60 | 10 |
| GO:MF | monocarboxylic acid binding | GO:0033293 | 0,001009251 | 2,996001009 | 61 | 10 |
| GO:MF | oxidoreductase activity, acting on the aldehyde or oxo group of donors | GO:0016903 | 0,001009251 | 2,996001009 | 61 | 10 |
| GO:MF | salicylic acid binding | GO:1901149 | 0,001639619 | 2,78525705 | 28 | 7 |
| GO:MF | oligopeptide binding | GO:1900750 | 0,002316402 | 2,635186057 | 12 | 5 |
| GO:MF | glutathione binding | GO:0043295 | 0,002316402 | 2,635186057 | 12 | 5 |

| | | | | | | |
|---|---|---|---|---|---|---|
| GO:MF | vitamin binding | GO:0019842 | 0,002739842 | 2,562274449 | 133 | 14 |
| GO:MF | aldehyde dehydrogenase (NAD+) activity | GO:0004029 | 0,00368426 | 2,433649739 | 13 | 5 |
| GO:MF | aldehyde dehydrogenase [NAD(P)+] activity | GO:0004030 | 0,00368426 | 2,433649739 | 13 | 5 |
| GO:MF | acetyl-CoA carboxylase activity | GO:0003989 | 0,004365374 | 2,35997854 | 7 | 4 |
| GO:MF | potassium ion binding | GO:0030955 | 0,005609504 | 2,251075539 | 14 | 5 |
| GO:MF | pyruvate kinase activity | GO:0004743 | 0,005609504 | 2,251075539 | 14 | 5 |
| GO:MF | ligase activity, forming carbon-nitrogen bonds | GO:0016879 | 0,007551507 | 2,121966385 | 61 | 9 |
| GO:MF | protein domain specific binding | GO:0019904 | 0,012403055 | 1,906471327 | 97 | 11 |
| GO:MF | heterocyclic compound binding | GO:1901363 | 0,012675717 | 1,897027466 | 3258 | 119 |
| GO:MF | NAD binding | GO:0051287 | 0,01325226 | 1,877710054 | 81 | 10 |
| GO:MF | binding | GO:0005488 | 0,015069233 | 1,821908862 | 15005 | 431 |
| GO:MF | intramolecular oxidoreductase activity, interconverting aldoses and ketoses | GO:0016861 | 0,016259654 | 1,788888697 | 17 | 5 |
| GO:MF | NADPH binding | GO:0070402 | 0,020288671 | 1,6927464 | 4 | 3 |
| GO:MF | oxidoreductase activity, acting on the aldehyde or oxo group of donors, NAD or NADP as acceptor | GO:0016620 | 0,027324711 | 1,563444418 | 42 | 7 |
| GO:MF | anion binding | GO:0043168 | 0,028361904 | 1,547264616 | 3218 | 116 |
| GO:MF | carbon-oxygen lyase activity | GO:0016835 | 0,03043674 | 1,516601863 | 165 | 14 |
| GO:MF | sulfur compound binding | GO:1901681 | 0,030440997 | 1,516541133 | 30 | 6 |
| GO:MF | oxidoreductase activity, acting on single donors with incorporation of molecular oxygen, incorporation of one atom of oxygen (internal monooxygenases or internal mixed function oxidases) | GO:0016703 | 0,049746736 | 1,303235407 | 5 | 3 |
| GO:MF | oxidoreductase activity, acting on metal ions, oxygen as acceptor | GO:0016724 | 0,049746736 | 1,303235407 | 5 | 3 |
| GO:MF | ferroxidase activity | GO:0004322 | 0,049746736 | 1,303235407 | 5 | 3 |
| GO:BP | small molecule metabolic process | GO:0044281 | 2,07E-52 | 51,68311778 | 1602 | 161 |
| GO:BP | carboxylic acid metabolic process | GO:0019752 | 7,21E-38 | 37,14189713 | 955 | 108 |
| GO:BP | oxoacid metabolic process | GO:0043436 | 4,75E-37 | 36,32355251 | 1067 | 113 |
| GO:BP | organic acid metabolic process | GO:0006082 | 5,20E-37 | 36,28413547 | 1068 | 113 |
| GO:BP | small molecule biosynthetic process | GO:0044283 | 9,04E-26 | 25,04384437 | 686 | 77 |
| GO:BP | organophosphate metabolic process | GO:0019637 | 4,38E-24 | 23,35878749 | 620 | 71 |
| GO:BP | amino acid metabolic process | GO:0006520 | 1,90E-23 | 22,72165849 | 435 | 59 |
| GO:BP | nucleoside phosphate metabolic process | GO:0006753 | 4,39E-23 | 22,35710247 | 366 | 54 |
| GO:BP | nucleobase-containing small molecule metabolic process | GO:0055086 | 7,27E-21 | 20,13825143 | 406 | 54 |
| GO:BP | proteinogenic amino acid metabolic process | GO:0170039 | 2,99E-19 | 18,52375459 | 271 | 43 |
| GO:BP | L-amino acid metabolic process | GO:0170033 | 4,03E-19 | 18,39473372 | 273 | 43 |
| GO:BP | alpha-amino acid metabolic process | GO:1901605 | 4,60E-19 | 18,33695671 | 317 | 46 |
| GO:BP | carboxylic acid biosynthetic process | GO:0046394 | 5,91E-18 | 17,22843559 | 556 | 59 |
| GO:BP | organic acid biosynthetic process | GO:0016053 | 5,91E-18 | 17,22843559 | 556 | 59 |
| GO:BP | cellular process | GO:0009987 | 1,83E-16 | 15,73689573 | 16600 | 500 |
| GO:BP | nucleotide metabolic process | GO:0009117 | 1,95E-16 | 15,70924339 | 274 | 40 |
| GO:BP | metabolic process | GO:0008152 | 4,81E-16 | 15,31754934 | 12629 | 417 |
| GO:BP | ribonucleotide metabolic process | GO:0009259 | 5,71E-16 | 15,2434157 | 185 | 33 |
| GO:BP | ribose phosphate metabolic process | GO:0019693 | 5,71E-16 | 15,2434157 | 185 | 33 |
| GO:BP | carbohydrate derivative metabolic process | GO:1901135 | 4,99E-15 | 14,30220571 | 659 | 60 |
| GO:BP | purine-containing compound metabolic process | GO:0072521 | 4,29E-14 | 13,36718297 | 271 | 37 |
| GO:BP | generation of precursor metabolites and energy | GO:0006091 | 8,54E-14 | 13,06870547 | 464 | 48 |
| GO:BP | organophosphate biosynthetic process | GO:0090407 | 1,14E-13 | 12,94134228 | 413 | 45 |
| GO:BP | amino acid biosynthetic process | GO:0008652 | 1,28E-13 | 12,89437219 | 220 | 33 |
| GO:BP | monocarboxylic acid metabolic process | GO:0032787 | 1,46E-13 | 12,8349791 | 489 | 49 |
| GO:BP | nucleoside phosphate biosynthetic process | GO:1901293 | 1,93E-13 | 12,71519111 | 223 | 33 |
| GO:BP | alpha-amino acid biosynthetic process | GO:1901607 | 2,00E-13 | 12,69849906 | 195 | 31 |
| GO:BP | purine nucleotide metabolic process | GO:0006163 | 1,94E-12 | 11,71220149 | 211 | 31 |
| GO:BP | proteinogenic amino acid biosynthetic process | GO:0170038 | 2,75E-12 | 11,56139819 | 158 | 27 |
| GO:BP | L-amino acid biosynthetic process | GO:0170034 | 2,75E-12 | 11,56139819 | 158 | 27 |
| GO:BP | purine ribonucleotide metabolic process | GO:0009150 | 6,11E-12 | 11,2136897 | 163 | 27 |
| GO:BP | photosynthesis | GO:0015979 | 3,10E-11 | 10,50826833 | 281 | 34 |
| GO:BP | nucleoside diphosphate metabolic process | GO:0009132 | 4,21E-11 | 10,37550694 | 78 | 19 |
| GO:BP | nucleoside triphosphate metabolic process | GO:0009141 | 4,55E-11 | 10,34233055 | 136 | 24 |
| GO:BP | nucleoside phosphate catabolic process | GO:1901292 | 6,94E-11 | 10,15858835 | 80 | 19 |
| GO:BP | nucleoside diphosphate catabolic process | GO:0009134 | 1,55E-10 | 9,808396109 | 73 | 18 |
| GO:BP | ribonucleoside triphosphate metabolic process | GO:0009199 | 1,71E-10 | 9,766910812 | 131 | 23 |
| GO:BP | pyridine-containing compound catabolic process | GO:0072526 | 2,01E-10 | 9,697498488 | 74 | 18 |
| GO:BP | pyruvate metabolic process | GO:0006090 | 2,58E-10 | 9,588457826 | 75 | 18 |
| GO:BP | sulfur compound metabolic process | GO:0006790 | 2,68E-10 | 9,571868421 | 320 | 35 |
| GO:BP | ribonucleoside diphosphate metabolic process | GO:0009185 | 3,30E-10 | 9,481217647 | 76 | 18 |

| | | | | | | |
|---|---|---|---|---|---|---|
| GO:BP | organophosphate catabolic process | GO:0046434 | 8,40E-10 | 9,075557247 | 91 | 19 |
| GO:BP | purine ribonucleoside diphosphate catabolic process | GO:0009181 | 1,57E-09 | 8,803971232 | 72 | 17 |
| GO:BP | ribonucleotide catabolic process | GO:0009261 | 1,57E-09 | 8,803971232 | 72 | 17 |
| GO:BP | purine nucleoside diphosphate catabolic process | GO:0009137 | 1,57E-09 | 8,803971232 | 72 | 17 |
| GO:BP | glycolytic process | GO:0006096 | 1,57E-09 | 8,803971232 | 72 | 17 |
| GO:BP | ADP metabolic process | GO:0046031 | 1,57E-09 | 8,803971232 | 72 | 17 |
| GO:BP | ADP catabolic process | GO:0046032 | 1,57E-09 | 8,803971232 | 72 | 17 |
| GO:BP | ribonucleoside diphosphate catabolic process | GO:0009191 | 1,57E-09 | 8,803971232 | 72 | 17 |
| GO:BP | purine ribonucleotide catabolic process | GO:0009154 | 1,57E-09 | 8,803971232 | 72 | 17 |
| GO:BP | purine nucleotide catabolic process | GO:0006195 | 2,00E-09 | 8,698964776 | 73 | 17 |
| GO:BP | pyridine nucleotide catabolic process | GO:0019364 | 2,00E-09 | 8,698964776 | 73 | 17 |
| GO:BP | pyridine-containing compound metabolic process | GO:0072524 | 2,33E-09 | 8,631809349 | 134 | 22 |
| GO:BP | nicotinamide nucleotide metabolic process | GO:0046496 | 2,41E-09 | 8,617823832 | 121 | 21 |
| GO:BP | purine ribonucleoside diphosphate metabolic process | GO:0009179 | 3,20E-09 | 8,49423014 | 75 | 17 |
| GO:BP | purine nucleoside diphosphate metabolic process | GO:0009135 | 3,20E-09 | 8,49423014 | 75 | 17 |
| GO:BP | pyridine nucleotide metabolic process | GO:0019362 | 3,35E-09 | 8,474822457 | 123 | 21 |
| GO:BP | nucleotide catabolic process | GO:0009166 | 5,06E-09 | 8,296182896 | 77 | 17 |
| GO:BP | biosynthetic process | GO:0009058 | 5,16E-09 | 8,286987181 | 7320 | 261 |
| GO:BP | ATP metabolic process | GO:0046034 | 5,46E-09 | 8,262619728 | 113 | 20 |
| GO:BP | purine nucleoside triphosphate metabolic process | GO:0009144 | 1,47E-08 | 7,832670964 | 119 | 20 |
| GO:BP | purine ribonucleoside triphosphate metabolic process | GO:0009205 | 1,47E-08 | 7,832670964 | 119 | 20 |
| GO:BP | response to cadmium ion | GO:0046686 | 1,52E-08 | 7,818935482 | 71 | 16 |
| GO:BP | carbohydrate derivative catabolic process | GO:1901136 | 2,45E-08 | 7,611318933 | 136 | 21 |
| GO:BP | purine-containing compound catabolic process | GO:0072523 | 3,33E-08 | 7,478050532 | 86 | 17 |
| GO:BP | cellular catabolic process | GO:0044248 | 3,40E-08 | 7,468958936 | 199 | 25 |
| GO:BP | catabolic process | GO:0009056 | 7,75E-08 | 7,110566135 | 1379 | 76 |
| GO:BP | pyruvate family amino acid metabolic process | GO:0009078 | 1,01E-07 | 6,996412388 | 31 | 11 |
| GO:BP | nucleobase-containing compound catabolic process | GO:0034655 | 1,22E-07 | 6,912194833 | 228 | 26 |
| GO:BP | purine-containing compound biosynthetic process | GO:0072522 | 5,61E-07 | 6,250928536 | 130 | 19 |
| GO:BP | response to metal ion | GO:0010038 | 1,40331E-06 | 5,852845239 | 219 | 24 |
| GO:BP | sulfur compound biosynthetic process | GO:0044272 | 2,85801E-06 | 5,543935758 | 175 | 21 |
| GO:BP | ribonucleoside monophosphate biosynthetic process | GO:0009156 | 6,78611E-06 | 5,168379161 | 44 | 11 |
| GO:BP | carbohydrate derivative biosynthetic process | GO:1901137 | 1,2391E-05 | 4,906892533 | 407 | 32 |
| GO:BP | response to cold | GO:0009409 | 1,47461E-05 | 4,831322648 | 410 | 32 |
| GO:BP | ribonucleoside monophosphate metabolic process | GO:0009161 | 1,8249E-05 | 4,738761464 | 48 | 11 |
| GO:BP | response to abiotic stimulus | GO:0009628 | 2,02636E-05 | 4,693282773 | 2131 | 95 |
| GO:BP | ribonucleotide biosynthetic process | GO:0009260 | 2,12935E-05 | 4,671753 | 99 | 15 |
| GO:BP | ribose phosphate biosynthetic process | GO:0046390 | 2,12935E-05 | 4,671753 | 99 | 15 |
| GO:BP | nucleotide biosynthetic process | GO:0009165 | 2,19852E-05 | 4,657869831 | 145 | 18 |
| GO:BP | dicarboxylic acid metabolic process | GO:0043648 | 2,82172E-05 | 4,549486798 | 87 | 14 |
| GO:BP | fatty acid metabolic process | GO:0006631 | 3,1262E-05 | 4,504983019 | 276 | 25 |
| GO:BP | nucleoside monophosphate biosynthetic process | GO:0009124 | 3,59212E-05 | 4,444649009 | 51 | 11 |
| GO:BP | L-leucine metabolic process | GO:0006551 | 4,32496E-05 | 4,364018021 | 23 | 8 |
| GO:BP | alpha-amino acid catabolic process | GO:1901606 | 5,54534E-05 | 4,256071777 | 65 | 12 |
| GO:BP | aromatic amino acid metabolic process | GO:0009072 | 5,85645E-05 | 4,232365631 | 92 | 14 |
| GO:BP | L-amino acid catabolic process | GO:0170035 | 6,88677E-05 | 4,161984297 | 43 | 10 |
| GO:BP | proteinogenic amino acid catabolic process | GO:0170040 | 6,88677E-05 | 4,161984297 | 43 | 10 |
| GO:BP | nucleoside monophosphate metabolic process | GO:0009123 | 0,000110738 | 3,955703575 | 69 | 12 |
| GO:BP | response to temperature stimulus | GO:0009266 | 0,00013894 | 3,857173226 | 644 | 40 |
| GO:BP | lipid metabolic process | GO:0006629 | 0,000161243 | 3,792518845 | 1098 | 57 |
| GO:BP | carbon fixation | GO:0015977 | 0,000179154 | 3,746773471 | 27 | 8 |
| GO:BP | amino acid catabolic process | GO:0009063 | 0,000180188 | 3,744274813 | 72 | 12 |
| GO:BP | branched-chain amino acid metabolic process | GO:0009081 | 0,000209786 | 3,678222655 | 48 | 10 |
| GO:BP | mRNA metabolic process | GO:0016071 | 0,000375459 | 3,42543732 | 569 | 36 |
| GO:BP | photosynthesis, dark reaction | GO:0019685 | 0,00042311 | 3,37354708 | 21 | 7 |
| GO:BP | reductive pentose-phosphate cycle | GO:0019253 | 0,00042311 | 3,37354708 | 21 | 7 |
| GO:BP | glutamine family amino acid biosynthetic process | GO:0009084 | 0,000441847 | 3,354727864 | 30 | 8 |
| GO:BP | aromatic amino acid family biosynthetic process | GO:0009073 | 0,000575656 | 3,239837114 | 66 | 11 |
| GO:BP | glutamine family amino acid metabolic process | GO:0009064 | 0,000605074 | 3,218191401 | 95 | 13 |
| GO:BP | cold acclimation | GO:0009631 | 0,000800756 | 3,096499874 | 55 | 10 |
| GO:BP | primary metabolic process | GO:0044238 | 0,000875541 | 3,057723411 | 10896 | 330 |
| GO:BP | acyl-CoA biosynthetic process | GO:0071616 | 0,001179221 | 2,928404772 | 24 | 7 |
| GO:BP | thioester biosynthetic process | GO:0035384 | 0,001179221 | 2,928404772 | 24 | 7 |
| GO:BP | monocarboxylic acid biosynthetic process | GO:0072330 | 0,00122317 | 2,912513235 | 249 | 21 |
| GO:BP | fatty acid biosynthetic process | GO:0006633 | 0,001306923 | 2,883750056 | 171 | 17 |
| GO:BP | pyruvate family amino acid catabolic process | GO:0009080 | 0,001801555 | 2,744352408 | 10 | 5 |
| GO:BP | response to stimulus | GO:0050896 | 0,002290901 | 2,639993738 | 6002 | 200 |

| | | | | | | |
|---|---|---|---|---|---|---|
| GO:BP | small molecule catabolic process | GO:0044282 | 0,002453104 | 2,610283955 | 179 | 17 |
| GO:BP | photosynthesis, light harvesting | GO:0009765 | 0,002495144 | 2,602904423 | 37 | 8 |
| GO:BP | mRNA processing | GO:0006397 | 0,002631742 | 2,579756737 | 468 | 30 |
| GO:BP | L-histidine metabolic process | GO:0006547 | 0,003234271 | 2,490223627 | 11 | 5 |
| GO:BP | imidazole-containing compound metabolic process | GO:0052803 | 0,003234271 | 2,490223627 | 11 | 5 |
| GO:BP | L-histidine biosynthetic process | GO:0000105 | 0,003234271 | 2,490223627 | 11 | 5 |
| GO:BP | lipid biosynthetic process | GO:0008610 | 0,003920353 | 2,406674807 | 630 | 36 |
| GO:BP | L-arginine biosynthetic process | GO:0006526 | 0,005429406 | 2,265247699 | 12 | 5 |
| GO:BP | UMP metabolic process | GO:0046049 | 0,005429406 | 2,265247699 | 12 | 5 |
| GO:BP | pyrimidine ribonucleoside monophosphate biosynthetic process | GO:0009174 | 0,005429406 | 2,265247699 | 12 | 5 |
| GO:BP | UMP biosynthetic process | GO:0006222 | 0,005429406 | 2,265247699 | 12 | 5 |
| GO:BP | pyruvate family amino acid biosynthetic process | GO:0009079 | 0,005688191 | 2,245025866 | 20 | 6 |
| GO:BP | phosphate-containing compound metabolic process | GO:0006796 | 0,005865341 | 2,231706759 | 2032 | 83 |
| GO:BP | phosphorus metabolic process | GO:0006793 | 0,00619887 | 2,207687446 | 2035 | 83 |
| GO:BP | amide metabolic process | GO:0043603 | 0,007974229 | 2,098311294 | 156 | 15 |
| GO:BP | pyrimidine ribonucleoside monophosphate metabolic process | GO:0009173 | 0,008639802 | 2,063496208 | 13 | 5 |
| GO:BP | response to chemical | GO:0042221 | 0,008758966 | 2,057547162 | 2940 | 110 |
| GO:BP | response to oxidative stress | GO:0006979 | 0,009946025 | 2,002350468 | 426 | 27 |
| GO:BP | endoplasmic reticulum tubular network organization | GO:0071786 | 0,010449825 | 1,980890992 | 7 | 4 |
| GO:BP | RNA splicing | GO:0008380 | 0,01051723 | 1,97809864 | 355 | 24 |
| GO:BP | cellular metabolic compound salvage | GO:0043094 | 0,011332415 | 1,945677539 | 73 | 10 |
| GO:BP | protein peptidyl-prolyl isomerization | GO:0000413 | 0,011954302 | 1,922475761 | 33 | 7 |
| GO:BP | toxin catabolic process | GO:0009407 | 0,01381589 | 1,85962112 | 46 | 8 |
| GO:BP | toxin metabolic process | GO:0009404 | 0,014400149 | 1,841633025 | 60 | 9 |
| GO:BP | response to toxic substance | GO:0009636 | 0,015266038 | 1,816273656 | 316 | 22 |
| GO:BP | regulation of fatty acid metabolic process | GO:0019217 | 0,018127187 | 1,74166958 | 24 | 6 |
| GO:BP | carboxylic acid catabolic process | GO:0046395 | 0,018729203 | 1,727480709 | 129 | 13 |
| GO:BP | organic acid catabolic process | GO:0016054 | 0,018729203 | 1,727480709 | 129 | 13 |
| GO:BP | L-leucine biosynthetic process | GO:0009098 | 0,019332728 | 1,713706863 | 15 | 5 |
| GO:BP | detoxification | GO:0098754 | 0,019466849 | 1,710704333 | 275 | 20 |
| GO:BP | pyrimidine ribonucleotide biosynthetic process | GO:0009220 | 0,023345566 | 1,631795591 | 25 | 6 |
| GO:BP | branched-chain amino acid biosynthetic process | GO:0009082 | 0,023345566 | 1,631795591 | 25 | 6 |
| GO:BP | pyrimidine ribonucleotide metabolic process | GO:0009218 | 0,023345566 | 1,631795591 | 25 | 6 |
| GO:BP | response to stress | GO:0006950 | 0,024699308 | 1,60731522 | 3733 | 131 |
| GO:BP | pyrimidine nucleoside monophosphate biosynthetic process | GO:0009130 | 0,02753816 | 1,560065079 | 16 | 5 |
| GO:BP | monosaccharide biosynthetic process | GO:0046364 | 0,030187824 | 1,520168193 | 51 | 8 |
| GO:BP | thioester metabolic process | GO:0035383 | 0,031664594 | 1,499426077 | 38 | 7 |
| GO:BP | acyl-CoA metabolic process | GO:0006637 | 0,031664594 | 1,499426077 | 38 | 7 |
| GO:BP | pyrimidine nucleoside monophosphate metabolic process | GO:0009129 | 0,03820521 | 1,417877414 | 17 | 5 |
| GO:BP | IMP metabolic process | GO:0046040 | 0,03820521 | 1,417877414 | 17 | 5 |
| GO:BP | IMP biosynthetic process | GO:0006188 | 0,03820521 | 1,417877414 | 17 | 5 |
| GO:BP | nucleotide salvage | GO:0043173 | 0,03820521 | 1,417877414 | 17 | 5 |
| GO:BP | water-soluble vitamin metabolic process | GO:0006767 | 0,039317825 | 1,40541051 | 84 | 10 |
| GO:BP | purine ribonucleoside monophosphate biosynthetic process | GO:0009168 | 0,046571057 | 1,331883907 | 28 | 6 |
| GO:BP | purine nucleoside monophosphate biosynthetic process | GO:0009127 | 0,046571057 | 1,331883907 | 28 | 6 |
| GO:BP | negative regulation of fatty acid metabolic process | GO:0045922 | 0,049648448 | 1,304094323 | 4 | 3 |
| GO:CC | cytoplasm | GO:0005737 | 3,02E-76 | 75,52014394 | 9524 | 509 |
| GO:CC | plastid | GO:0009536 | 8,93E-72 | 71,04914536 | 2856 | 269 |
| GO:CC | cytosol | GO:0005829 | 4,58E-65 | 64,33877588 | 2420 | 238 |
| GO:CC | intracellular anatomical structure | GO:0005622 | 5,29E-60 | 59,27639559 | 13505 | 576 |
| GO:CC | chloroplast | GO:0009507 | 7,70E-60 | 59,11333744 | 2351 | 227 |
| GO:CC | plastid stroma | GO:0009532 | 9,64E-57 | 56,01572331 | 769 | 128 |
| GO:CC | chloroplast stroma | GO:0009570 | 2,51E-56 | 55,59946345 | 762 | 127 |
| GO:CC | intracellular membrane-bounded organelle | GO:0043231 | 1,58E-36 | 35,80207077 | 11808 | 506 |
| GO:CC | membrane-bounded organelle | GO:0043227 | 2,52E-36 | 35,59860905 | 11823 | 506 |
| GO:CC | intracellular organelle | GO:0043229 | 4,46E-33 | 32,35050321 | 12312 | 512 |
| GO:CC | organelle | GO:0043226 | 6,78E-33 | 32,16883901 | 12326 | 512 |
| GO:CC | plastid envelope | GO:0009526 | 3,18E-17 | 16,49716585 | 988 | 87 |
| GO:CC | thylakoid | GO:0009579 | 7,34E-16 | 15,13430208 | 596 | 63 |
| GO:CC | organelle envelope | GO:0031967 | 2,18E-15 | 14,66217994 | 1451 | 106 |
| GO:CC | chloroplast envelope | GO:0009941 | 3,45E-12 | 11,46266112 | 692 | 62 |
| GO:CC | cellular anatomical structure | GO:0110165 | 8,97E-11 | 10,04741613 | 18498 | 601 |
| GO:CC | chloroplast thylakoid | GO:0009534 | 9,75E-10 | 9,011144542 | 517 | 48 |
| GO:CC | plastid thylakoid | GO:0031976 | 1,12E-09 | 8,951371932 | 519 | 48 |
| GO:CC | photosynthetic membrane | GO:0034357 | 3,88E-08 | 7,410662222 | 443 | 41 |

| | | | | | | |
|---|---|---|---|---|---|---|
| GO:CC | thylakoid membrane | GO:0042651 | 1,25E-07 | 6,90299782 | 442 | 40 |
| GO:CC | chloroplast thylakoid membrane | GO:0009535 | 1,28E-07 | 6,892356004 | 424 | 39 |
| GO:CC | plastid thylakoid membrane | GO:0055035 | 1,47E-07 | 6,833577396 | 426 | 39 |
| GO:CC | outer membrane | GO:0019867 | 6,59E-07 | 6,181376751 | 546 | 44 |
| GO:CC | organelle outer membrane | GO:0031968 | 6,59E-07 | 6,181376751 | 546 | 44 |
| GO:CC | mitochondrion | GO:0005739 | 1,07844E-06 | 5,967205457 | 1720 | 95 |
| GO:CC | spliceosomal complex | GO:0005681 | 2,13685E-06 | 5,67022695 | 130 | 19 |
| GO:CC | nuclear speck | GO:0016607 | 2,75889E-05 | 4,559266003 | 83 | 14 |
| GO:CC | plastid membrane | GO:0042170 | 4,30347E-05 | 4,366181358 | 674 | 46 |
| GO:CC | cytoplasmic side of endoplasmic reticulum membrane | GO:0098554 | 0,000598528 | 3,222915799 | 5 | 4 |
| GO:CC | thylakoid lumen | GO:0031977 | 0,001998524 | 2,699290713 | 74 | 11 |
| GO:CC | nuclear body | GO:0016604 | 0,002260377 | 2,645819171 | 119 | 14 |
| GO:CC | apoplast | GO:0048046 | 0,002630933 | 2,579890208 | 489 | 33 |
| GO:CC | endoplasmic reticulum tubular network | GO:0071782 | 0,003989917 | 2,399036139 | 7 | 4 |
| GO:CC | photosystem | GO:0009521 | 0,005484825 | 2,260837252 | 97 | 12 |
| GO:CC | protein-containing complex | GO:0032991 | 0,007224396 | 2,141198457 | 2604 | 113 |
| GO:CC | secretory vesicle | GO:0099503 | 0,009406675 | 2,026563858 | 170 | 16 |
| GO:CC | plastoglobule | GO:0010287 | 0,01461109 | 1,83531738 | 62 | 9 |
| GO:CC | photosystem II | GO:0009523 | 0,017800489 | 1,749568071 | 78 | 10 |
| GO:CC | plastid thylakoid lumen | GO:0031978 | 0,020303928 | 1,692419946 | 51 | 8 |
| GO:CC | chloroplast thylakoid lumen | GO:0009543 | 0,020303928 | 1,692419946 | 51 | 8 |
| GO:CC | intracellular membraneless organelle | GO:0043232 | 0,024358978 | 1,613340936 | 2568 | 109 |
| GO:CC | membraneless organelle | GO:0043228 | 0,024358978 | 1,613340936 | 2568 | 109 |
| GO:CC | vacuole | GO:0005773 | 0,025744319 | 1,589318588 | 1094 | 55 |
| GO:CC | membrane protein complex | GO:0098796 | 0,046549609 | 1,332083965 | 452 | 28 |
| KEGG | Metabolic pathways | KEGG:01100 | 1,02E-16 | 15,98999378 | 2337 | 214 |
| KEGG | Biosynthesis of amino acids | KEGG:01230 | 1,28E-12 | 11,89318629 | 242 | 48 |
| KEGG | Carbon metabolism | KEGG:01200 | 6,96E-12 | 11,15770204 | 271 | 50 |
| KEGG | Biosynthesis of secondary metabolites | KEGG:01110 | 1,27E-09 | 8,895242546 | 1283 | 128 |
| KEGG | Pyruvate metabolism | KEGG:00620 | 1,19406E-06 | 5,922974751 | 96 | 22 |
| KEGG | Carbon fixation in photosynthetic organisms | KEGG:00710 | 2,13441E-06 | 5,67072158 | 68 | 18 |
| KEGG | Glycolysis / Gluconeogenesis | KEGG:00010 | 3,83241E-06 | 5,416528233 | 119 | 24 |
| KEGG | 2-Oxocarboxylic acid metabolism | KEGG:01210 | 8,94954E-05 | 4,048199272 | 103 | 20 |
| KEGG | Arginine biosynthesis | KEGG:00220 | 0,001427539 | 2,845412053 | 36 | 10 |
| KEGG | Histidine metabolism | KEGG:00340 | 0,003106431 | 2,50773825 | 19 | 7 |
| KEGG | Glyoxylate and dicarboxylate metabolism | KEGG:00630 | 0,006735476 | 2,171631732 | 77 | 14 |
| KEGG | Alanine, aspartate and glutamate metabolism | KEGG:00250 | 0,00721844 | 2,141556666 | 51 | 11 |
| KEGG | Glutathione metabolism | KEGG:00480 | 0,015565539 | 1,807835824 | 103 | 16 |
| KEGG | C5-Branched dibasic acid metabolism | KEGG:00660 | 0,017246295 | 1,763304181 | 7 | 4 |
| KEGG | Spliceosome | KEGG:03040 | 0,019976165 | 1,699487893 | 205 | 25 |
| KEGG | Fructose and mannose metabolism | KEGG:00051 | 0,049083772 | 1,309062072 | 63 | 11 |
| KEGG | Valine, leucine and isoleucine biosynthesis | KEGG:00290 | 0,049652206 | 1,304061455 | 21 | 6 |
| WP | Glycolysis | WP:WP2621 | 0,002925614 | 2,533783035 | 37 | 10 |
| WP | Glucosinolate biosynthesis from methionine | WP:WP4597 | 0,036664787 | 1,435750836 | 15 | 5 |

### **8** NMR – Spectra

(Sorted according to molecule numbering)

Compound 8: $^1H$ – NMR ($CDCl_3$, 400 MHz)

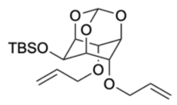


**8**

$C_{19}H_{32}O_6Si$
MW: 384,5440

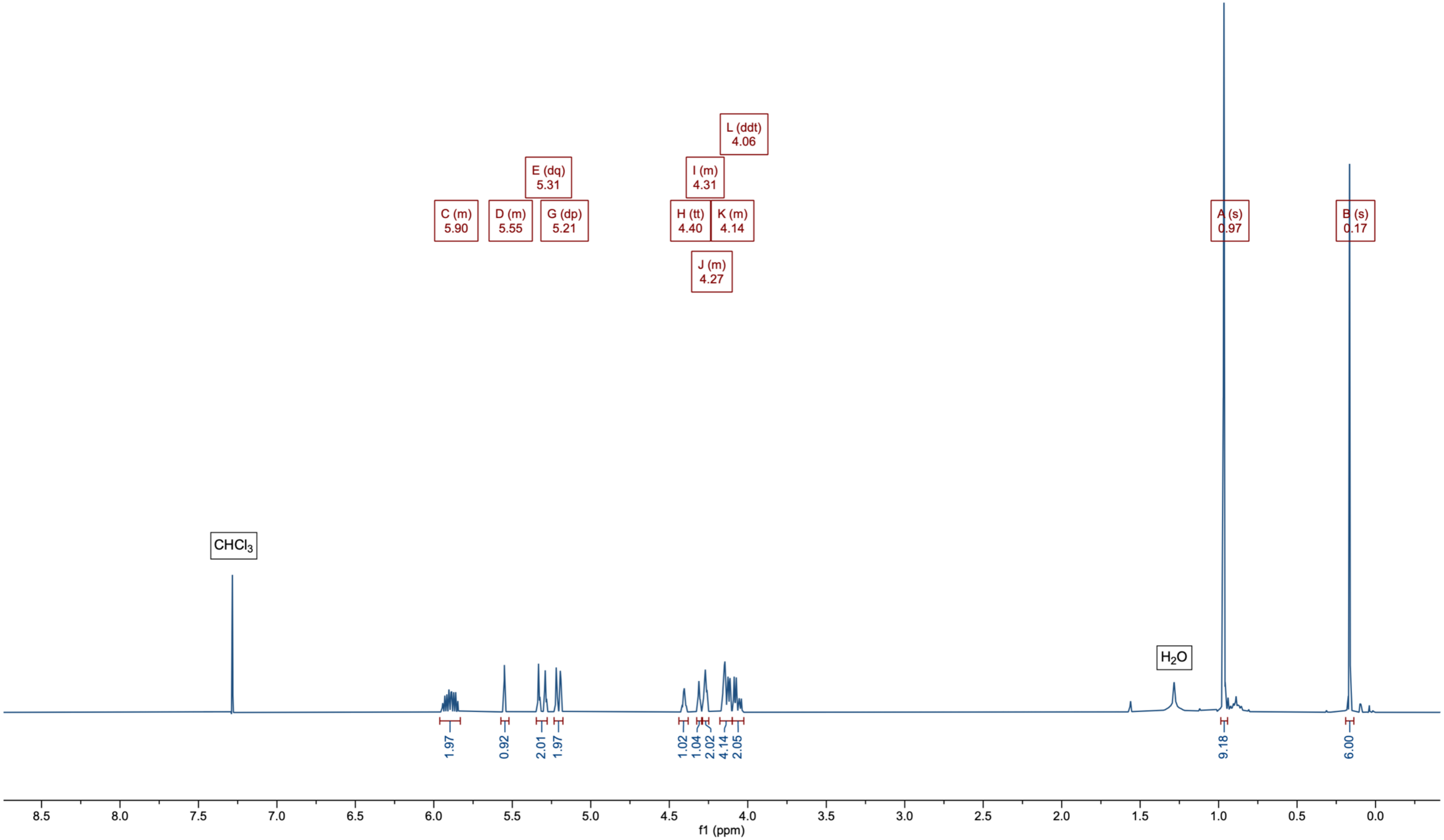

### Compound 8: $^{13}C\{^{1}H\}$ – NMR ($CDCl_3$, 101 MHz)

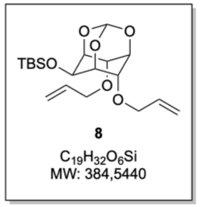


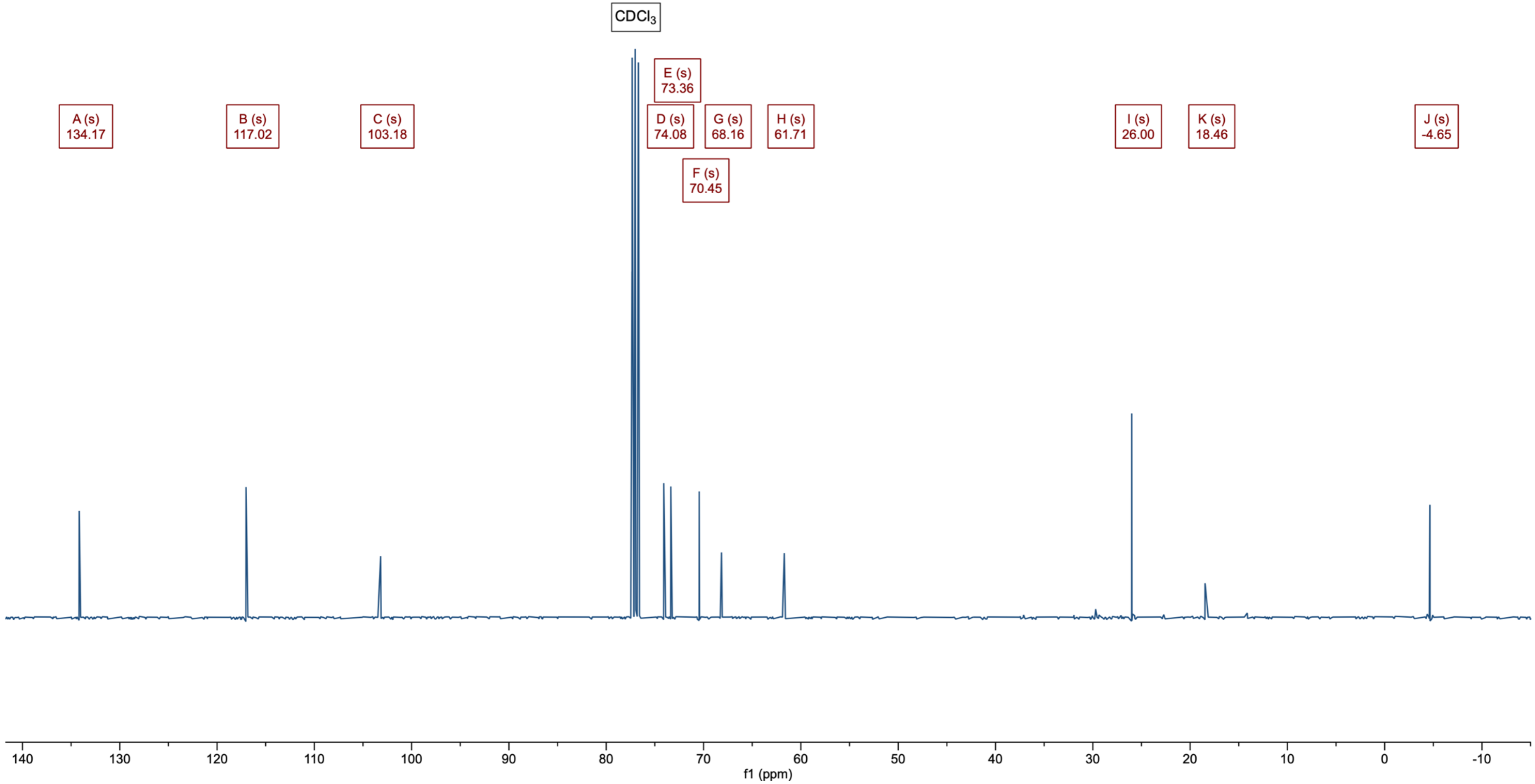

Compound 9: $^{1}H$ – NMR (MeCN-$d_3$, 400 MHz)

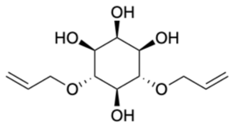

9

$C_{12}H_{20}O_6$
MW: 260,2860

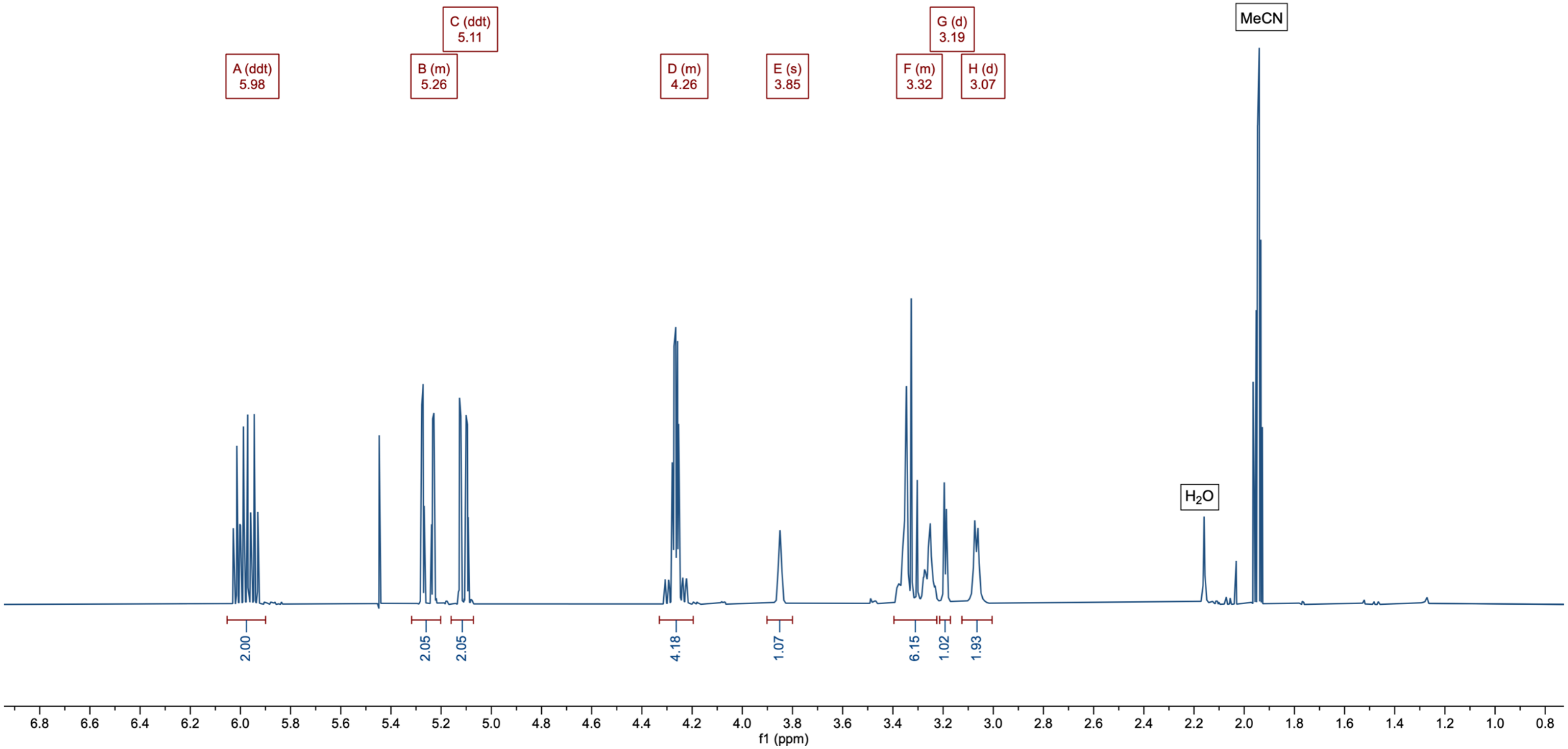

Compound 9: $^{13}C\{^{1}H\}$ – NMR (MeCN-$d_3$, 101 MHz)

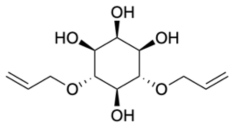

9

$C_{12}H_{20}O_6$
MW: 260,2860

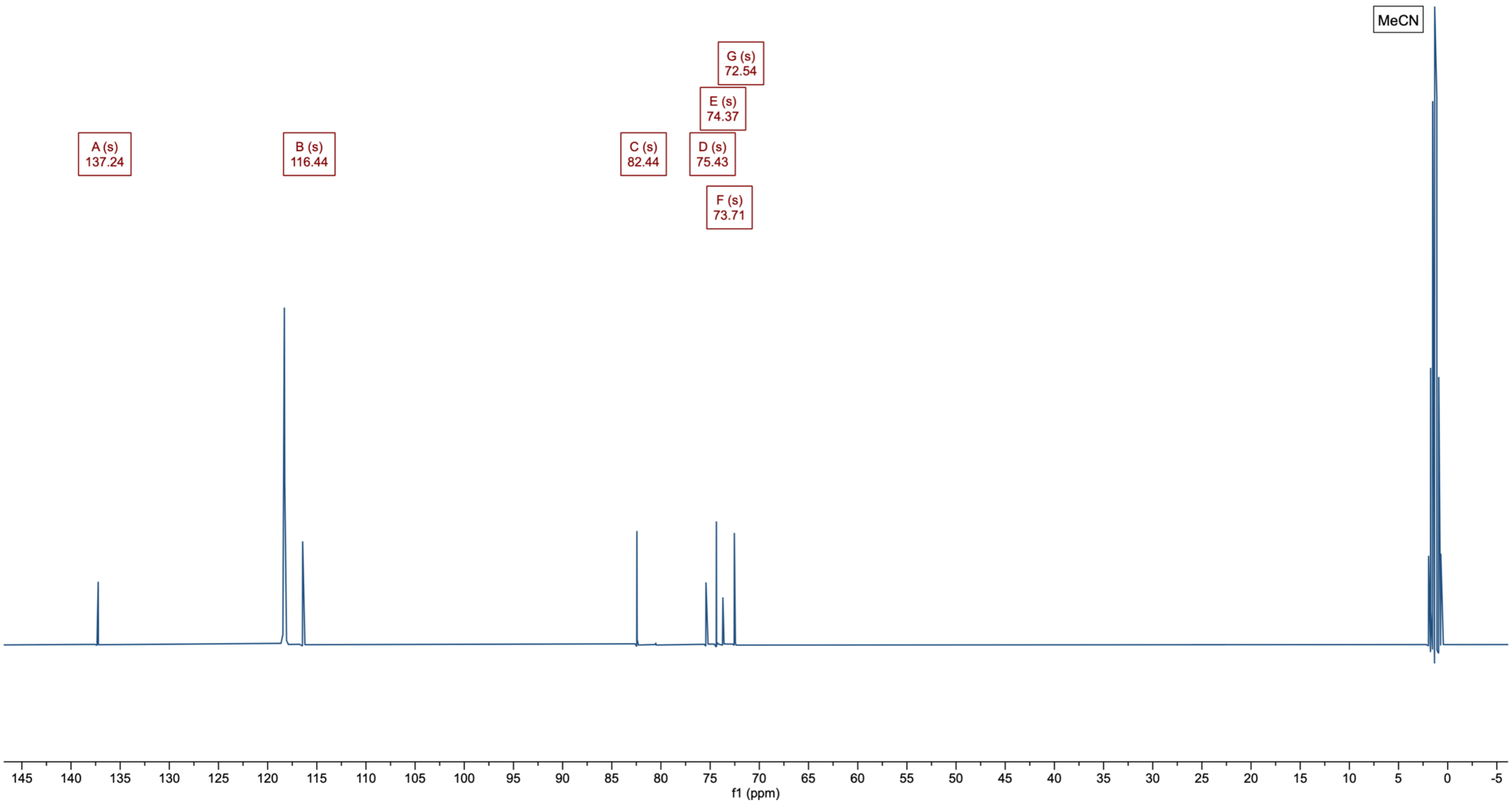

Compound 10: $^1H$ – NMR ($CDCl_3$, 400 MHz)

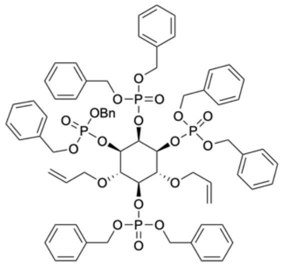


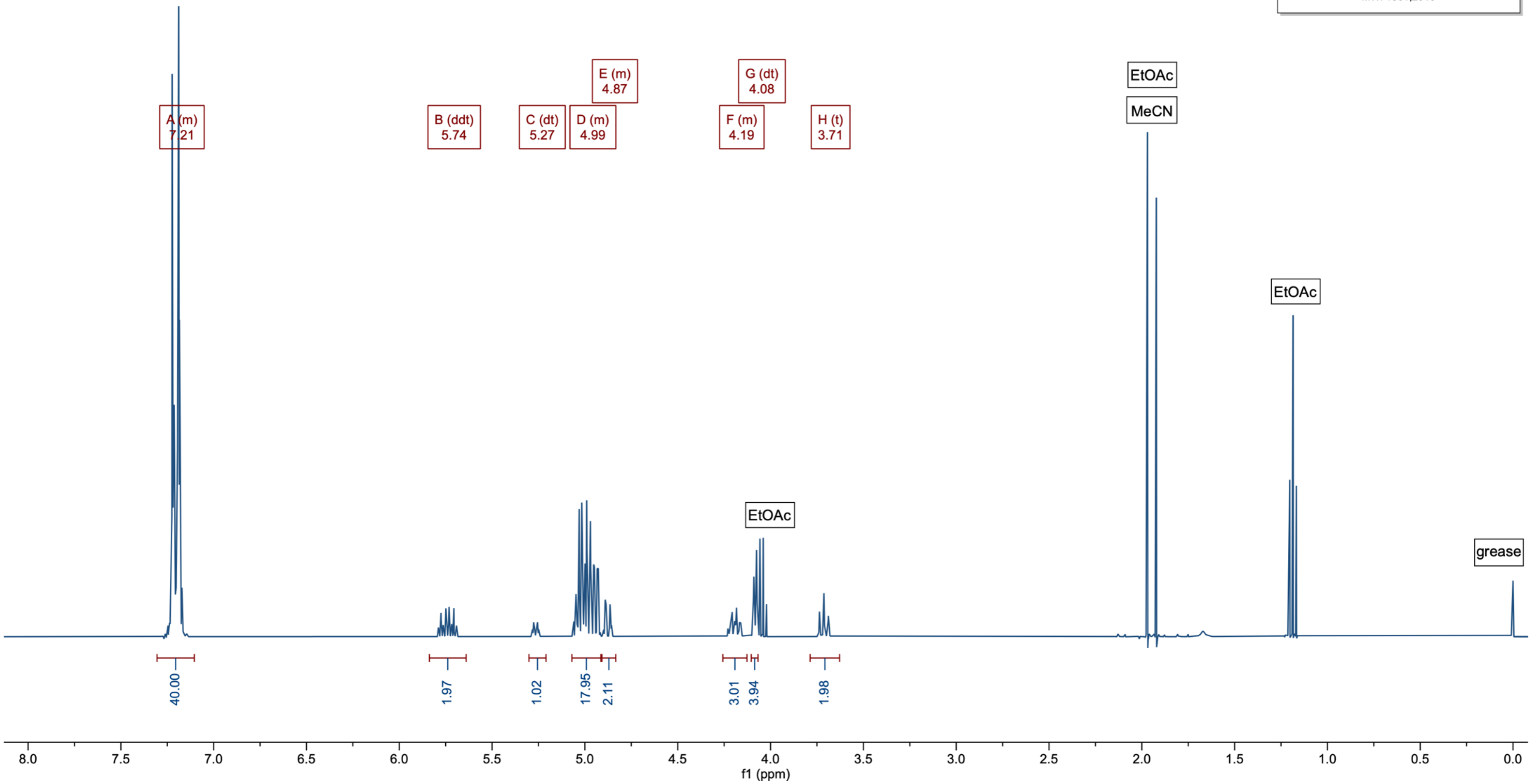

Compound 10: $^{31}P\{^{1}H\}$ – NMR ($CDCl_3$, 162 MHz)

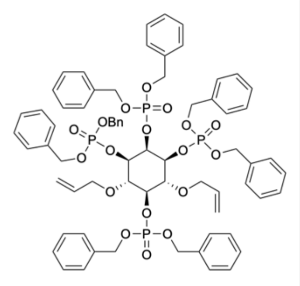


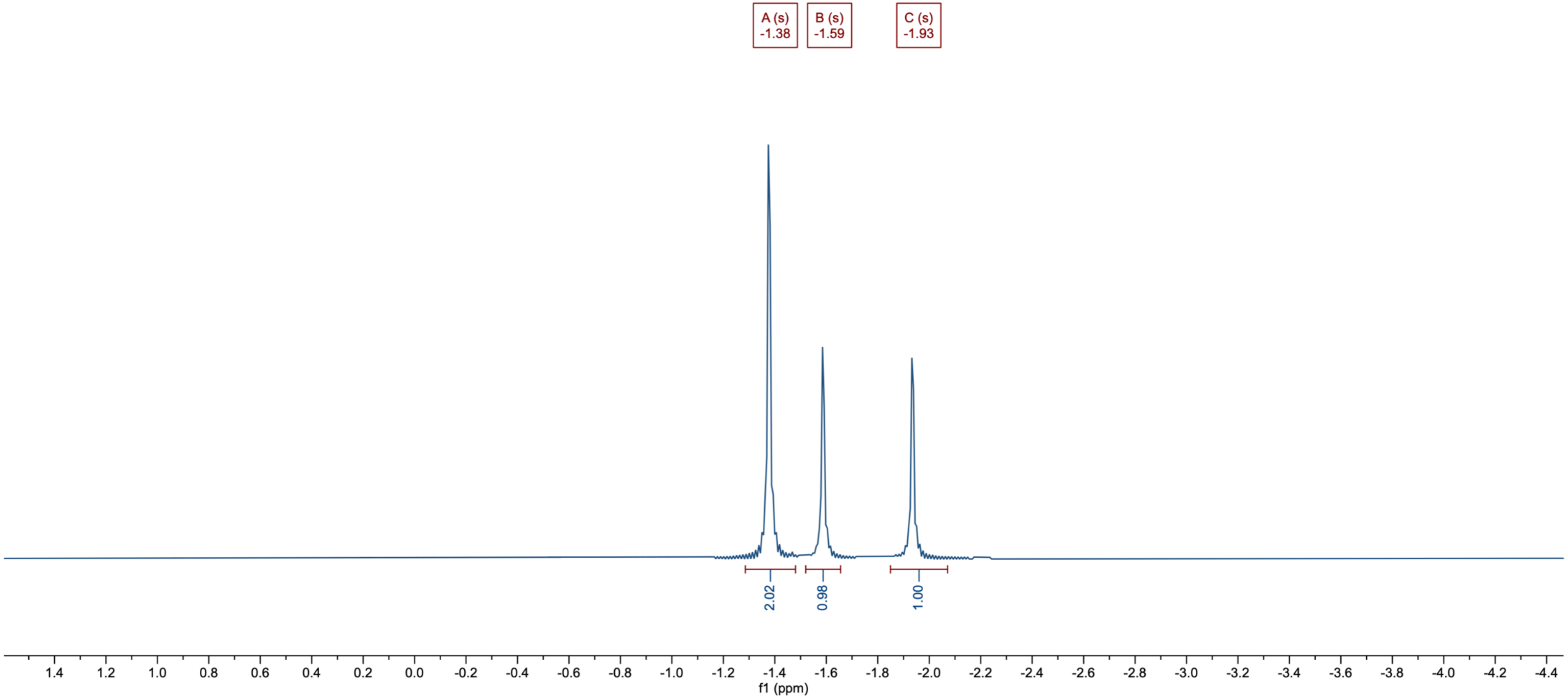

#### Compound 10: $^{13}C\{^{1}H\}$ – NMR ($CDCl_3$, 101 MHz)

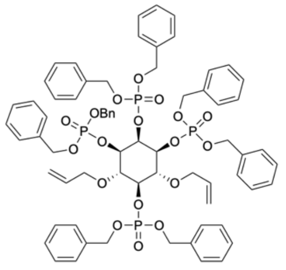

**10**

$C_{68}H_{72}O_{18}P_4$
MW: 1301,2010

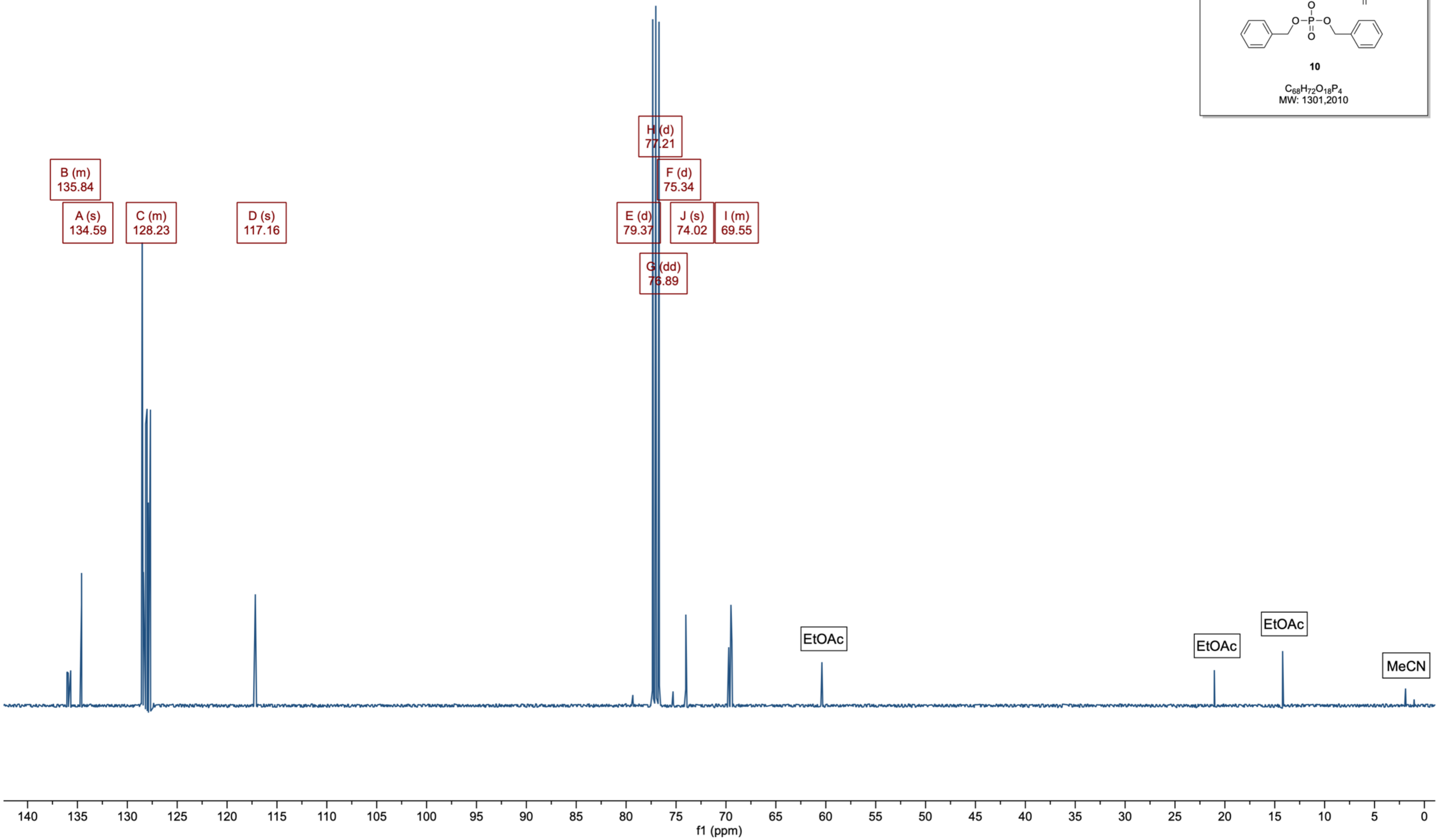

Compound 11: $^{1}H$ – NMR ($CDCl_3$, 400 MHz)

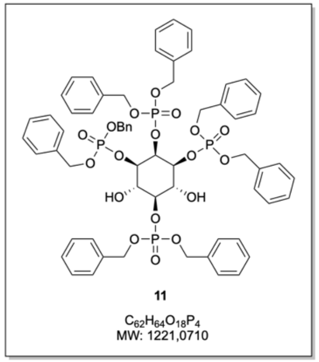


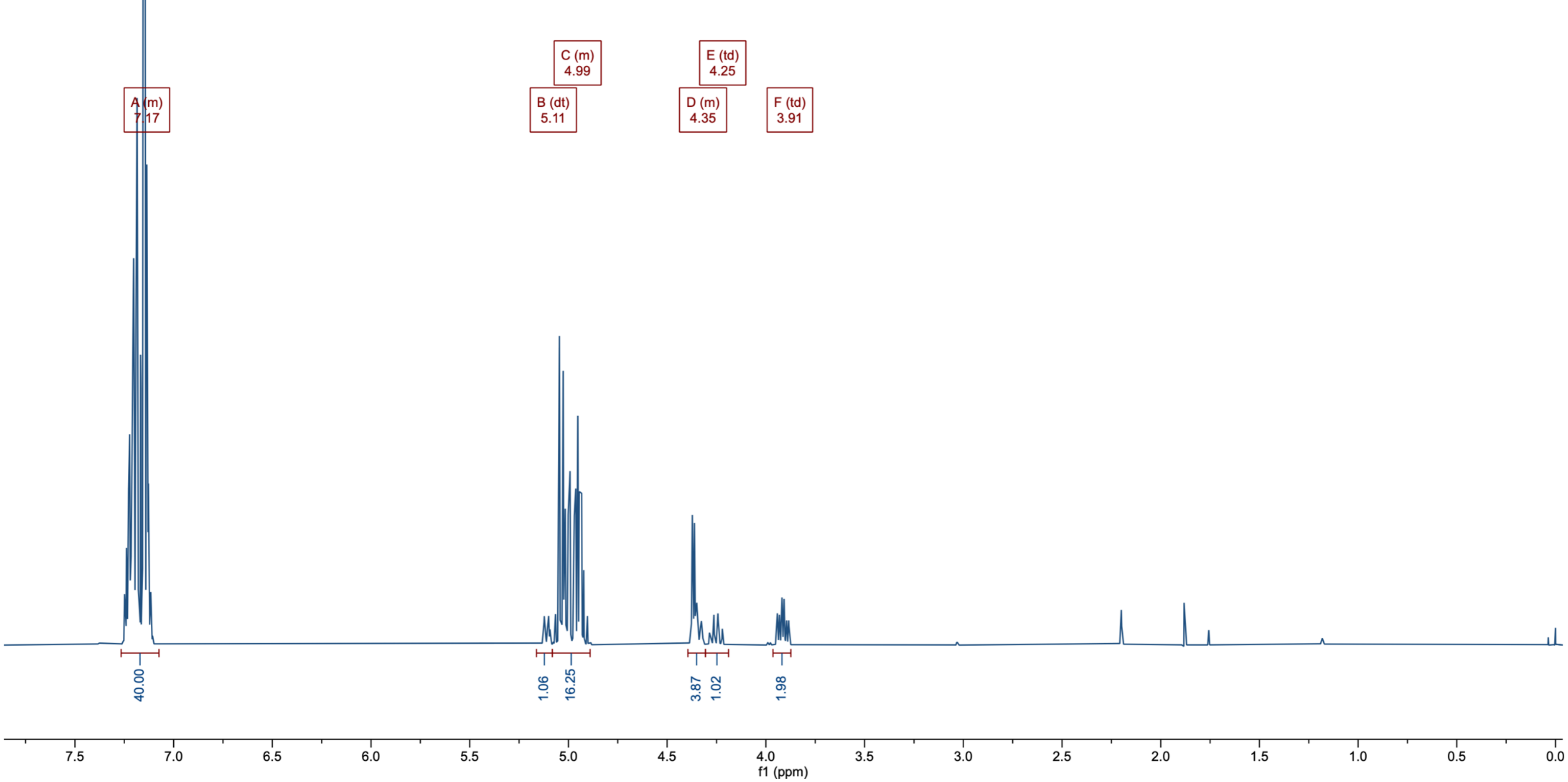

Compound 11: $^{31}P\{^{1}H\}$ – NMR ($CDCl_3$, 162 MHz)

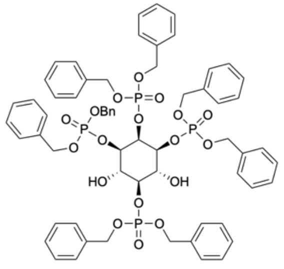


**11**

$C_{62}H_{64}O_{18}P_4$
MW: 1221,0710

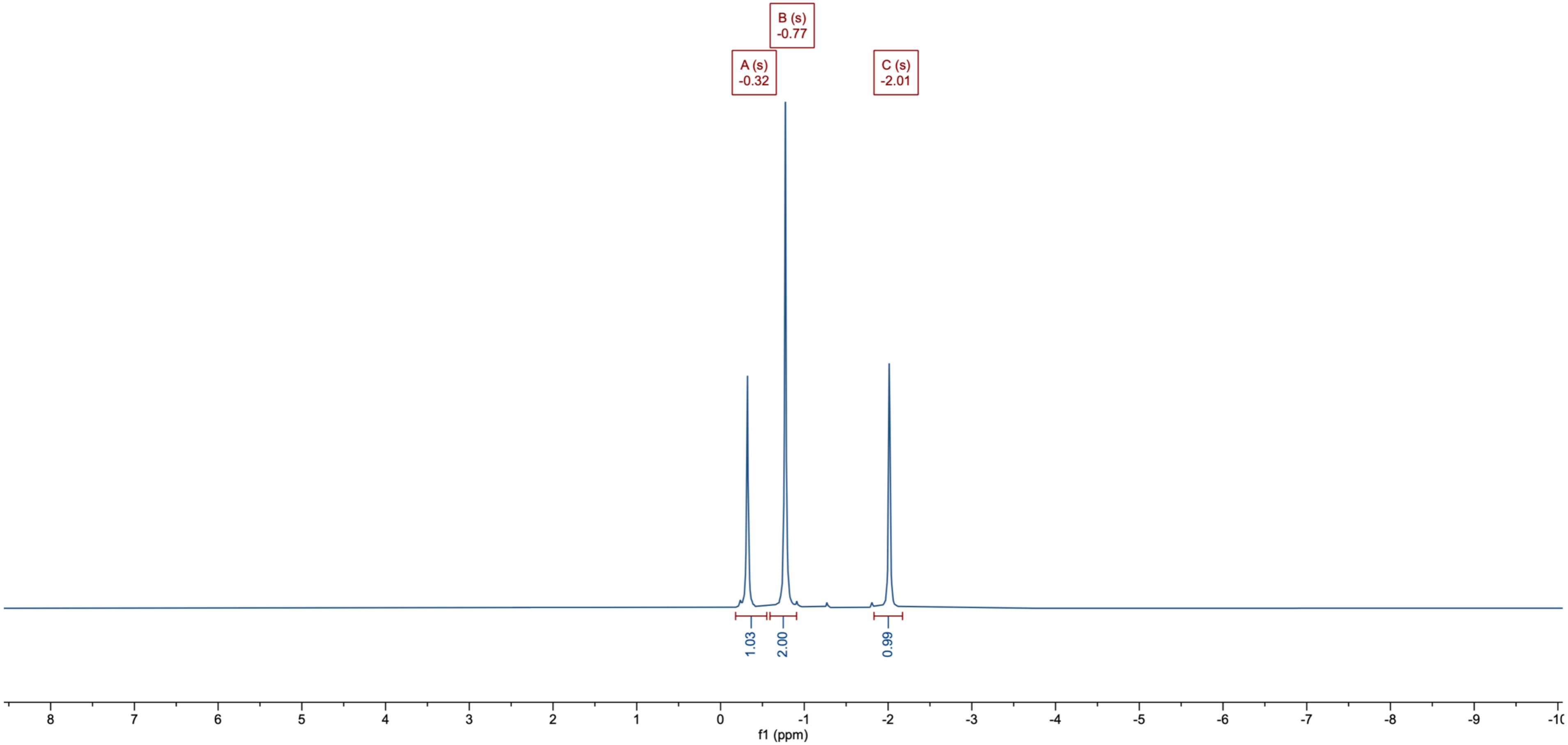

#### Compound 11: $^{13}C\{^1H\}$ – NMR ($CDCl_3$, 101 MHz)

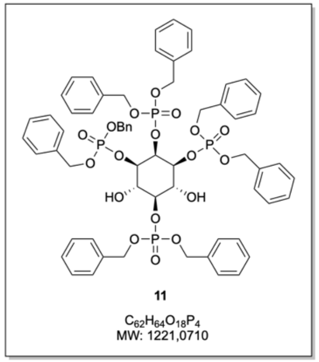

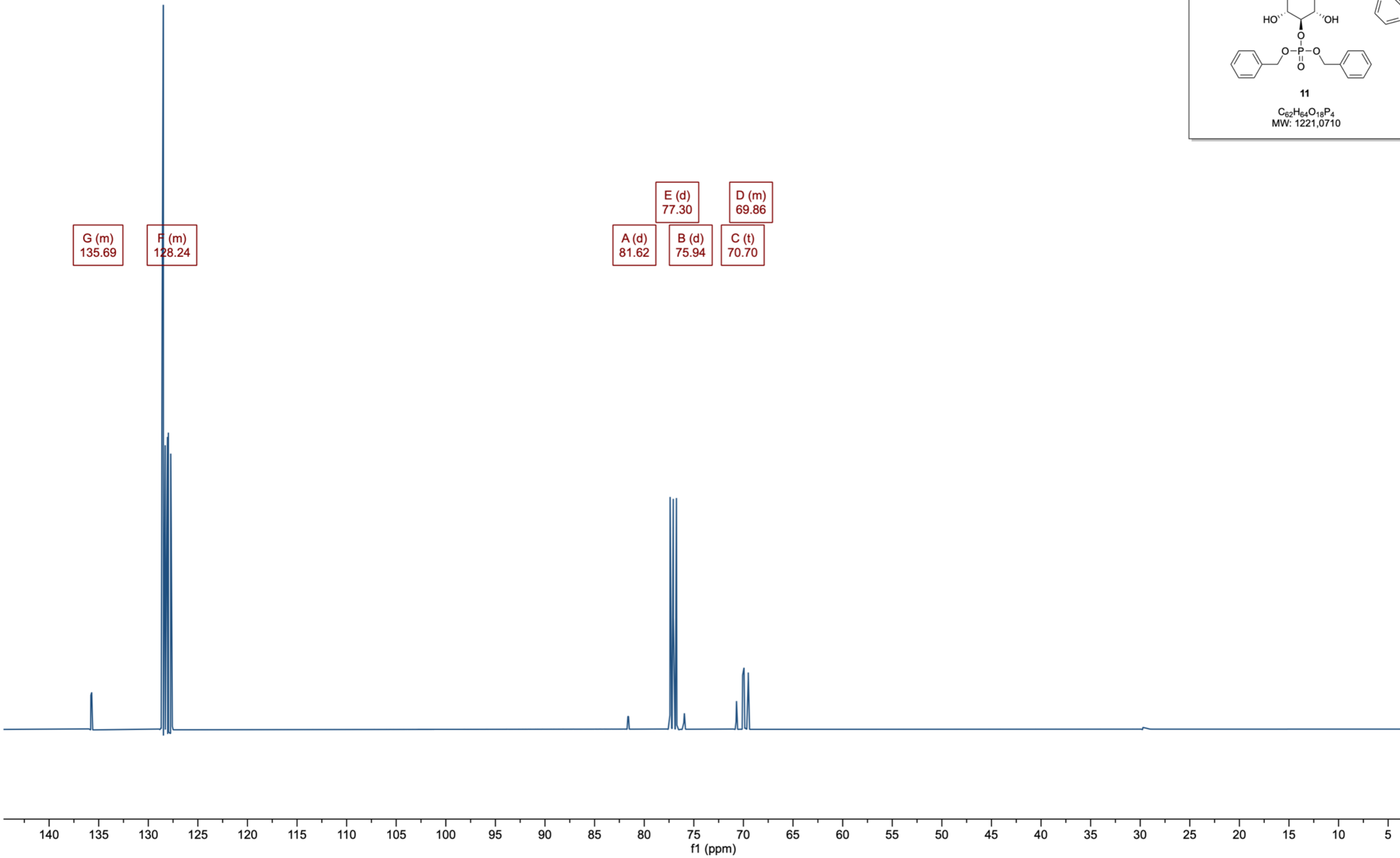

#### Compound 12: $^{1}H$ – NMR ($CDCl_3$, 400 MHz)

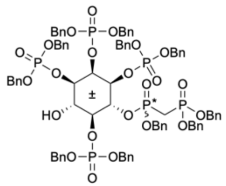

**12**

$C_{84}H_{86}O_{23}P_6$
MW: 1649,4316

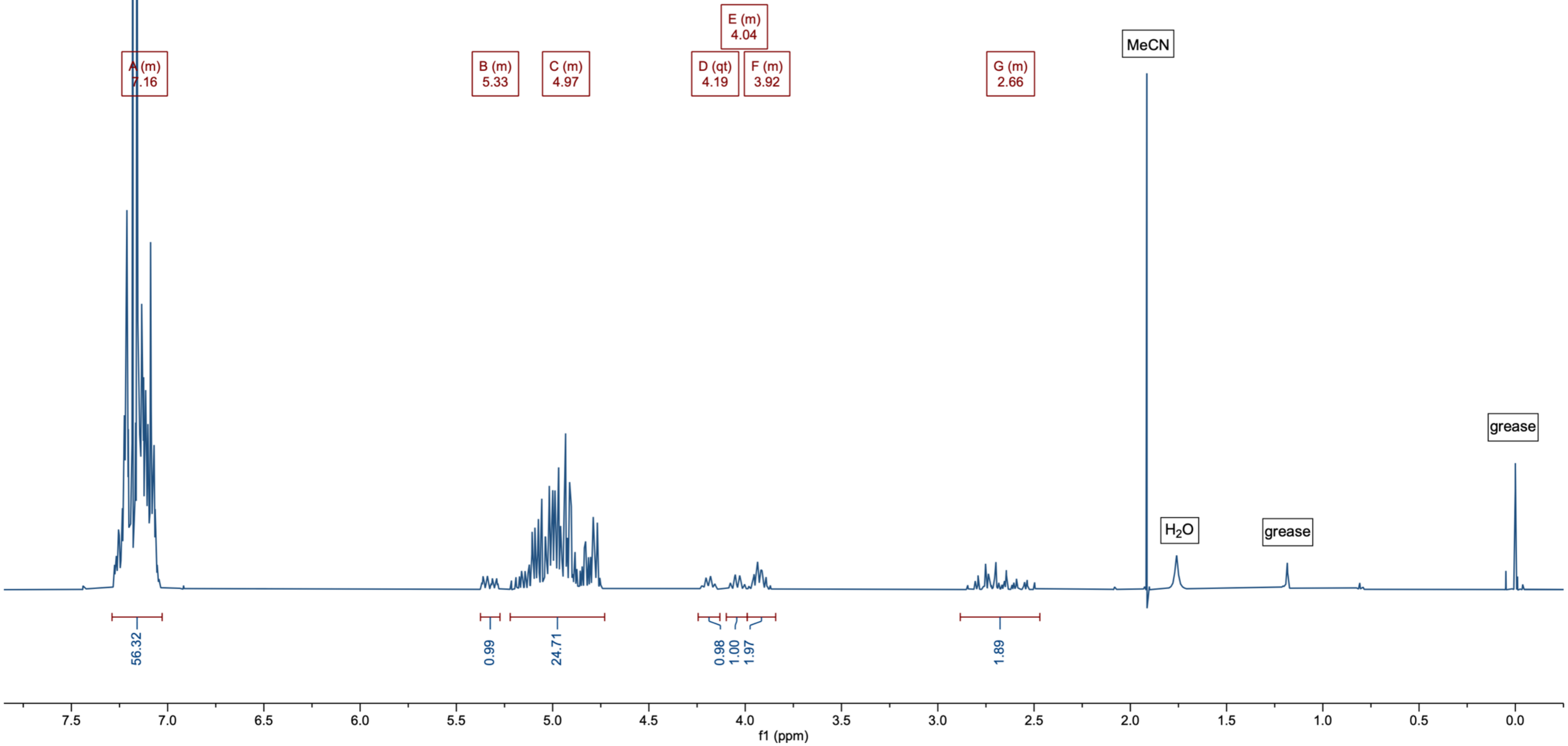

Compound 12: $^{31}P\{^{1}H\}$ – NMR ($CDCl_3$, 162 MHz)

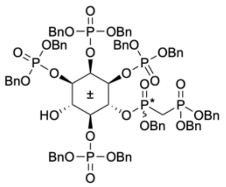

**12**

$C_{84}H_{86}O_{23}P_6$
MW: 1649,4316

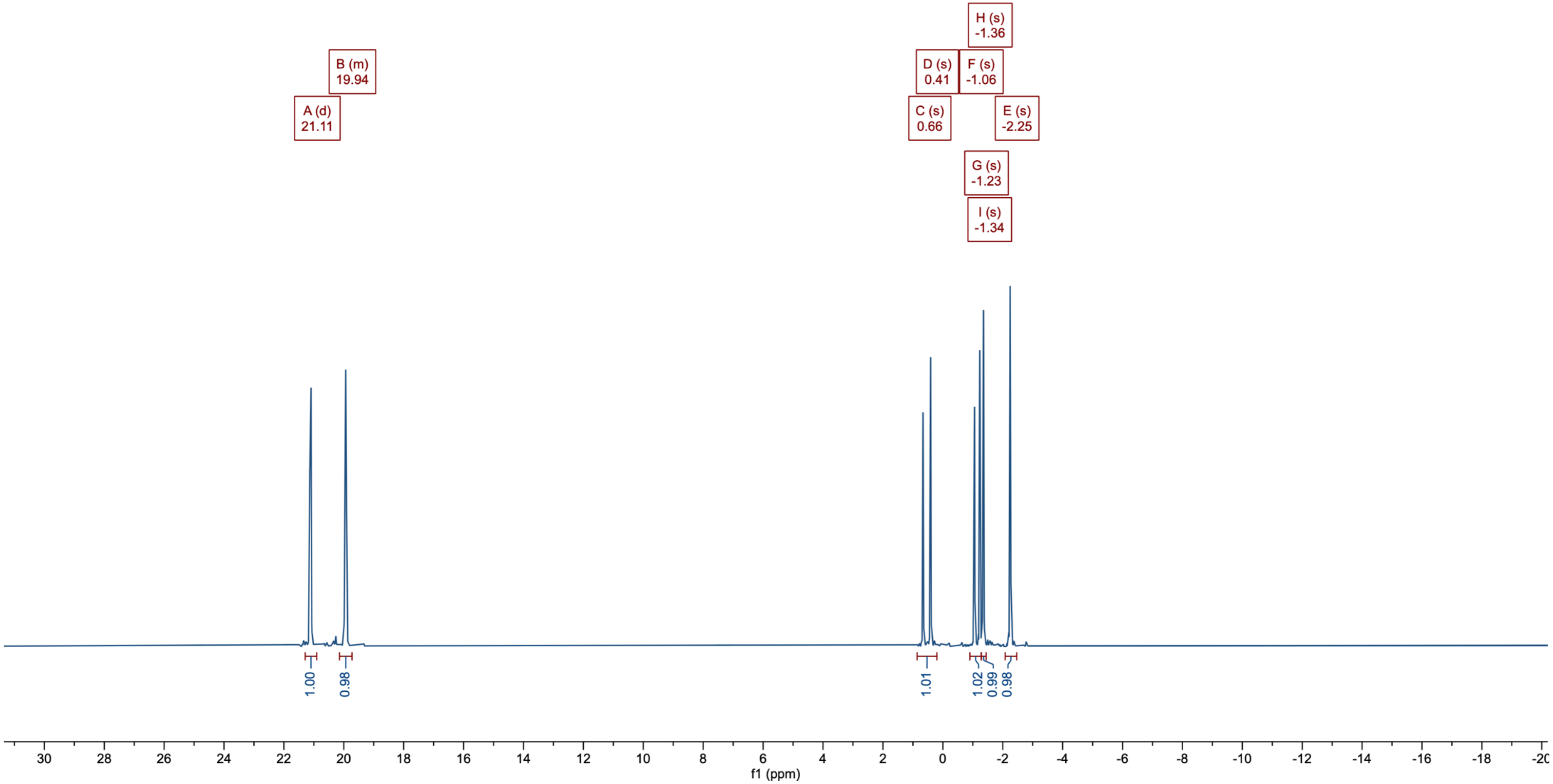

Compound 12: $^{13}C\{^{1}H\}$ – NMR ($CDCl_3$, 101 MHz)

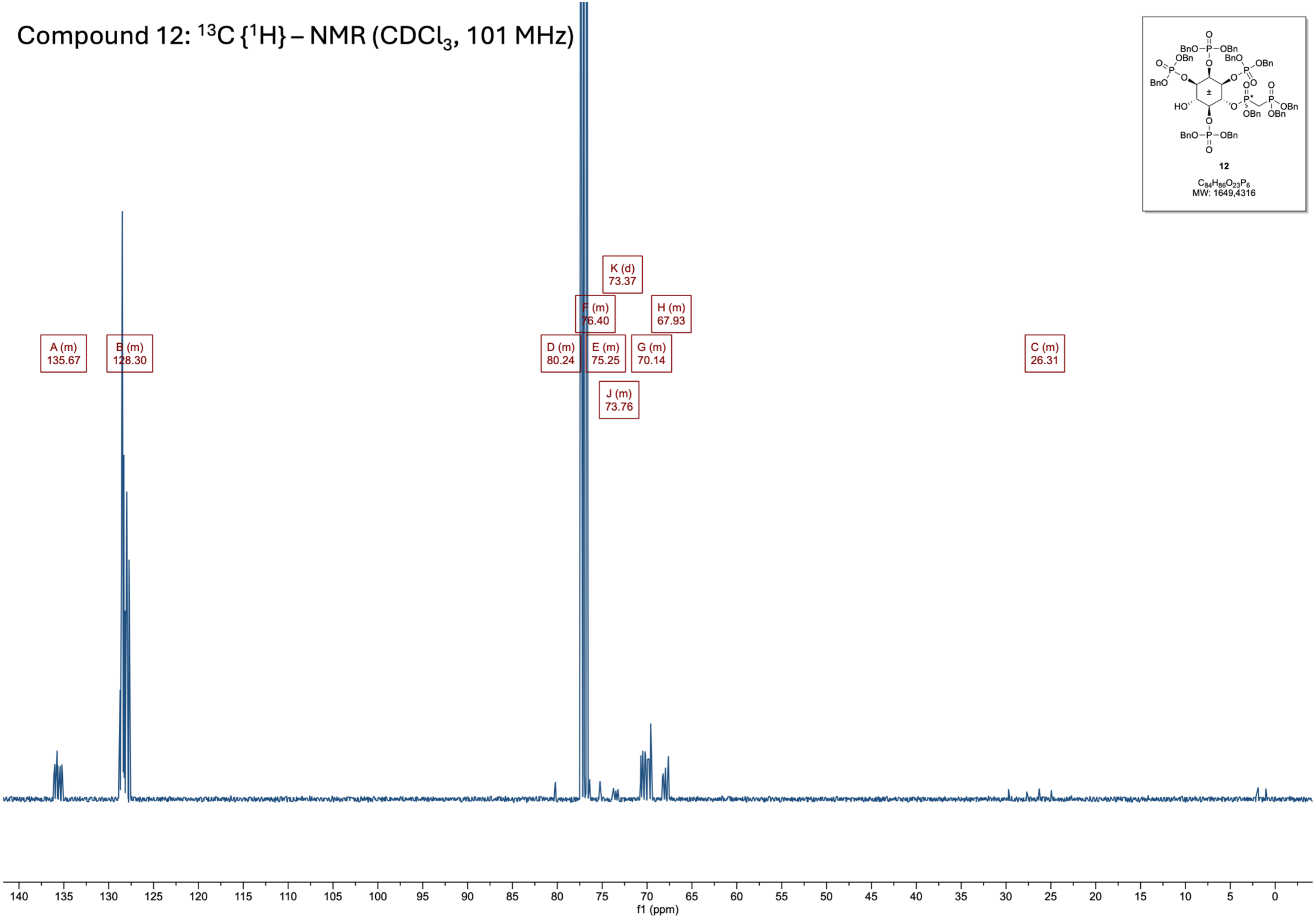


**12**

$C_{84}H_{86}O_{23}P_6$
MW: 1649,4316

### Compound 13: $^{1}H$ – NMR ($CDCl_3$, 400 MHz)

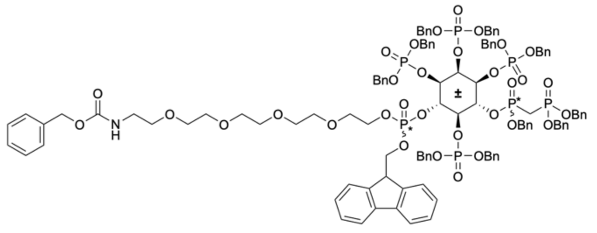

**13**

$C_{116}H_{124}NO_{32}P_7$
MW: 2261,0593

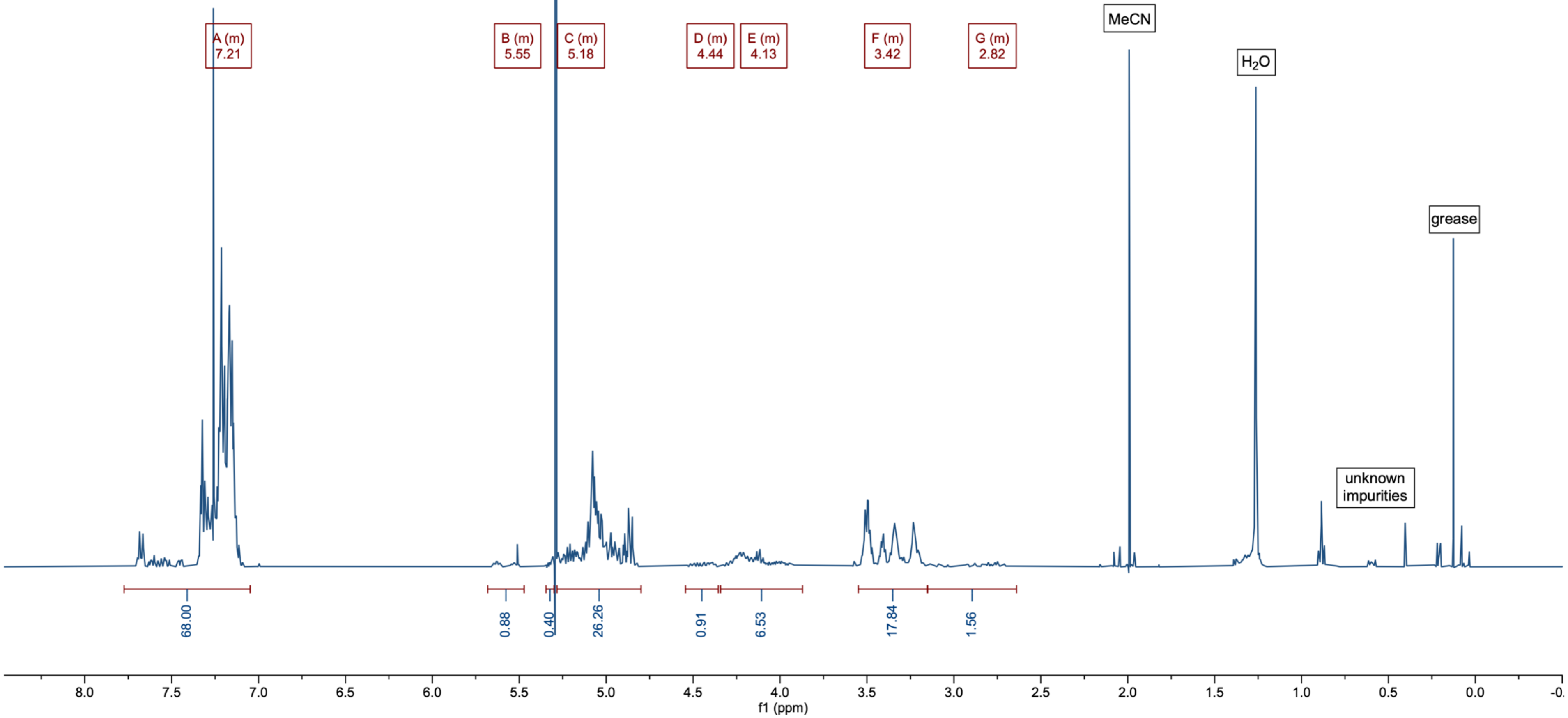

Compound 13: $^{31}P\{^{1}H\}$ – NMR ($CDCl_3$, 162 MHz)

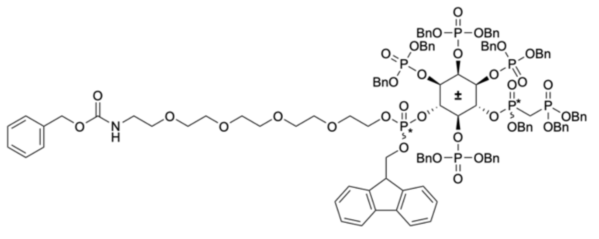

13

$C_{116}H_{124}NO_{32}P_7$
MW: 2261,0593

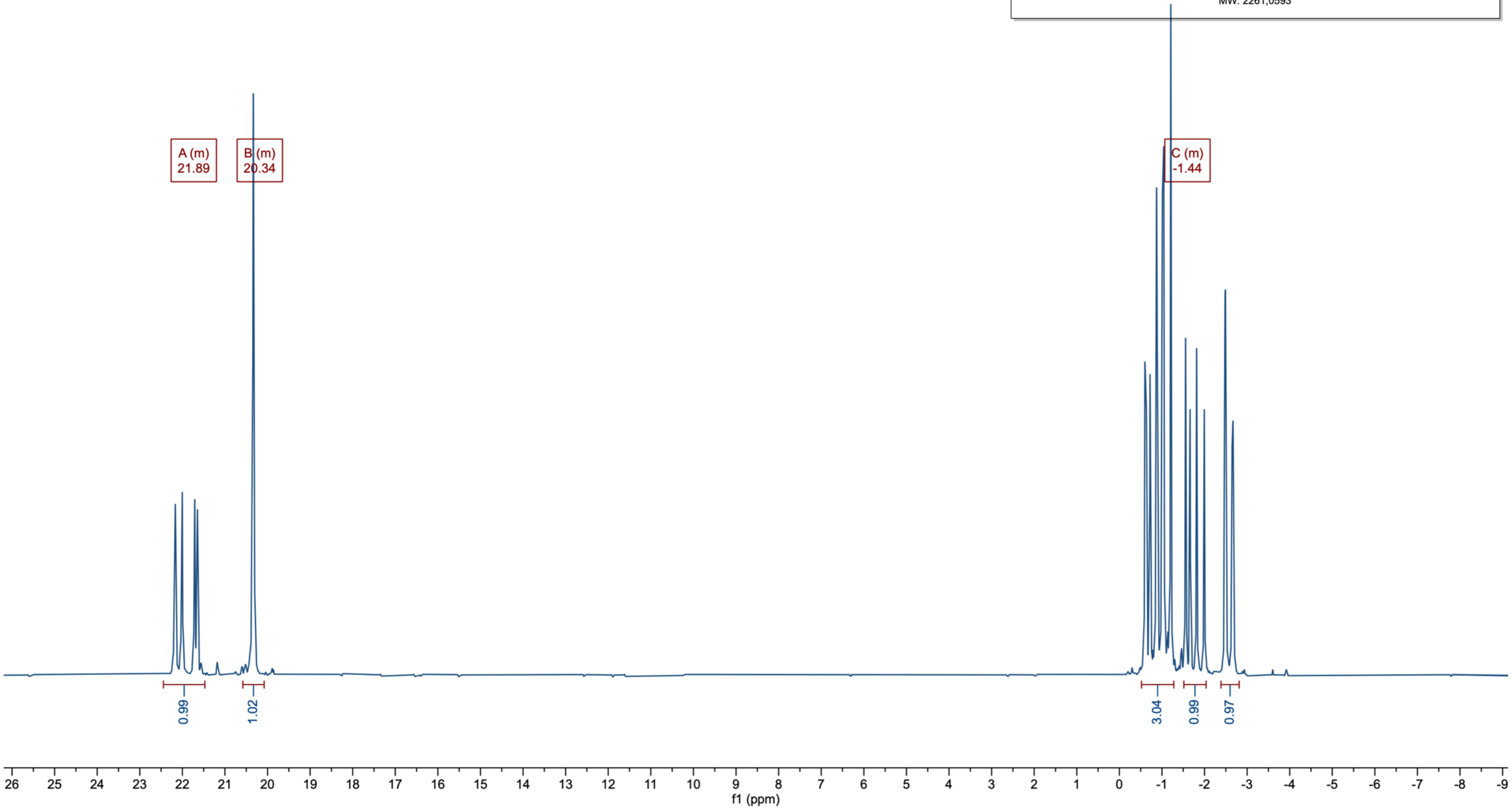

Compound 13: $^{13}C\{^{1}H\}$ – NMR ($CDCl_3$, 101 MHz)

13

$C_{116}H_{124}NO_{32}P_7$
MW: 2261,0593

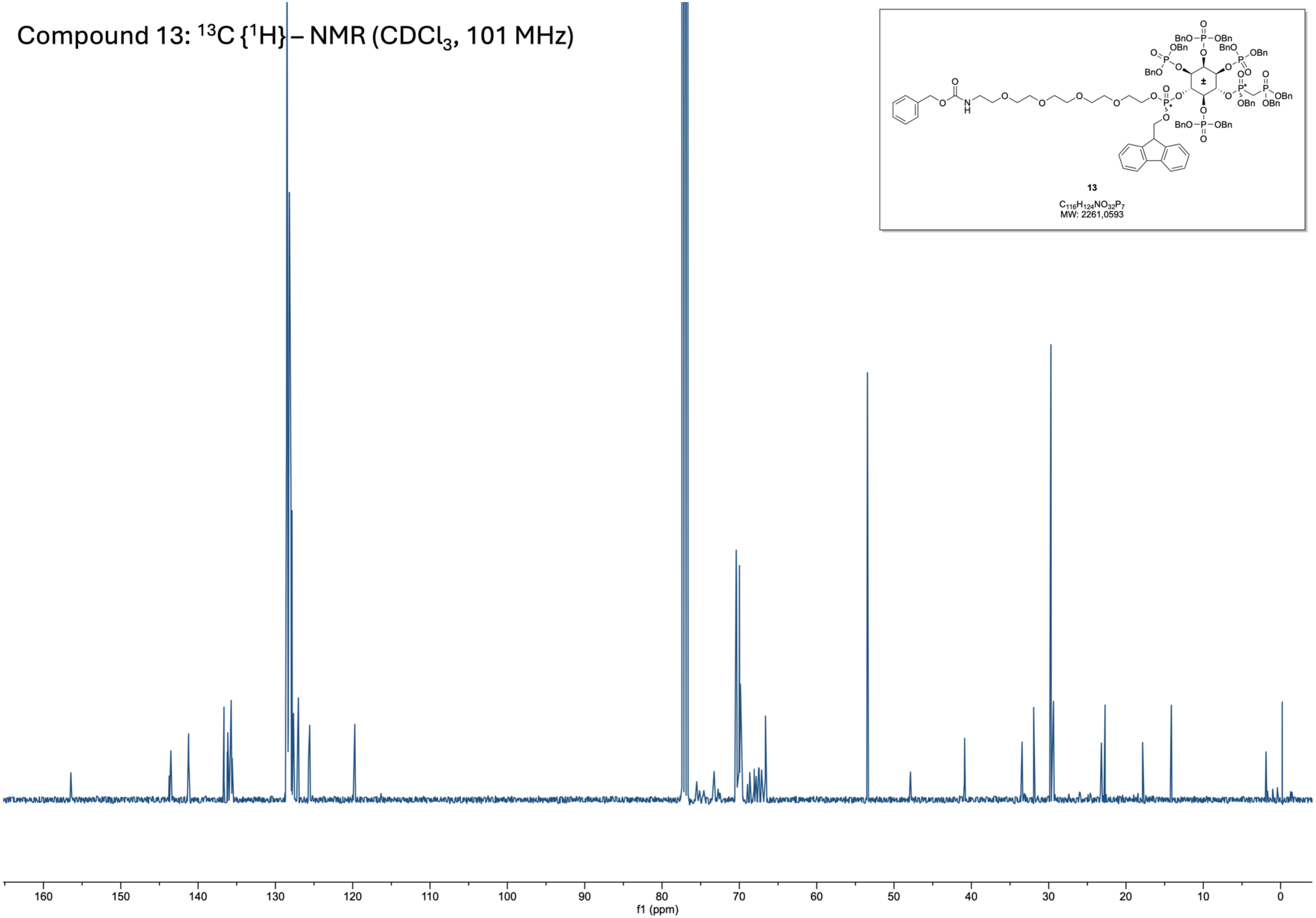

Compound 14: $^{1}H$ – NMR ($D_2O$, 400 MHz)

*Supplemented with $N(iPr)_2$ to enhance resolution*

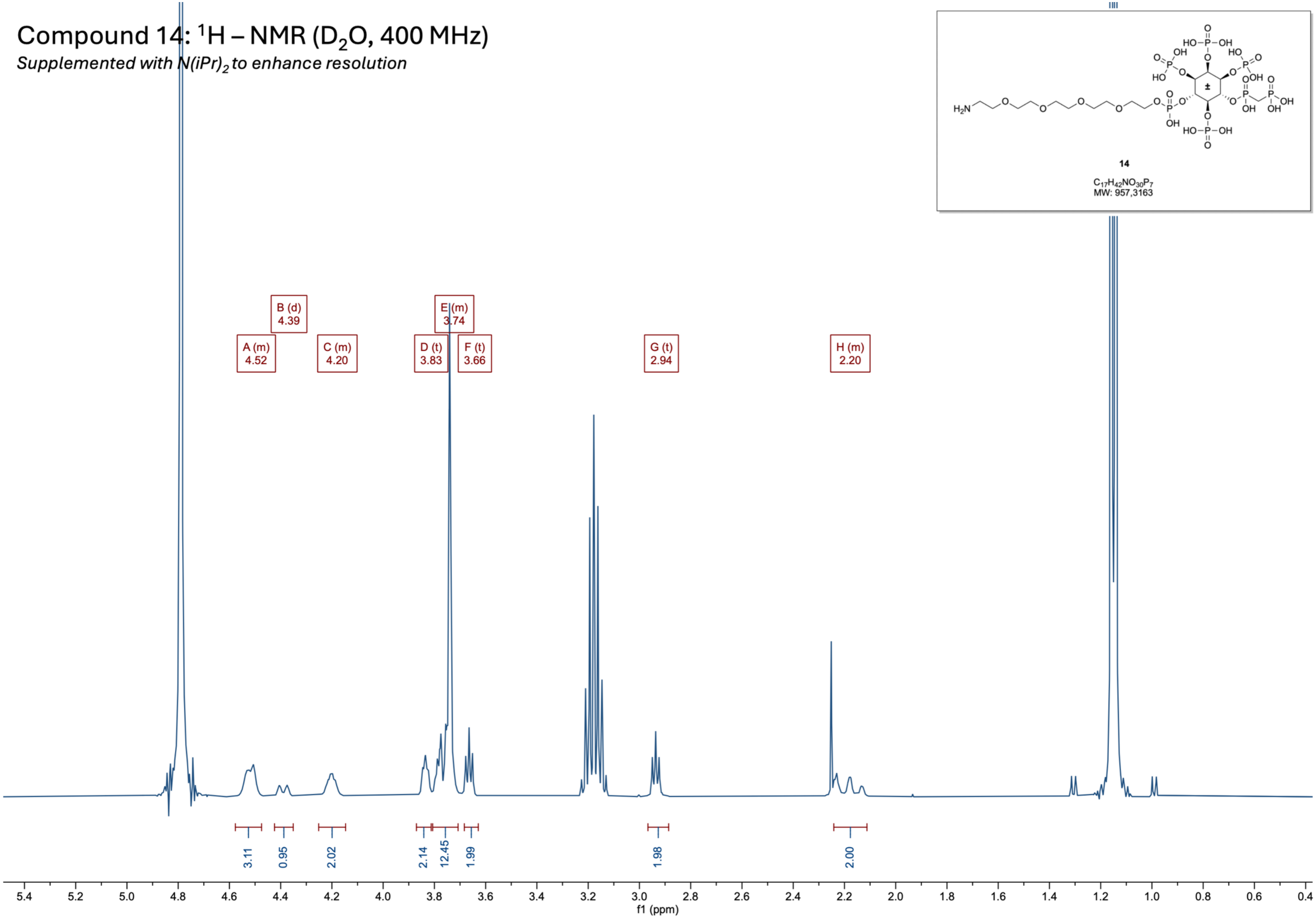

### Compound 14: $^{31}P\{^{1}H\}$ – NMR ($D_2O$, 162 MHz)

*Supplemented with $N(iPr)_2$ to enhance resolution*

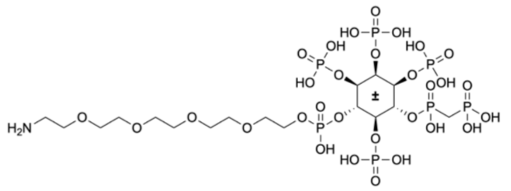

**14**

$C_{17}H_{42}NO_{30}P_7$
MW: 957,3163

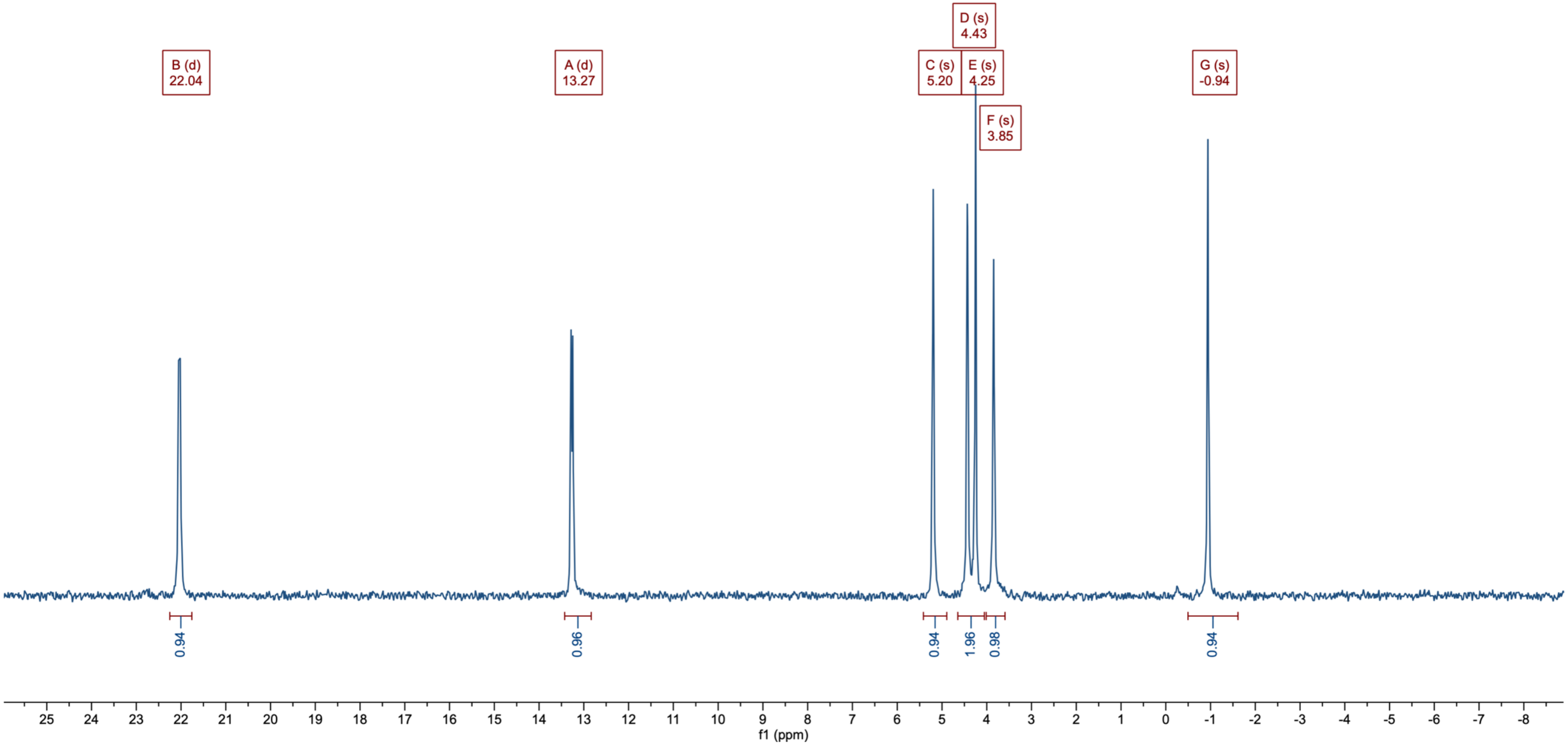

Compound 14: $^{13}C\{^1H\}$ – NMR ($D_2O$, 101 MHz)

*Supplemented with $N(iPr)_2$ to enhance resolution*

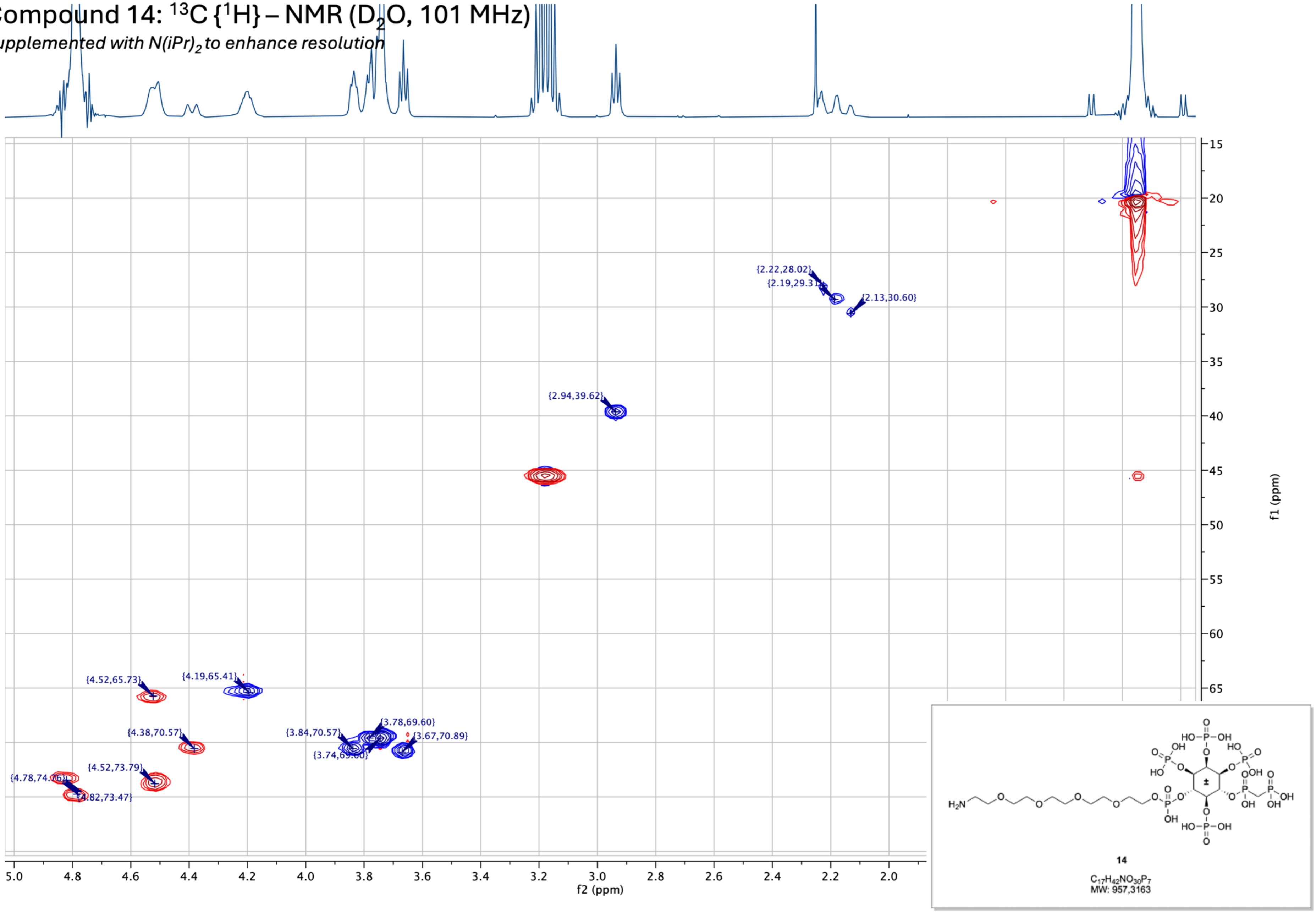


14

$C_{17}H_{42}NO_{30}P_7$
MW: 957,3163

Compound 16: $^{1}H$ – NMR ($CDCl_3$, 400 MHz)

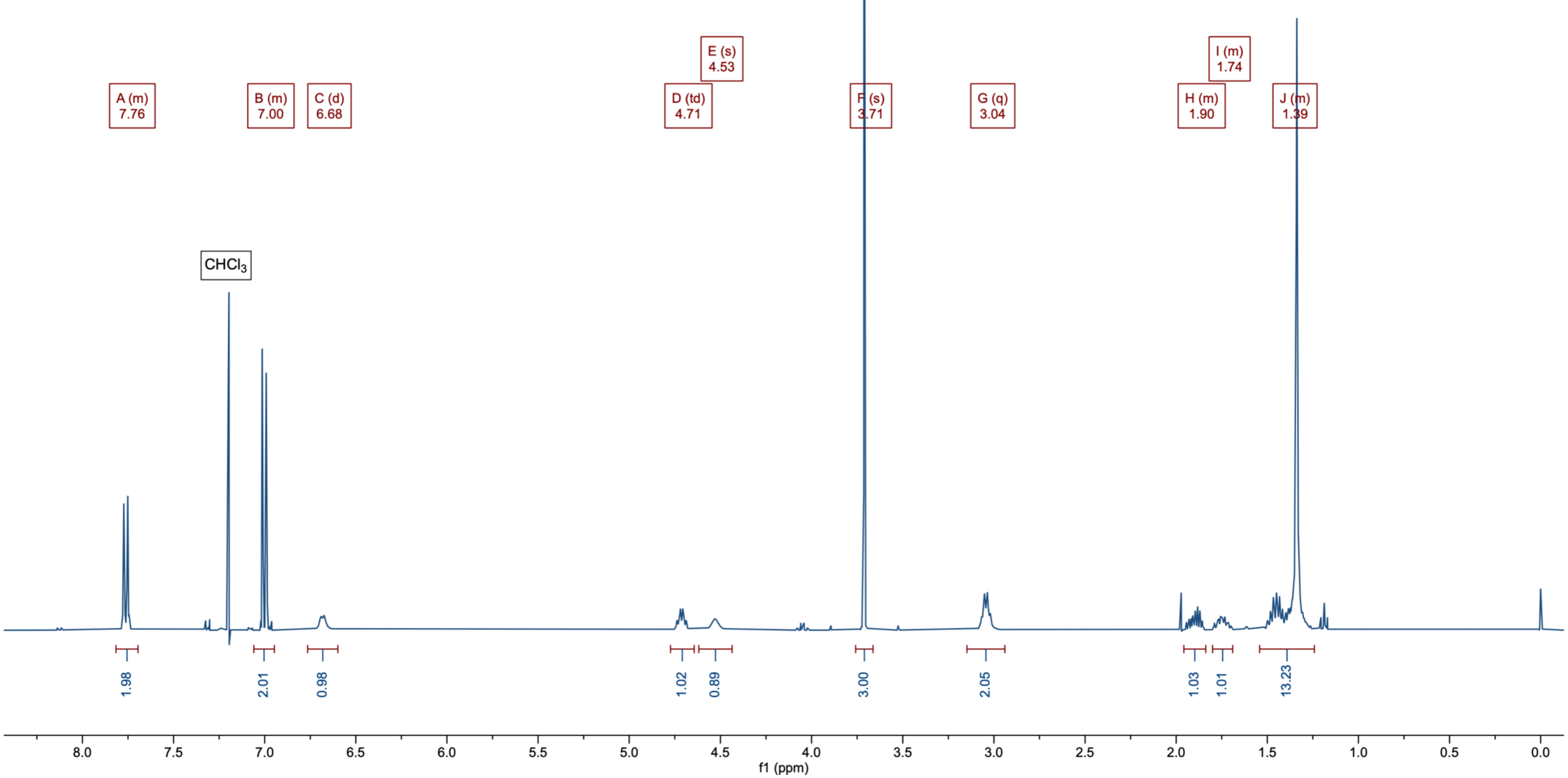


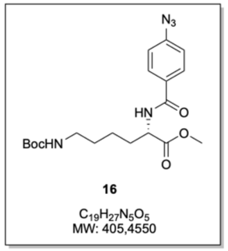

Compound 16: $^{13}C\{^{1}H\}$ – NMR ($CDCl_3$, 101 MHz)

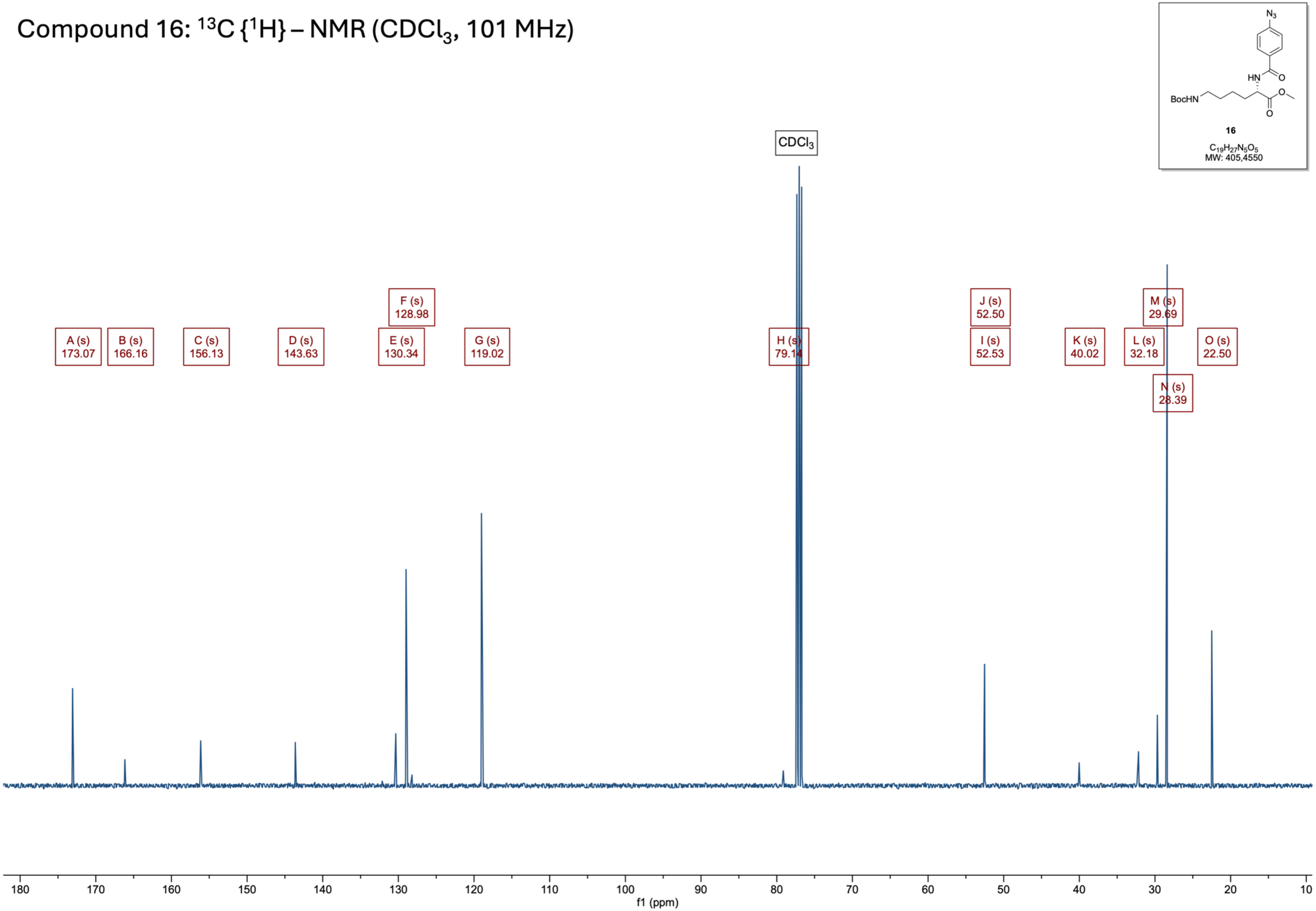

Compound 17: $^{1}H$ – NMR ($CDCl_3$, 400 MHz)

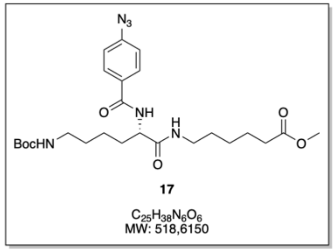


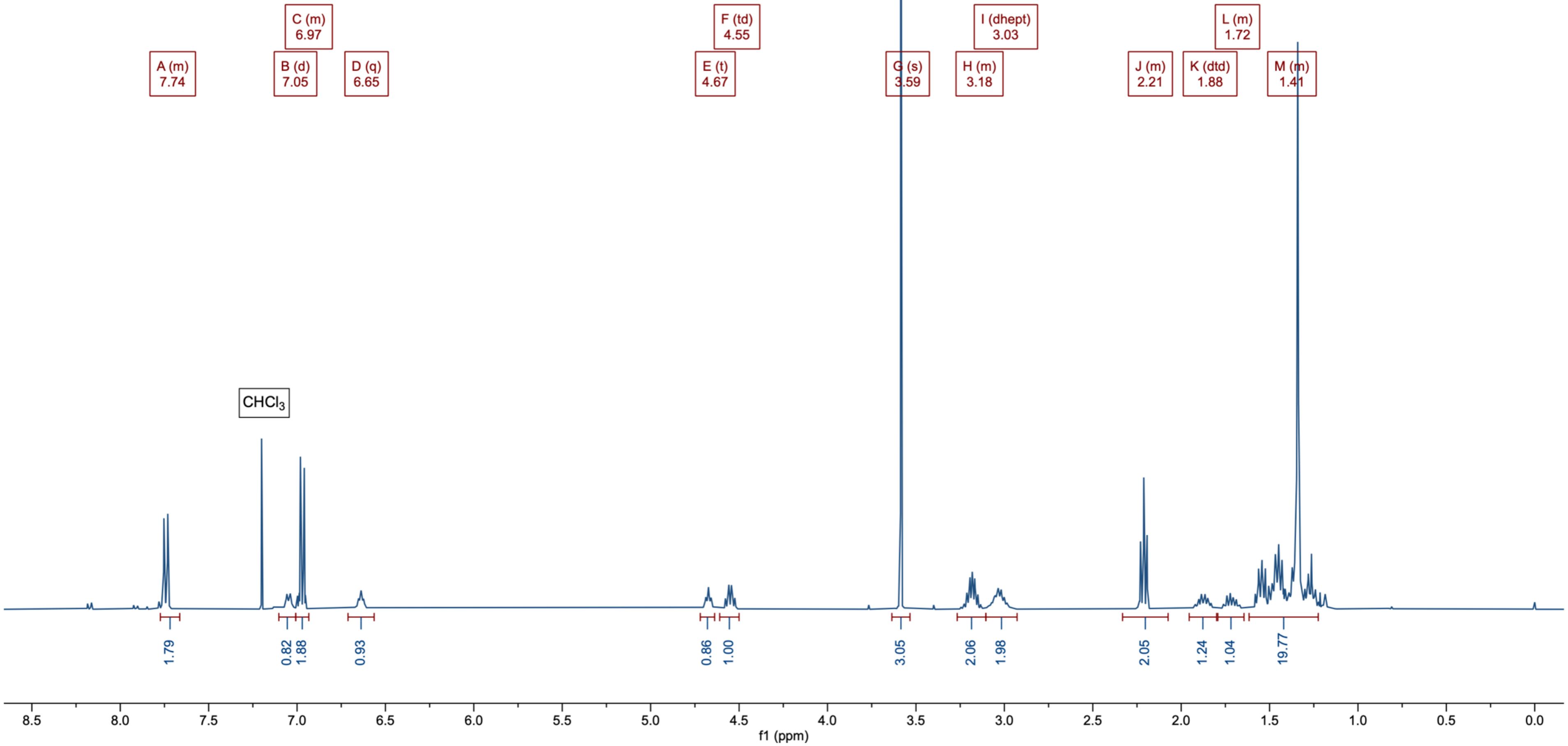

Compound 17: $^{13}C\{^{1}H\}$ – NMR ($CDCl_3$, 101 MHz)

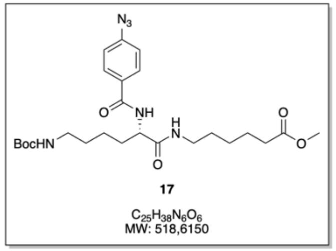


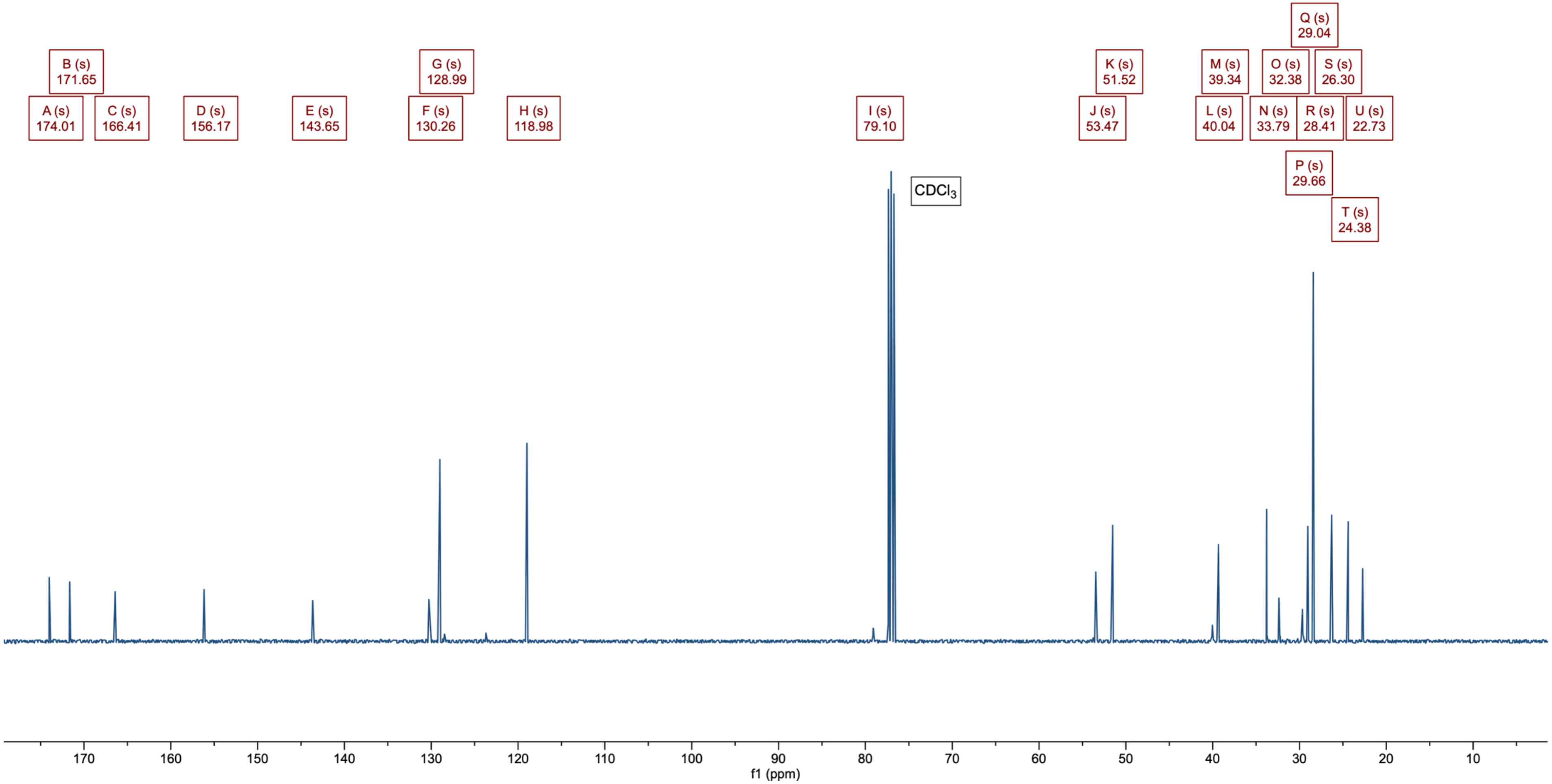

#### Compound 18: $^1H$ – NMR ($CDCl_3$, 400 MHz)

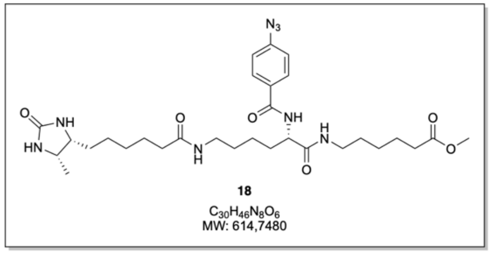


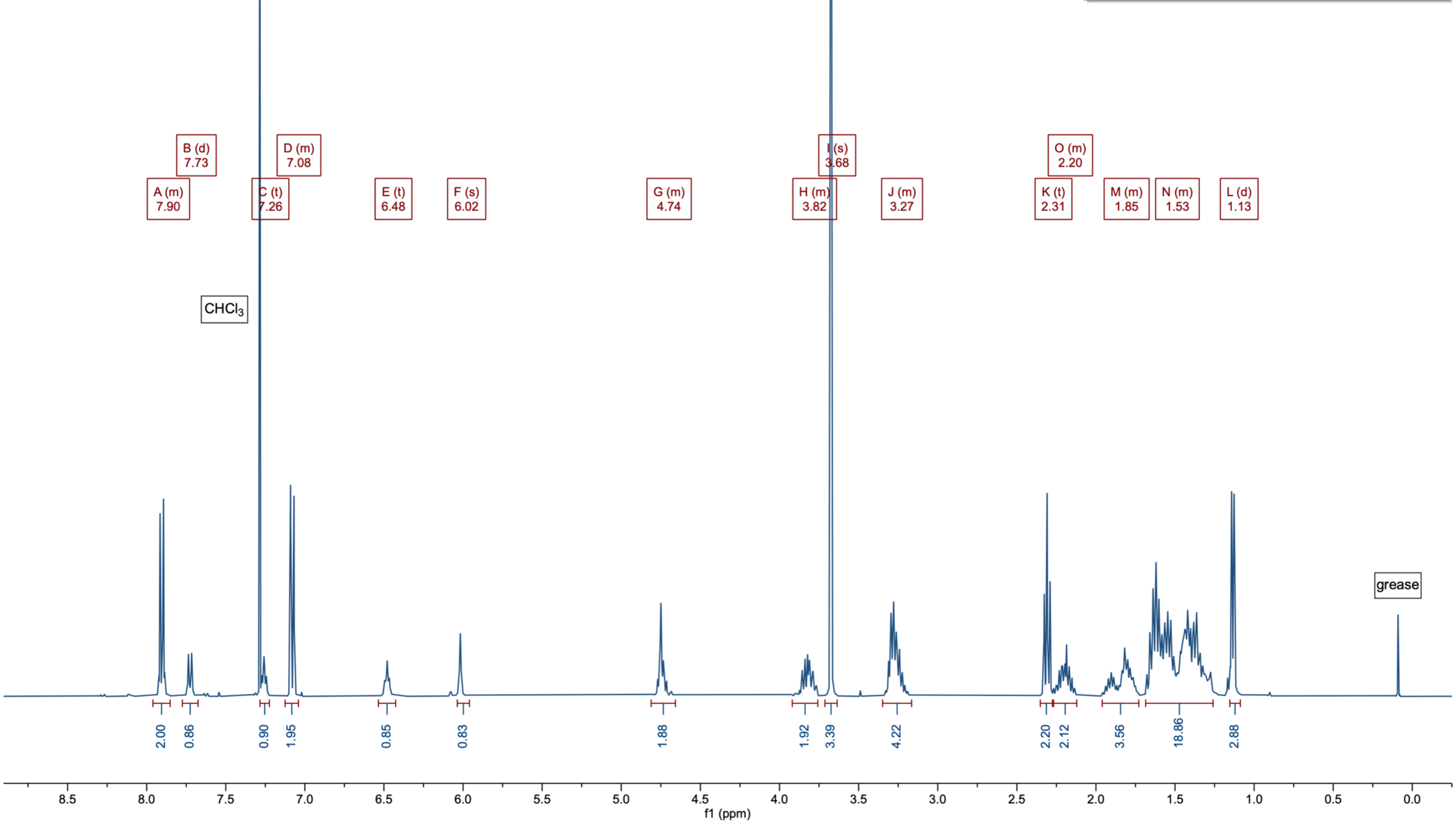

Compound 18: $^{13}C\{^{1}H\}$ – NMR ($CDCl_3$, 101 MHz)

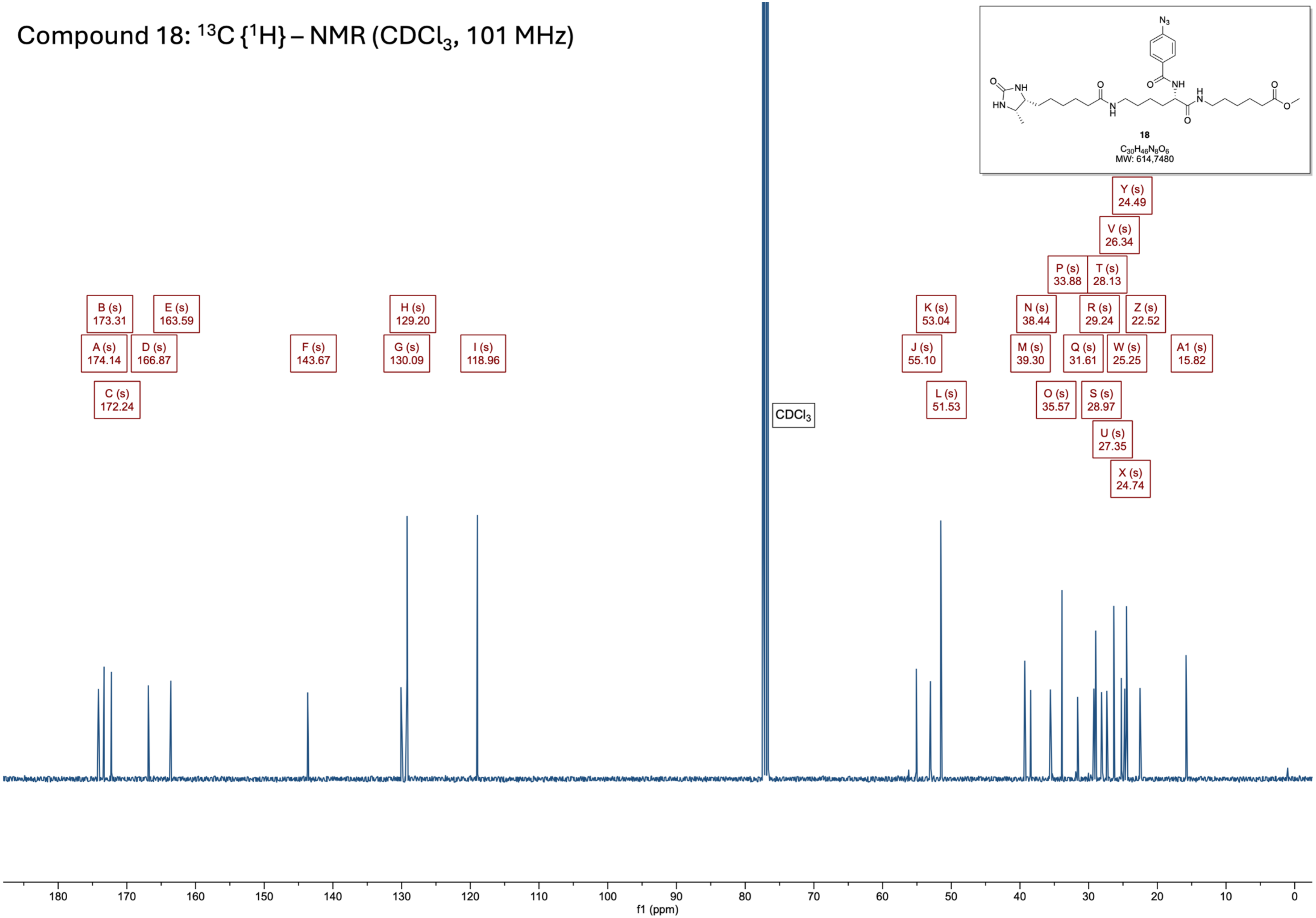

Compound 19: $^{1}H$ – NMR (DMSO-$d_6$, 400 MHz)

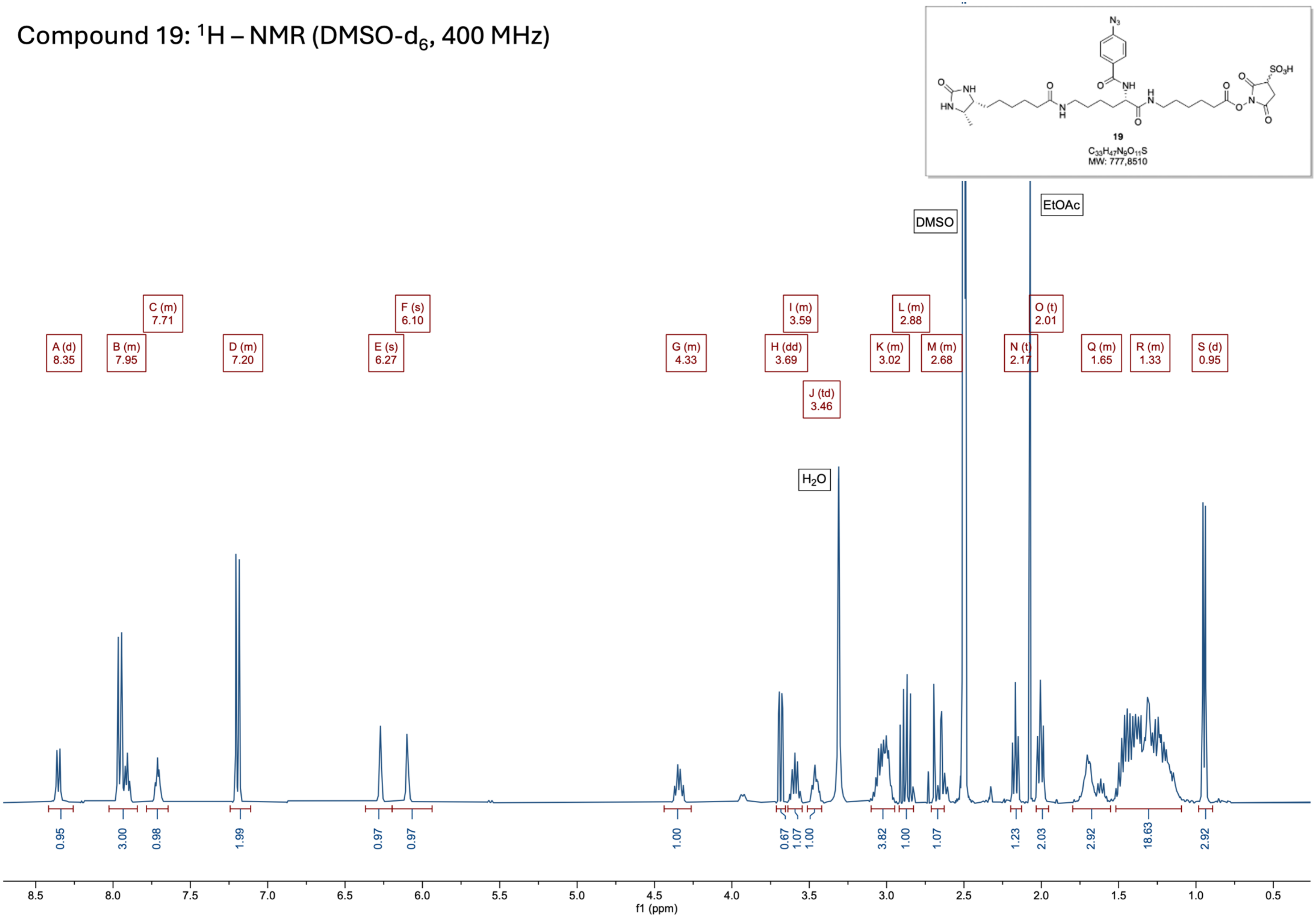

### Compound 19: $^{13}C\{^1H\}$ – NMR (DMSO-$d_6$, 101 MHz)

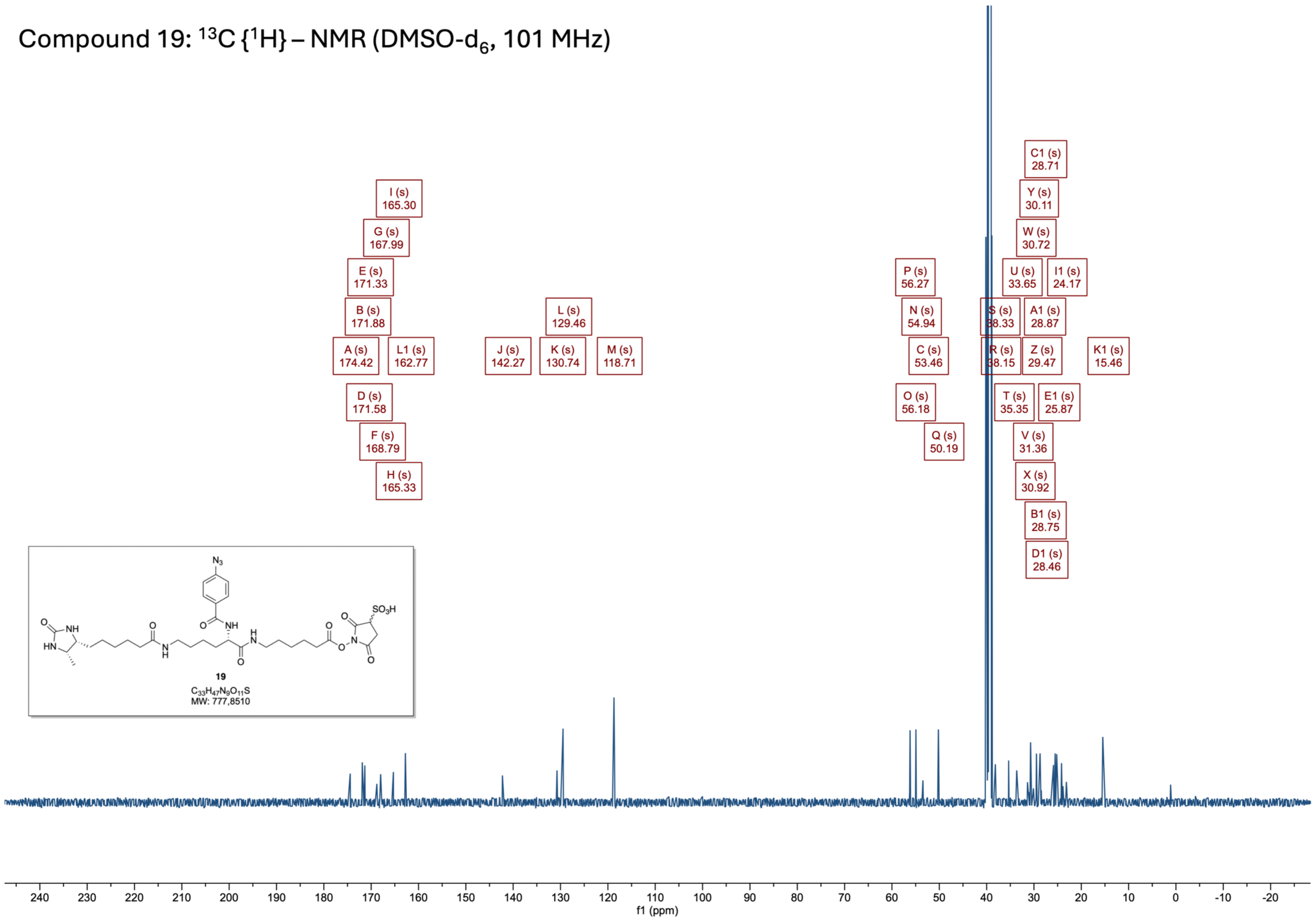

Compound 20: $^{1}H$ – NMR (MeOH-$d_4$, 400 MHz)

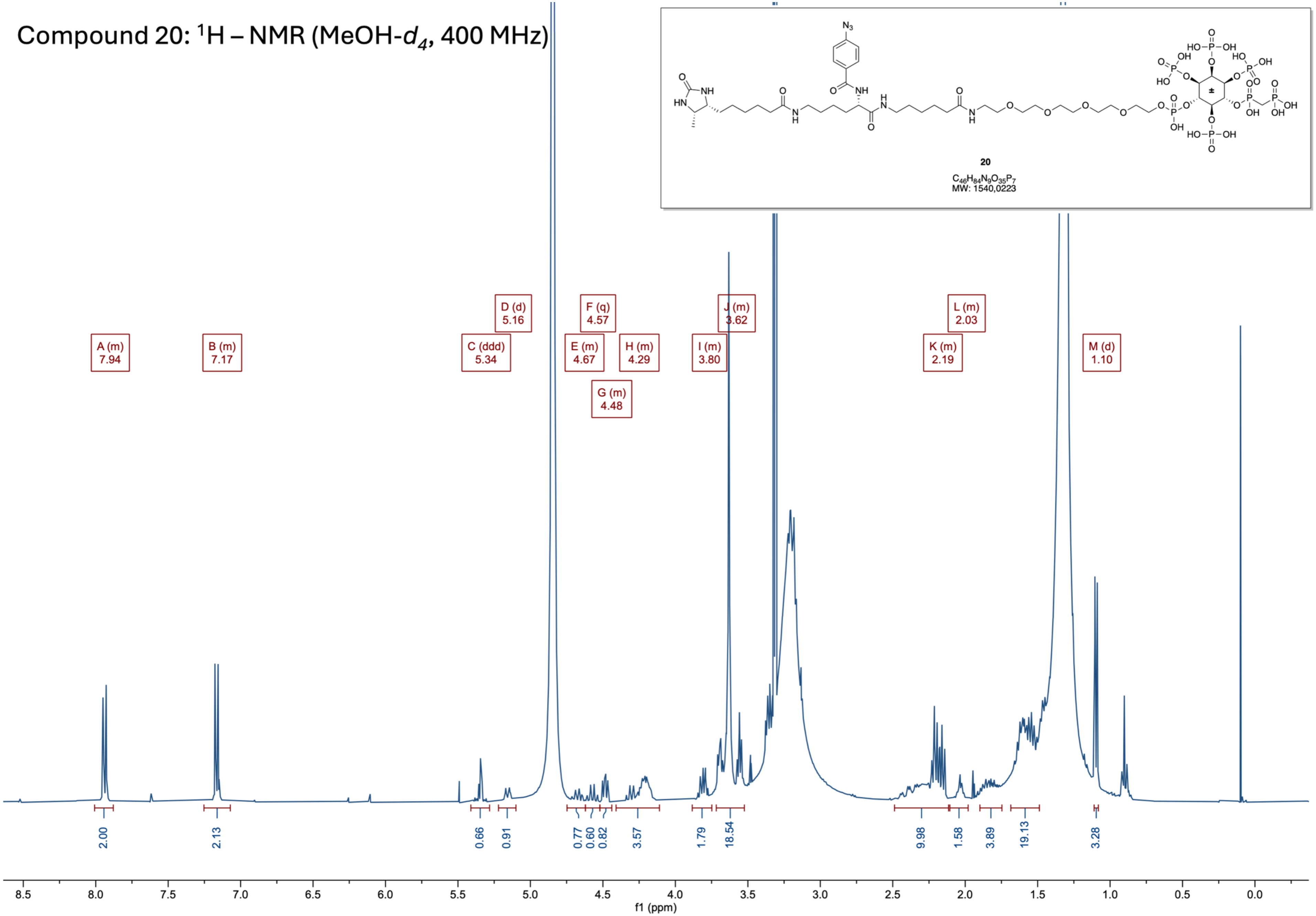

Compound 20: $^{31}P\{^{1}H\}$ – NMR (MeOH-$d_4$, 162 MHz)

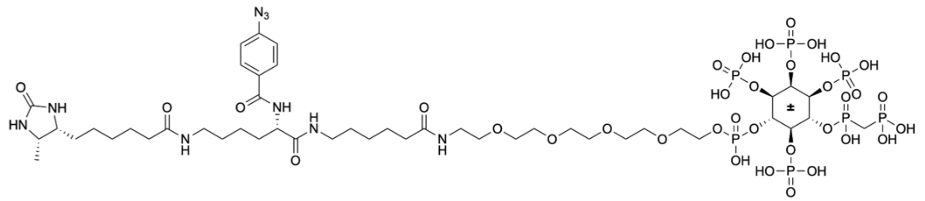

**20**

$C_{46}H_{84}N_9O_{35}P_7$
MW: 1540,0223

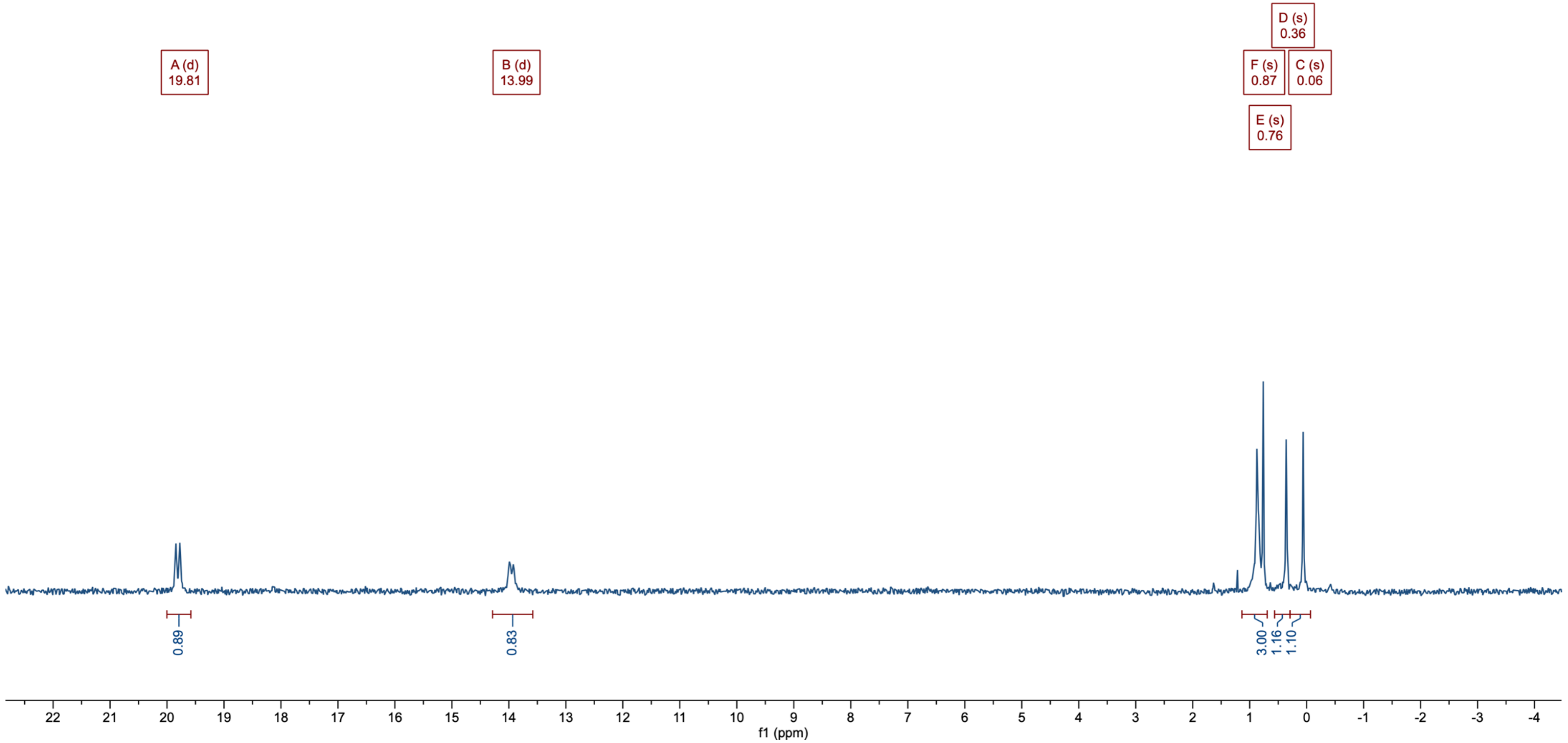

### Compound 20: $^{13}C\{^{1}H\}$ – NMR (MeOH-$d_4$, 101 MHz)

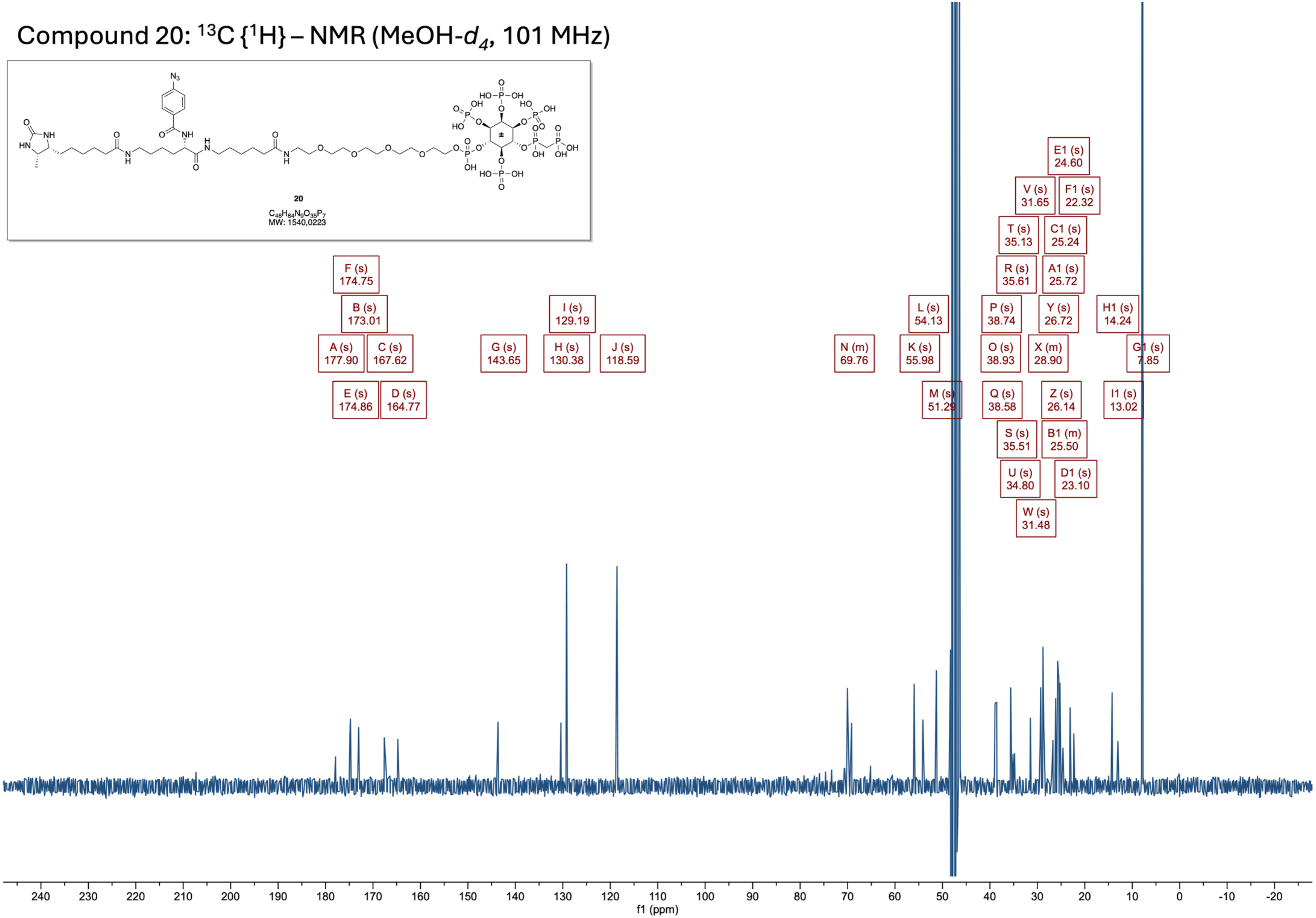

Compound 21: $^1$H – NMR (MeOH-$d_4$, 400 MHz)

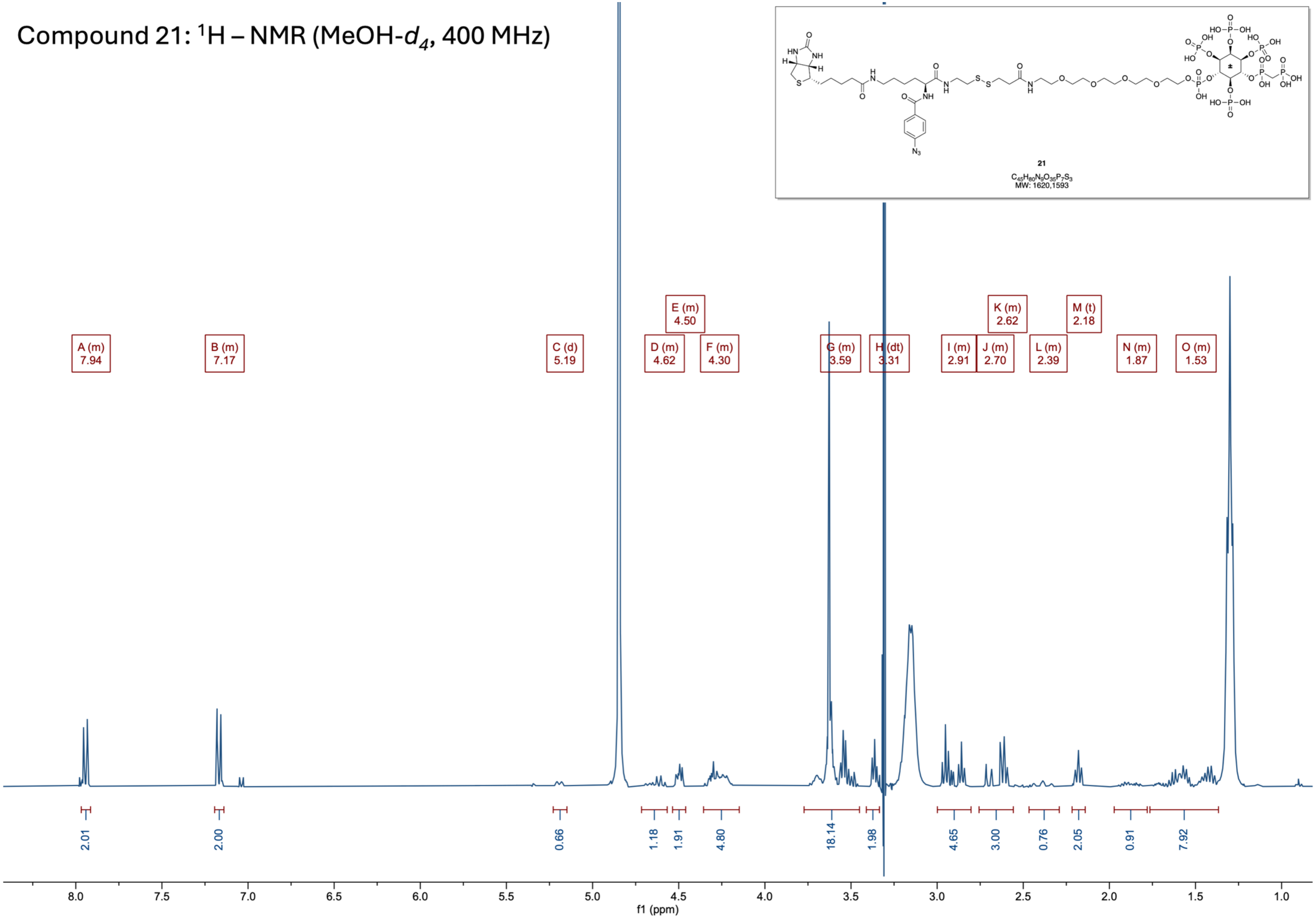

Compound 21: $^{31}P\{^{1}H\}$ – NMR (MeOH-$d_4$, 162 MHz)

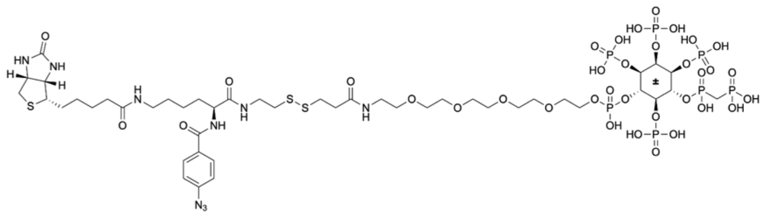

21

$C_{45}H_{80}N_9O_{35}P_7S_3$

MW: 1620,1593

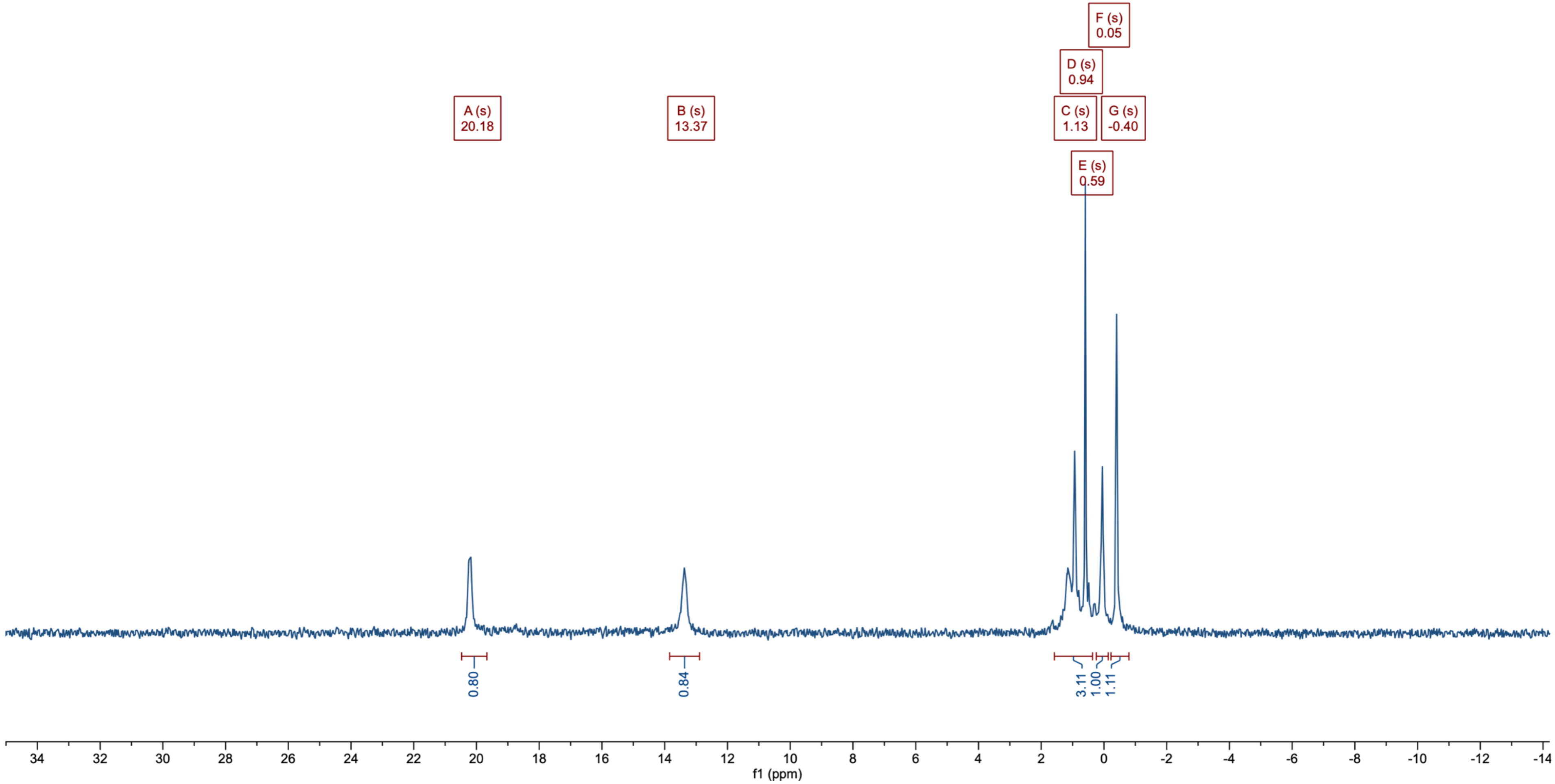

Compound 21: $^{13}C\{^1H\}$ – NMR (MeOH-$d_4$, 101 MHz)

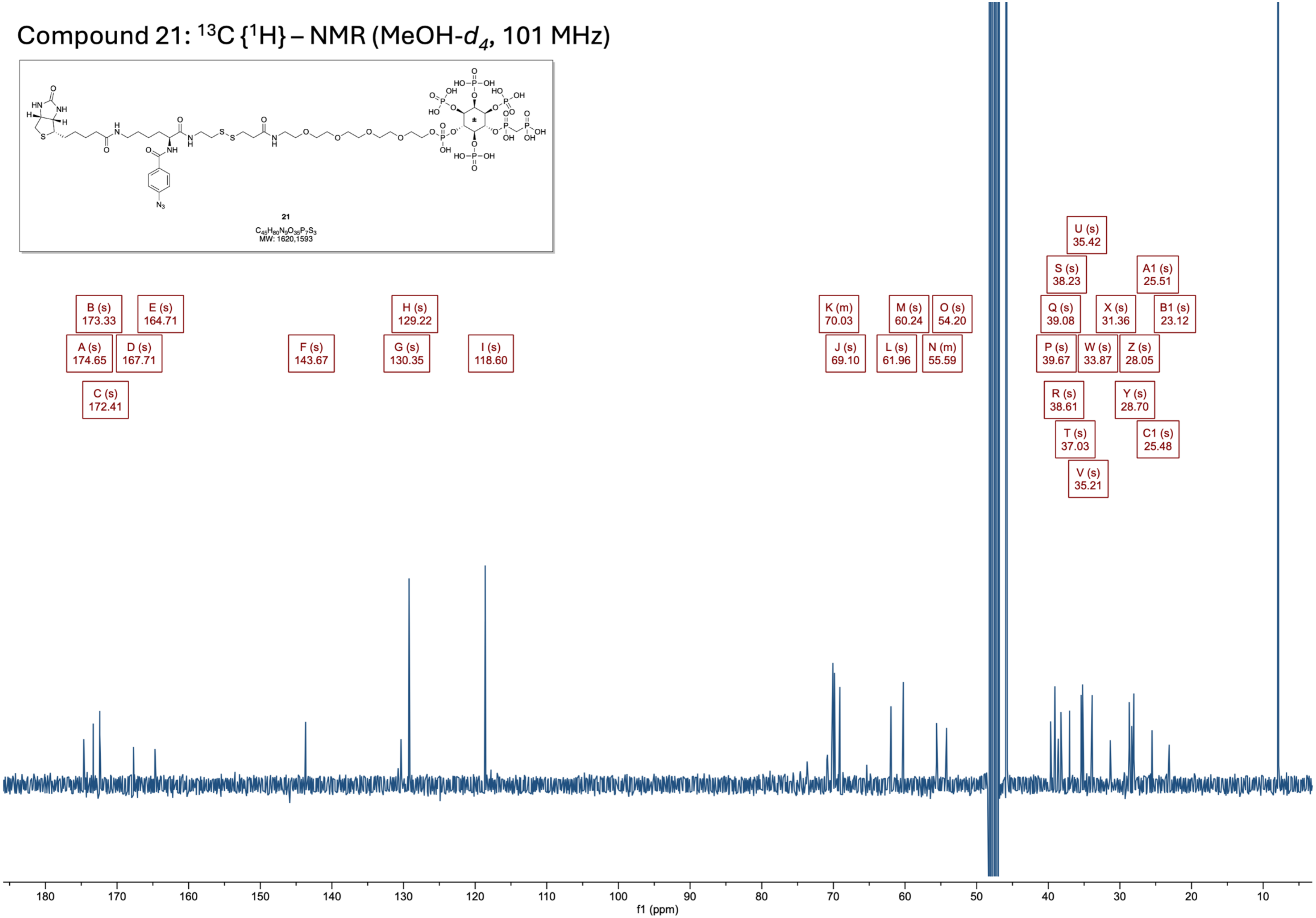

### **9** Mass Spectra, CE Electropherograms & HPLC

(Sorted according to molecule numbering)

### Compound 8: HRMS (APCI) Analysis

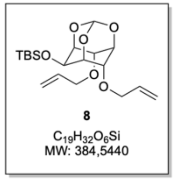


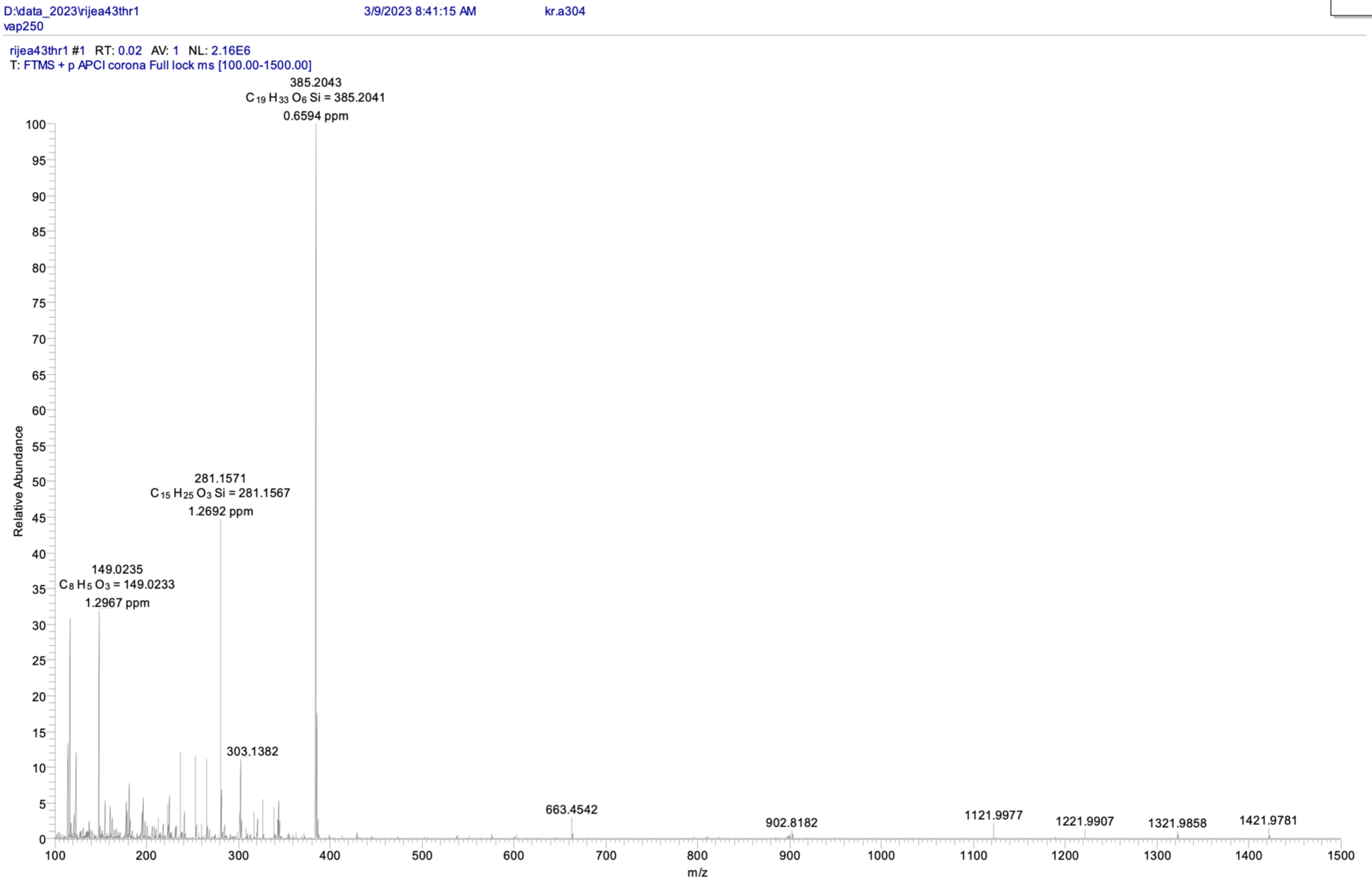

### Compound 9: HRMS (APCI) Analysis

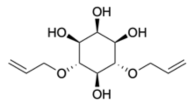

**9**

$C_{12}H_{20}O_6$
MW: 260,2860

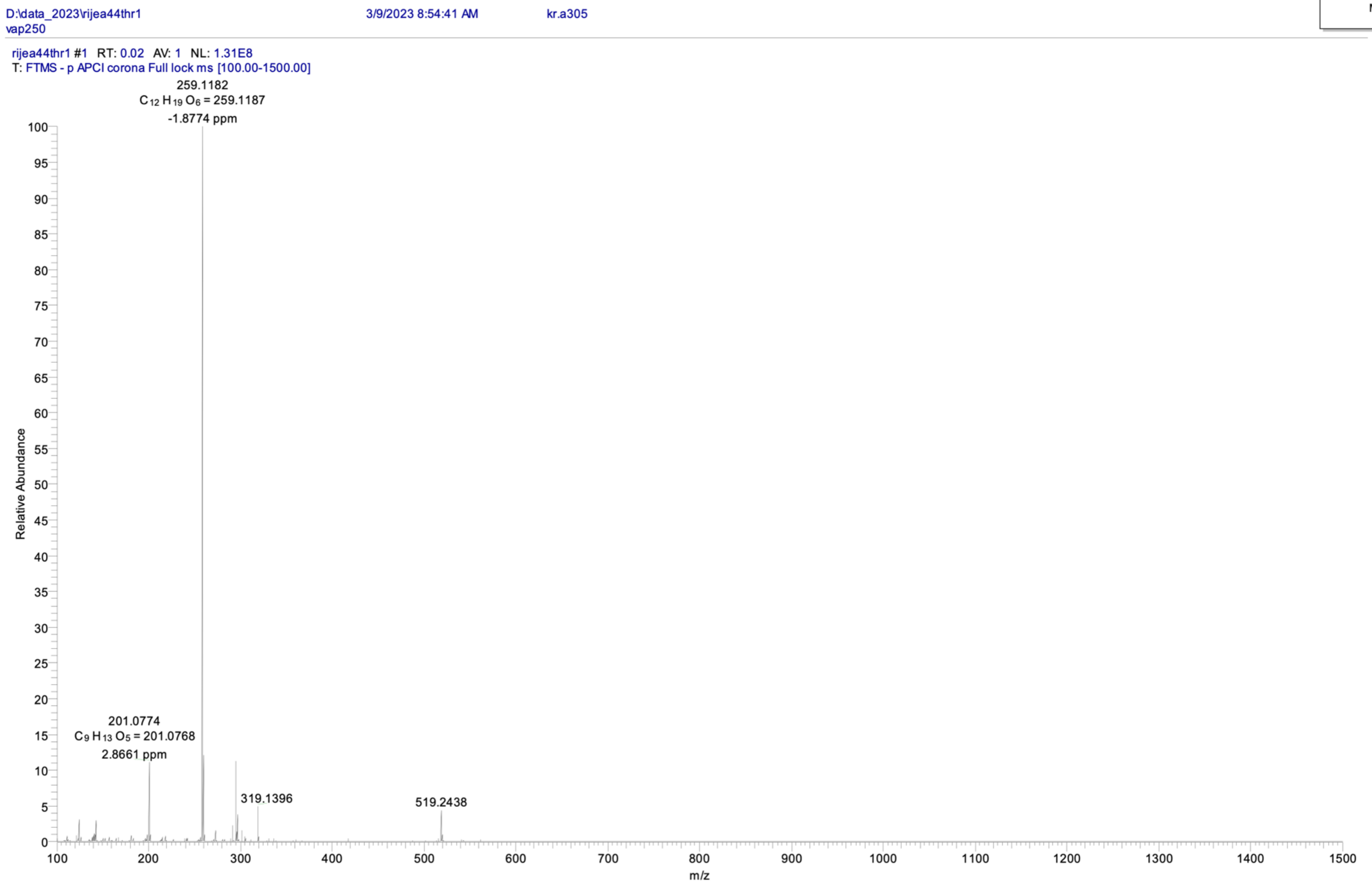

### Compound 10: HRMS (ESI) Analysis

#### Analysis Report

Agilent Trusted Answers

**Sample Information**

| | | | |
|---|---|---|---|
| **Name** | kr.a450 | **Data File Path** | D:\MassHunter\Data\2025\04\rijea75dis01.d |
| **Sample ID** | | **Acq. Time (Local)** | 4/7/2025 1:52:49 PM (UTC+02:00) |
| **Instrument** | QTOF-2 | **Method Path (Acq)** | D:\MassHunter\Methods\Christoph\direkt0,2mlACN.m |
| **MS Type** | QTOF | **Version (Acq SW)** | 6200 series TOF/6500 series Q-TOF 10.1 (48.0) |
| **Inj. Vol. (ul)** | 0 | **IRM Status** | Some ions missed |
| **Position** | | **Method Path (DA)** | |
| **Plate Pos.** | | **Target Source Path** | |
| **Operator** | | **Result Summary** | |

**Sample Chromatograms**

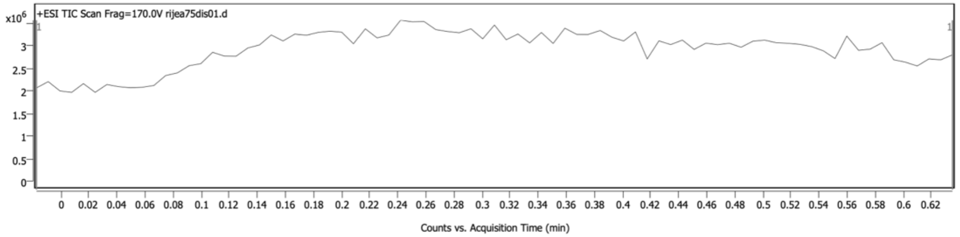


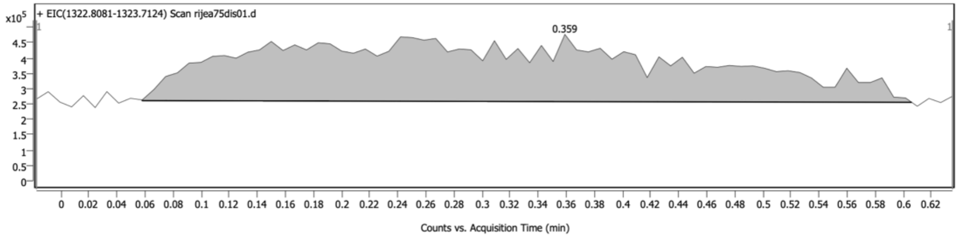


*Chromatogram Peaks*

| Peak | Start | RT | End | Height | Area | Area % | SNR |
|---|---|---|---|---|---|---|---|
| 1 | 0.058 | 0.359 | 0.606 | 218633 | 4384157 | 100.00 | |

**Sample Spectra**

**+ Scan (rt: 0.200-0.250 min) Sub**

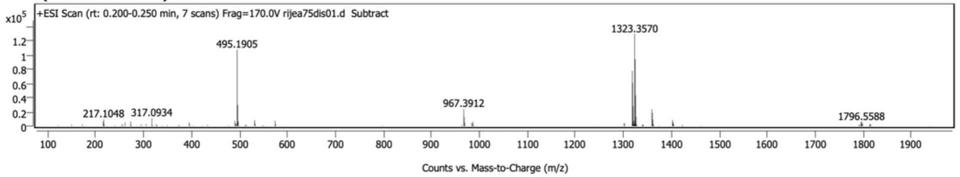


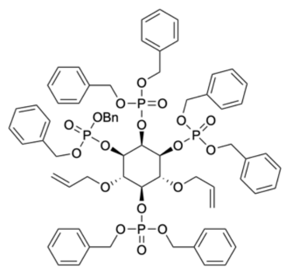

**10**

$C_{68}H_{72}O_{18}P_4$
MW: 1301,2010

### Compound 11: HRMS (ESI) Analysis

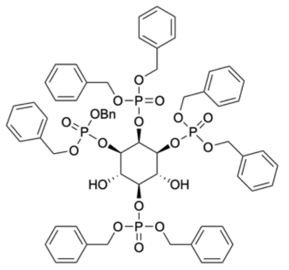

**11**

$C_{62}H_{64}O_{18}P_4$
MW: 1221,0710

**Sample Chromatograms**

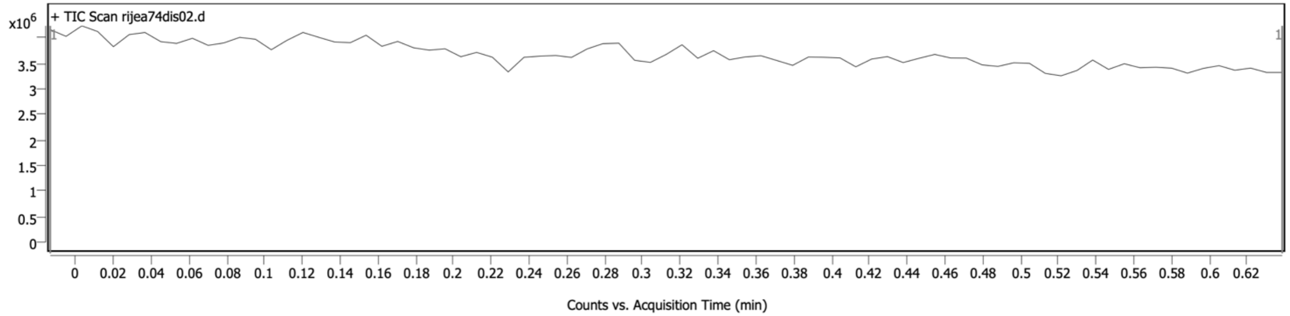


**Sample Spectra**

**+ Scan (rt: -0.005-0.011 min) Sub**

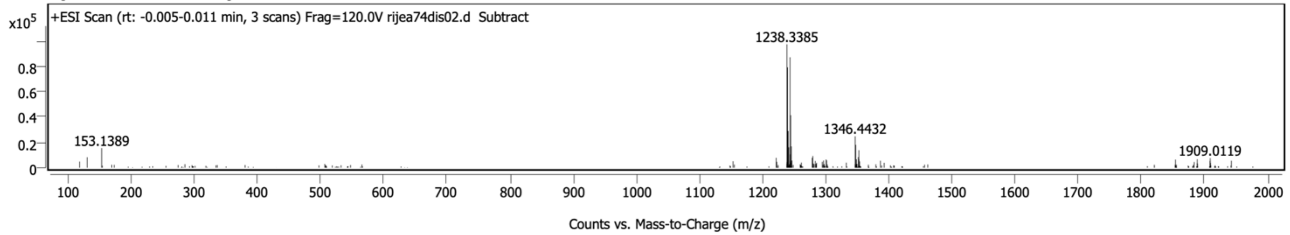

### Compound 12: HRMS (ESI) Analysis

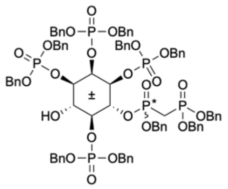


**12**

$C_{84}H_{86}O_{23}P_6$
MW: 1649,4316

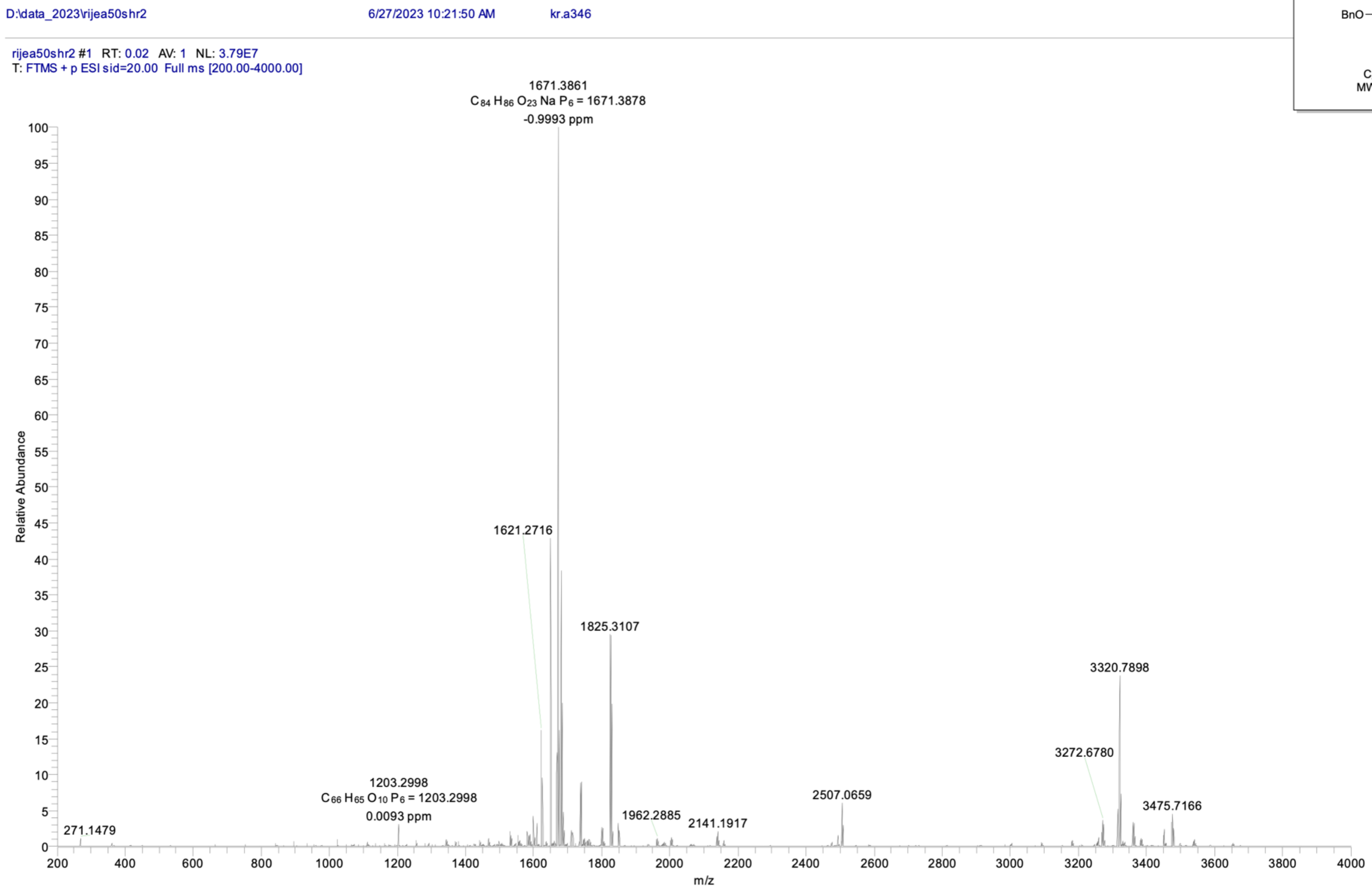

### Compound 13: HRMS (ESI) Analysis

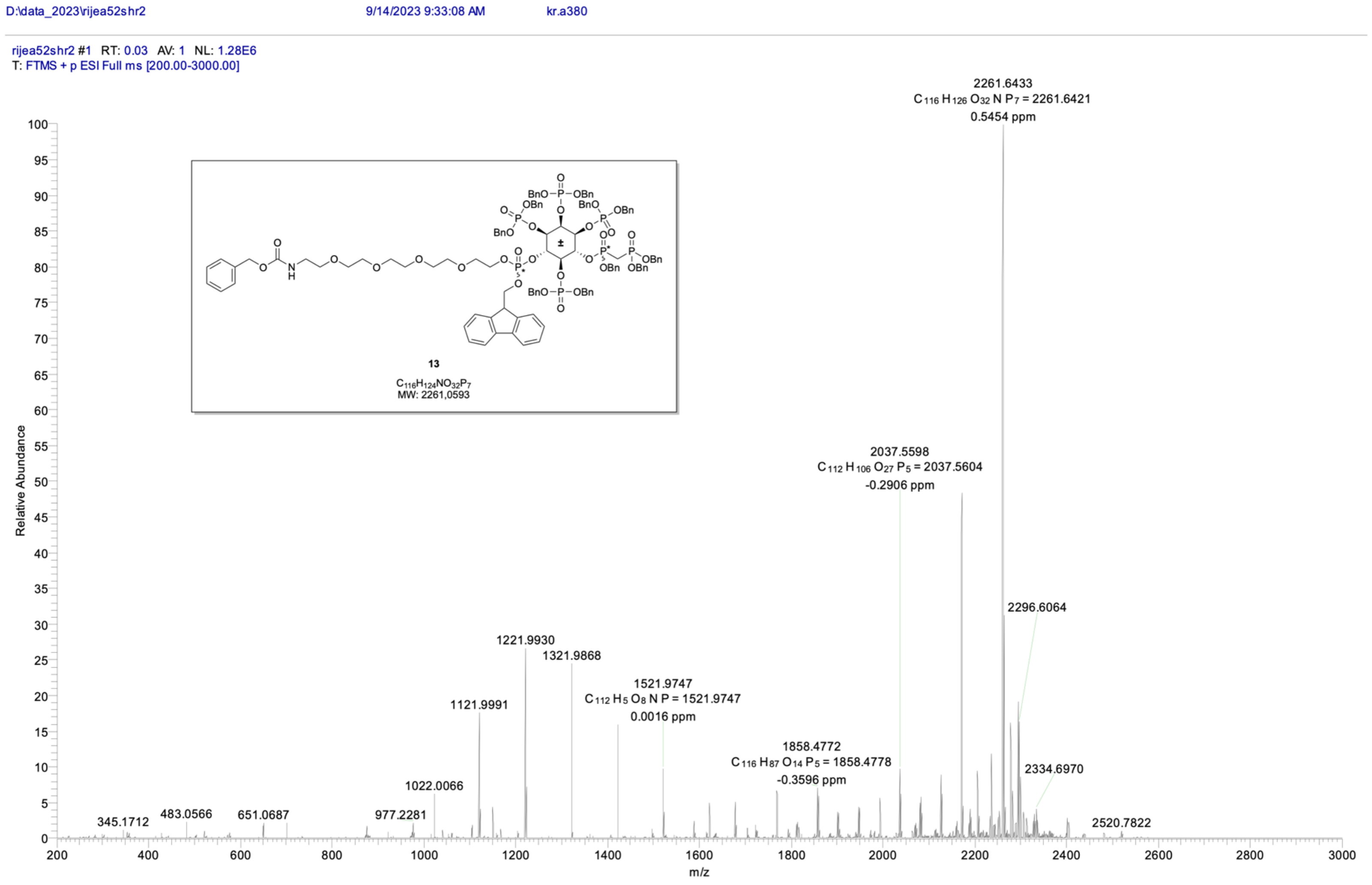

### Compound 14: HRMS (ESI) Analysis

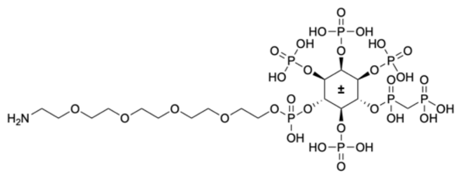

**14**

$C_{17}H_{42}NO_{30}P_7$
MW: 957,3163

| | | | | | |
|---|---|---|---|---|---|
| **Sample Name** | KR-A381 | **Position** | 42 | **Instrument Name** | QTOF-1 |
| **User Name** | | **Inj Vol** | Unknown / Injection Program | **InjPosition** | |
| **Sample Type** | Sample | **IRM Calibration Status** | Success | **Data Filename** | KR-A381.d |
| **ACQ Method** | Standard method.m | **Comment** | | **Acquired Time** | 16.09.2023 19:54:29 (UTC+02:00) |

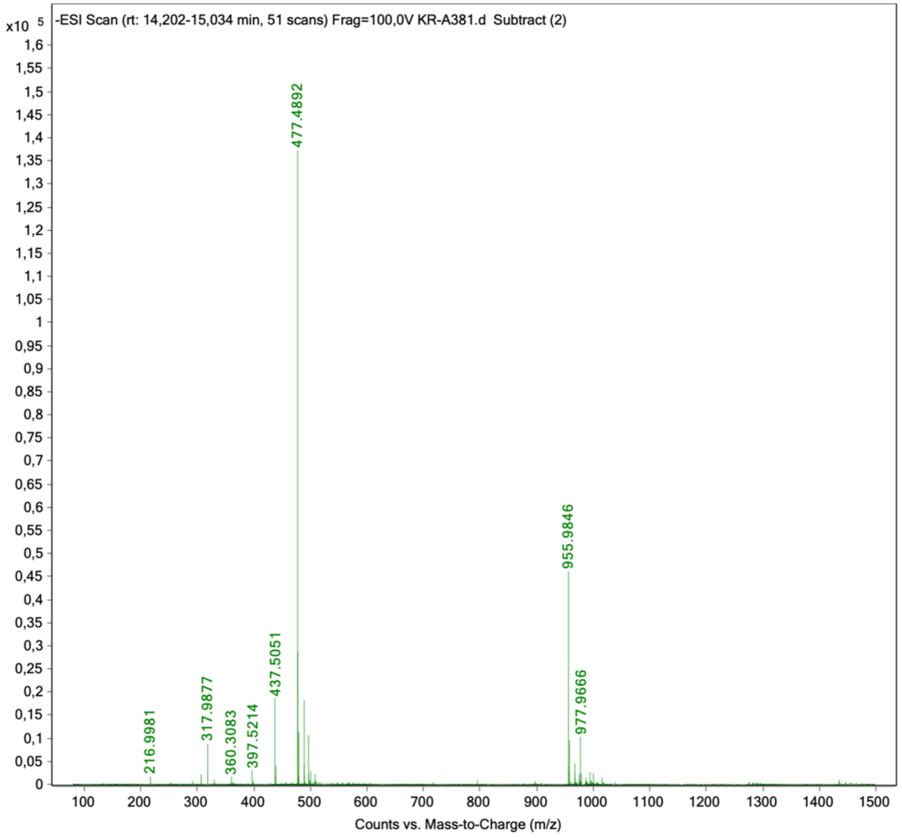

### Compound 16: HRMS (ESI) Analysis

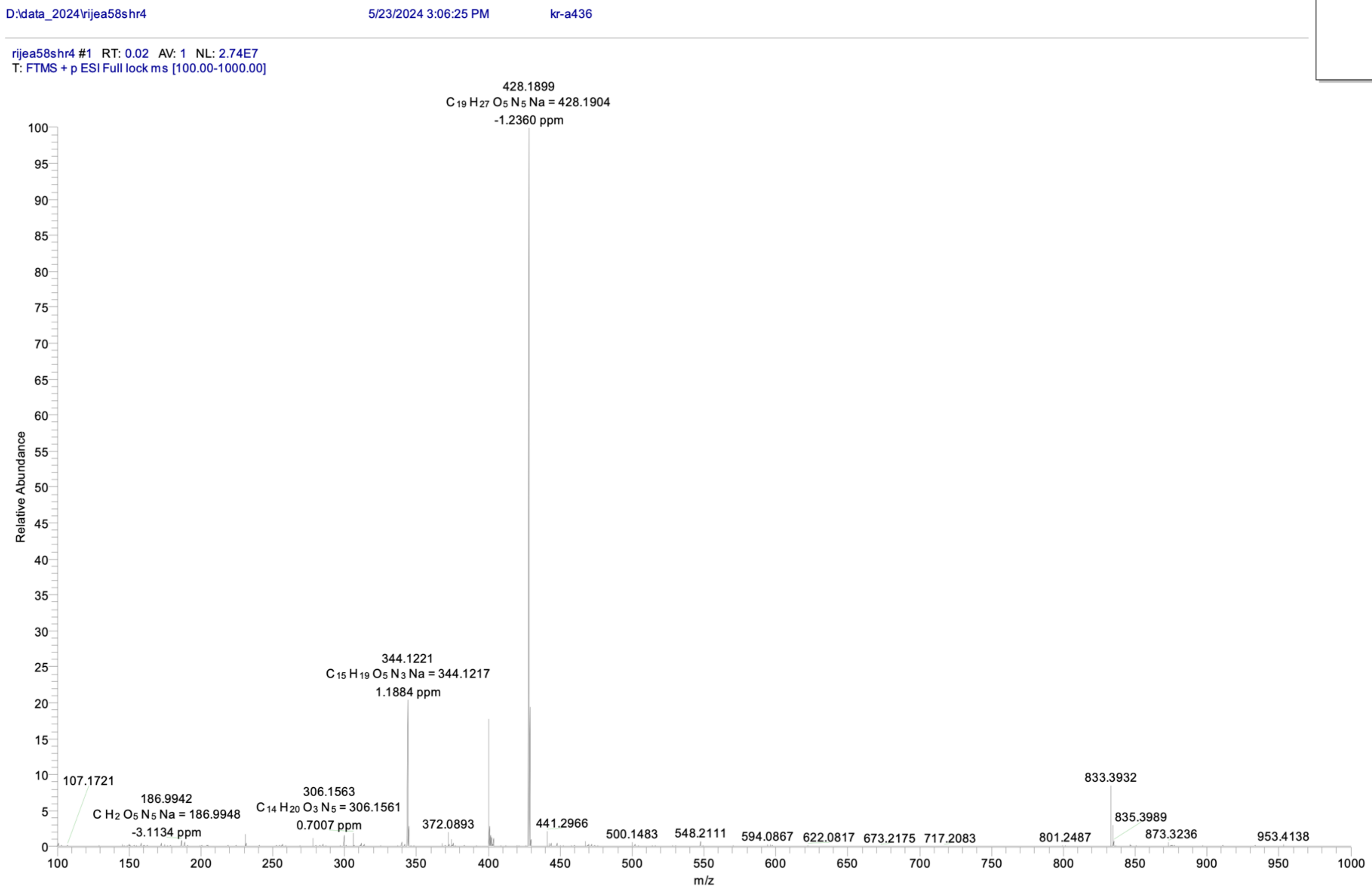


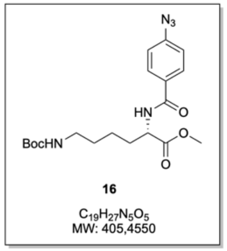

### Compound 17: HRMS (ESI) Analysis

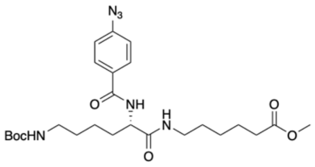


**17**

$C_{25}H_{38}N_6O_6$
MW: 518,6150

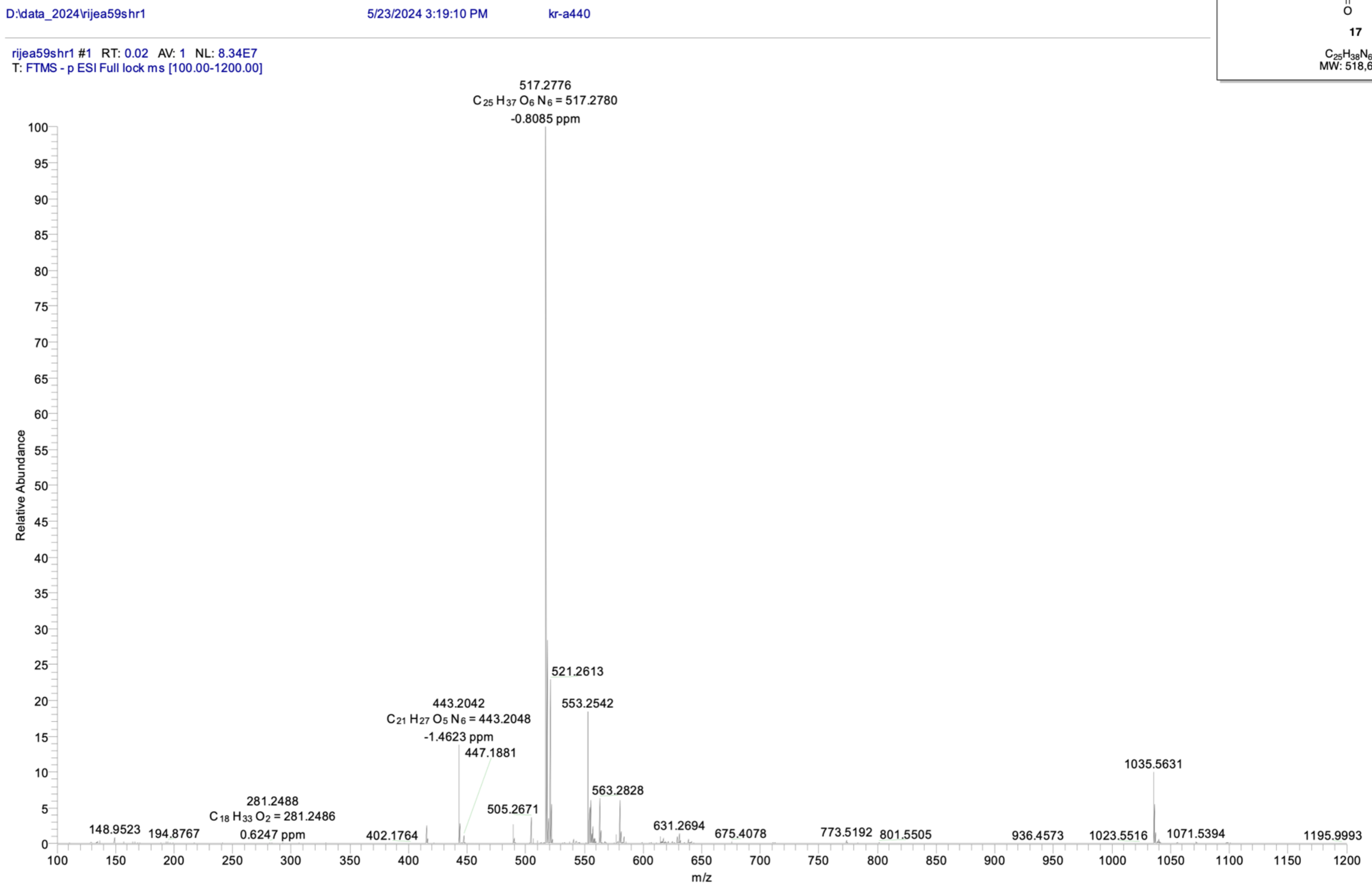

### Compound 18: HRMS (ESI) Analysis

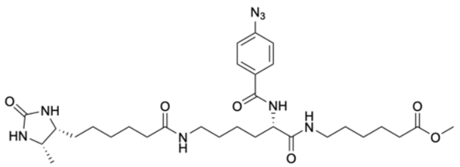


**18**

$C_{30}H_{46}N_8O_6$
MW: 614,7480

#### Analysis Report

Agilent | Trusted Answers

##### Sample Information

| | | | |
|---|---|---|---|
| **Name** | kr-a441 | **Data File Path** | D:\MassHunter\Data\2025\04\rijea73dis01.d |
| **Sample ID** | | **Acq. Time (Local)** | 4/2/2025 3:35:30 PM (UTC+02:00) |
| **Instrument** | QTOF-2 | **Method Path (Acq)** | D:\MassHunter\Methods\Christoph\direkt0,2mlACN.m |
| **MS Type** | QTOF | **Version (Acq SW)** | 6200 series TOF/6500 series Q-TOF 10.1 (48.0) |
| **Inj. Vol. (ul)** | 0 | **IRM Status** | All ions missed |
| **Position** | | **Method Path (DA)** | |
| **Plate Pos.** | | **Target Source Path** | |
| **Operator** | | **Result Summary** | |

##### Sample Chromatograms

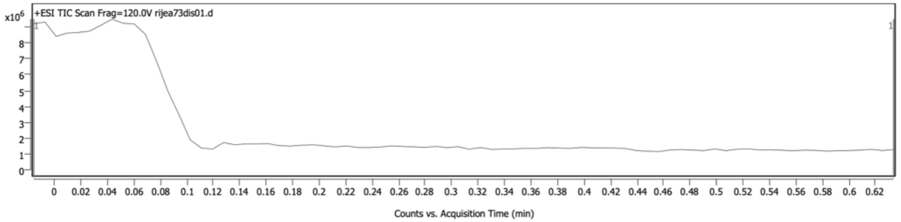


##### Sample Spectra

**+ Scan (rt: 0.027-0.060 min) Sub**

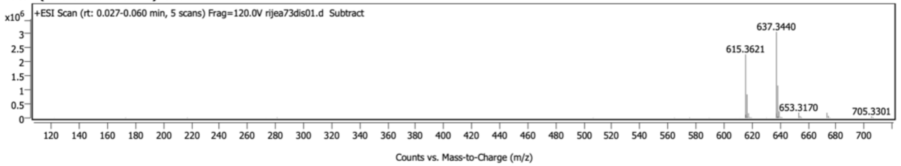


*Spectrum Peaks*

| m/z | Z | Abund | Abund % | m/z (Calc) | Diff (ppm) | Ion Species | Formula | Ion Type |
|---|---|---|---|---|---|---|---|---|
| 615.3621 | 1 | 2271542 | 73.92 | 615.3613 | 1.22 | (M+H)+ | C30 H46 N8 O6 | |
| 616.3648 | 1 | 841146 | 27.37 | 616.3642 | 0.91 | (M+H)+ | C30 H46 N8 O6 | |
| 617.3669 | 1 | 173511 | 5.65 | 617.3668 | 0.10 | (M+H)+ | C30 H46 N8 O6 | |
| 618.3693 | 1 | 31521 | 1.03 | 618.3694 | -0.07 | (M+H)+ | C30 H46 N8 O6 | |
| 637.3440 | 1 | 3072784 | 100.00 | 637.3433 | 1.23 | (M+Na)+ | C30 H46 N8 O6 | |
| 638.3468 | 1 | 1158552 | 37.70 | 638.3461 | 1.08 | (M+Na)+ | C30 H46 N8 O6 | |
| 639.3491 | 1 | 240628 | 7.83 | 639.3488 | 0.51 | (M+Na)+ | C30 H46 N8 O6 | |
| 640.3512 | 1 | 41502 | 1.35 | 640.3513 | -0.23 | (M+Na)+ | C30 H46 N8 O6 | |
| 615.4147 | 1 | 115485 | 3.76 | | | | | |
| 616.4184 | 1 | 42547 | 1.38 | | | | | |
| 637.3968 | 1 | 163628 | 5.33 | | | | | |
| 638.4011 | 1 | 58200 | 1.89 | | | | | |
| 653.3170 | 1 | 195182 | 6.35 | | | | | |
| 654.3200 | 1 | 68973 | 2.24 | | | | | |
| 673.4143 | 1 | 187053 | 6.09 | | | | | |
| 674.4166 | 1 | 72719 | 2.37 | | | | | |
| 705.3301 | | 57257 | 1.86 | | | | | |

*Spectrum Identification Table*

| Best | ID Source | Name | Formula | Species | m/z | Diff (ppm) | CAS | Score | Score (Lib) | Score (DB) | Score (MFG) | Lib/DB |
|---|---|---|---|---|---|---|---|---|---|---|---|---|
| No | MFG | | C30 H46 N8 O6 | (M+H)+ | 615.3621 | 1.07 | | 99.09 | | | 99.09 | |
| No | MFG | | C30 H46 N8 O6 | (M+Na)+ | 637.3440 | 1.18 | | 98.85 | | | 98.85 | |

##### Sample Information

| | | | |
|---|---|---|---|
| **Name** | kr-a441 | **Data File Path** | D:\MassHunter\Data\2025\04\rijea73dis02.d |
| **Sample ID** | | **Acq. Time (Local)** | 4/2/2025 3:36:59 PM (UTC+02:00) |
| **Instrument** | QTOF-2 | **Method Path (Acq)** | D:\MassHunter\Methods\Christoph\direkt0,2mlACN.m |
| **MS Type** | QTOF | **Version (Acq SW)** | 6200 series TOF/6500 series Q-TOF 10.1 (48.0) |
| **Inj. Vol. (ul)** | 0 | **IRM Status** | Some ions missed |
| **Position** | | **Method Path (DA)** | |
| **Plate Pos.** | | **Target Source Path** | |
| **Operator** | | **Result Summary** | |

### Compound 19: HRMS (ESI) Analysis

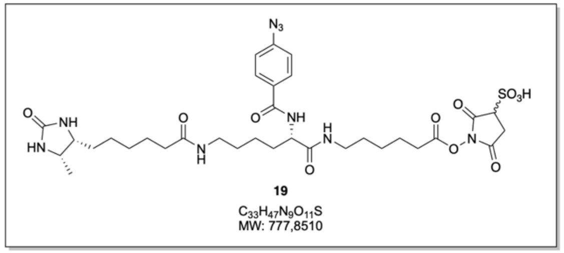


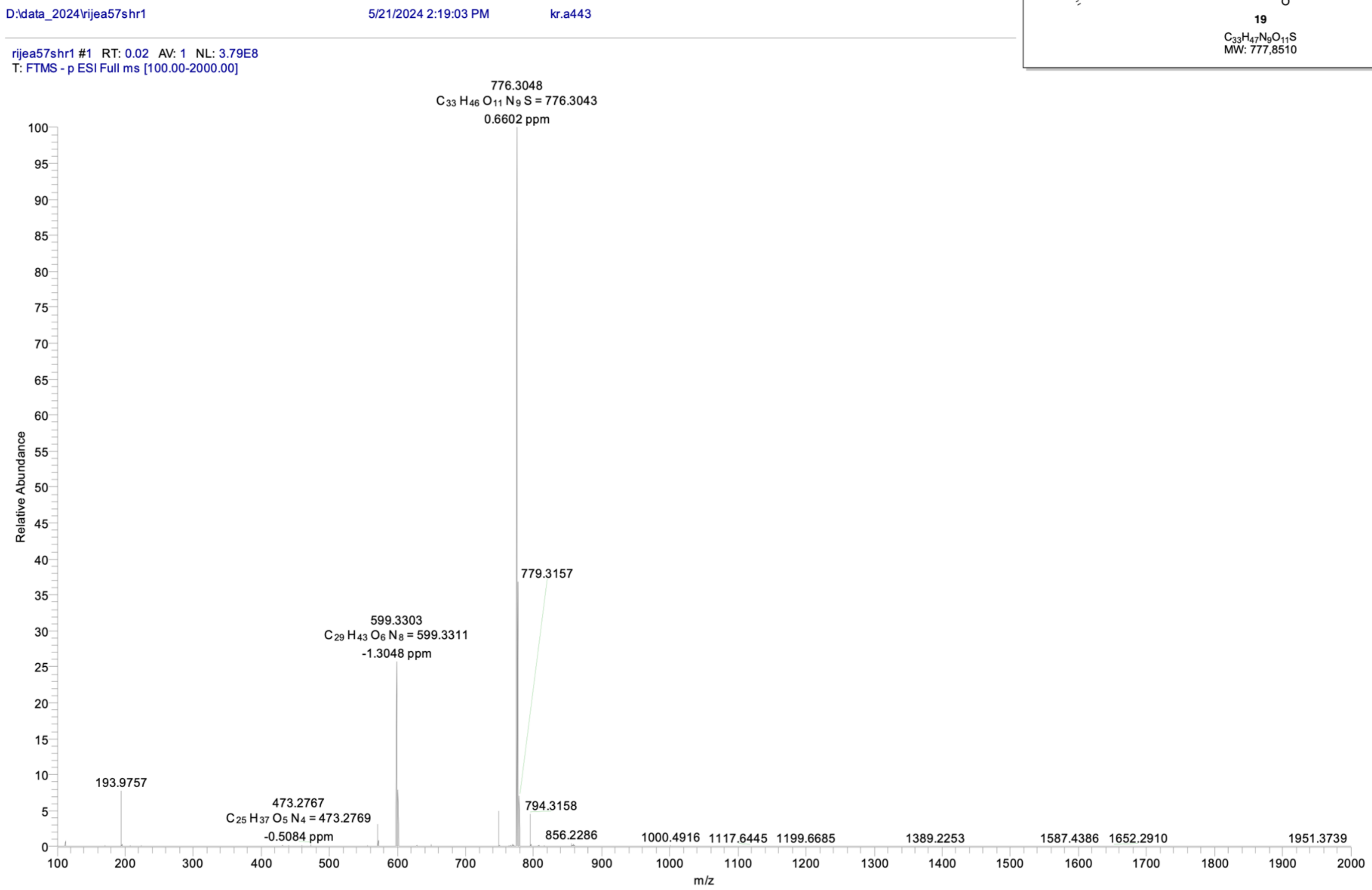

### Compound 20: HRMS (ESI) Analysis

#### Analysis Report

Agilent | Trusted Answers

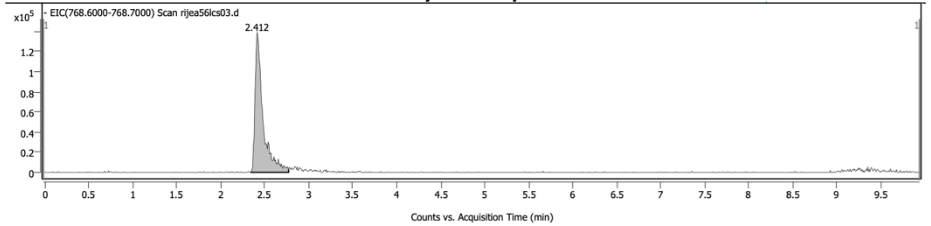


*Chromatogram Peaks*

| Peak | Start | RT | End | Height | Area | Area % | SNR |
|---|---|---|---|---|---|---|---|
| 1 | 2.338 | 2.412 | 2.772 | 138777 | 878178 | 100.00 | |

**Sample Spectra**

**- Scan (rt: 2.379-2.546 min) Sub** **Peak 1 from - EIC(768.6000-768.7000) Scan**

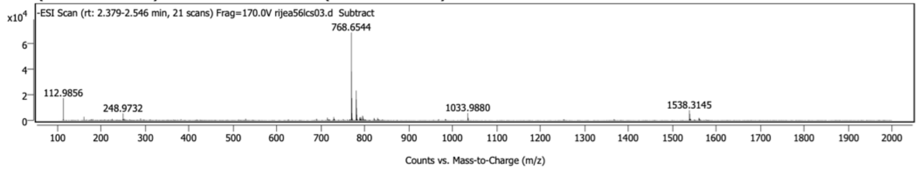


*Spectrum Peaks*

| m/z | Z | Abund | Abund % | m/z (Calc) | Diff (ppm) | Ion Species | Formula | Ion Type |
|---|---|---|---|---|---|---|---|---|
| 768.6544 | 2 | 69065 | 100.00 | 768.6544 | 0.02 | (M-2H)-2 | C46 H84 N9 O35 P7 | |
| 769.1559 | 2 | 38364 | 55.55 | 769.1559 | 0.02 | (M-2H)-2 | C46 H84 N9 O35 P7 | |
| 769.6570 | 2 | 17442 | 25.25 | 769.6571 | -0.14 | (M-2H)-2 | C46 H84 N9 O35 P7 | |
| 770.1586 | 2 | 5688 | 8.24 | 770.1584 | 0.30 | (M-2H)-2 | C46 H84 N9 O35 P7 | |
| 770.6581 | 2 | 1883 | 2.73 | 770.6596 | -1.91 | (M-2H)-2 | C46 H84 N9 O35 P7 | |
| 1033.9880 | 1 | 5745 | 8.32 | 1033.9870 | 0.97 | (M-H)- | C35 H30 N8 O16 P7 | |
| 1034.9911 | 1 | 1939 | 2.81 | 1034.9900 | 1.04 | (M-H)- | C35 H30 N8 O16 P7 | |
| 1538.3145 | 1 | 8035 | 11.63 | 1538.3160 | -0.98 | (M-H)- | C46 H84 N9 O35 P7 | |
| 1539.3178 | 1 | 5105 | 7.39 | 1539.3191 | -0.80 | (M-H)- | C46 H84 N9 O35 P7 | |
| 112.9856 | | 17479 | 25.31 | | | | | |
| 160.0153 | | 2846 | 4.12 | | | | | |
| 248.9732 | | 5479 | 7.93 | | | | | |
| 288.9497 | | 1544 | 2.24 | | | | | |
| 714.3572 | | 2133 | 3.09 | | | | | |
| 728.6712 | | 2671 | 3.87 | | | | | |
| 768.7146 | | 1576 | 2.28 | | | | | |
| 779.6401 | 2 | 23427 | 33.92 | | | | | |
| 780.1414 | 2 | 16464 | 23.84 | | | | | |
| 780.6406 | 2 | 9638 | 13.95 | | | | | |
| 781.1410 | 2 | 4459 | 6.46 | | | | | |
| 787.6287 | | 2294 | 3.32 | | | | | |
| 789.6770 | | 2632 | 3.81 | | | | | |
| 790.6319 | | 2120 | 3.07 | | | | | |
| 794.3145 | 1 | 3701 | 5.36 | | | | | |
| 795.3193 | 1 | 2920 | 4.23 | | | | | |
| 796.3203 | 1 | 1420 | 2.06 | | | | | |
| 820.6401 | | 1796 | 2.60 | | | | | |
| 828.6280 | | 1896 | 2.74 | | | | | |
| 1560.2886 | 1 | 1879 | 2.72 | | | | | |
| 1561.2918 | 1 | 1693 | 2.45 | | | | | |

*Spectrum Identification Table*

| Best | ID Source | Name | Formula | Species | m/z | Diff (ppm) | CAS | Score | Score (Lib) | Score (DB) | Score (MFG) | Lib/DB |
|---|---|---|---|---|---|---|---|---|---|---|---|---|
| No | MFG | | C46 H84 N9 O35 P7 | (M-2H)-2 | 768.6544 | -0.02 | | 98.55 | | | 98.55 | |
| No | MFG | | C35 H30 N8 O16 P7 | (M-H)- | 1033.9880 | 1.00 | | 83.69 | | | 83.69 | |
| No | MFG | | C36 H36 N O21 P7 | (M-H)- | 1033.9880 | 0.89 | | 83.15 | | | 83.15 | |
| No | MFG | | C46 H84 N9 O35 P7 | (M-H)- | 1538.3145 | -0.79 | | 82.83 | | | 82.83 | |

**MassHunter Qual 10.0**
**(End of Report)**



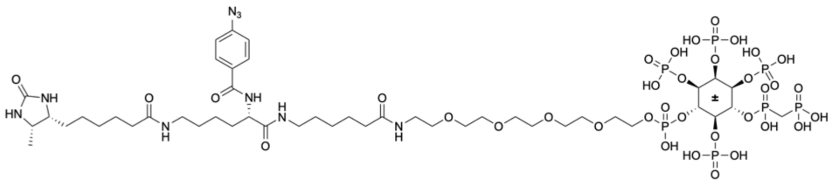

**20**

$C_{46}H_{84}N_9O_{35}P_7$
MW: 1540,0223

### Compound 20: HPLC Analysis

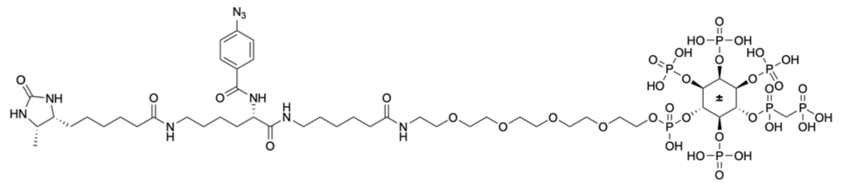

20

$C_{46}H_{84}N_9O_{35}P_7$
MW: 1540,0223

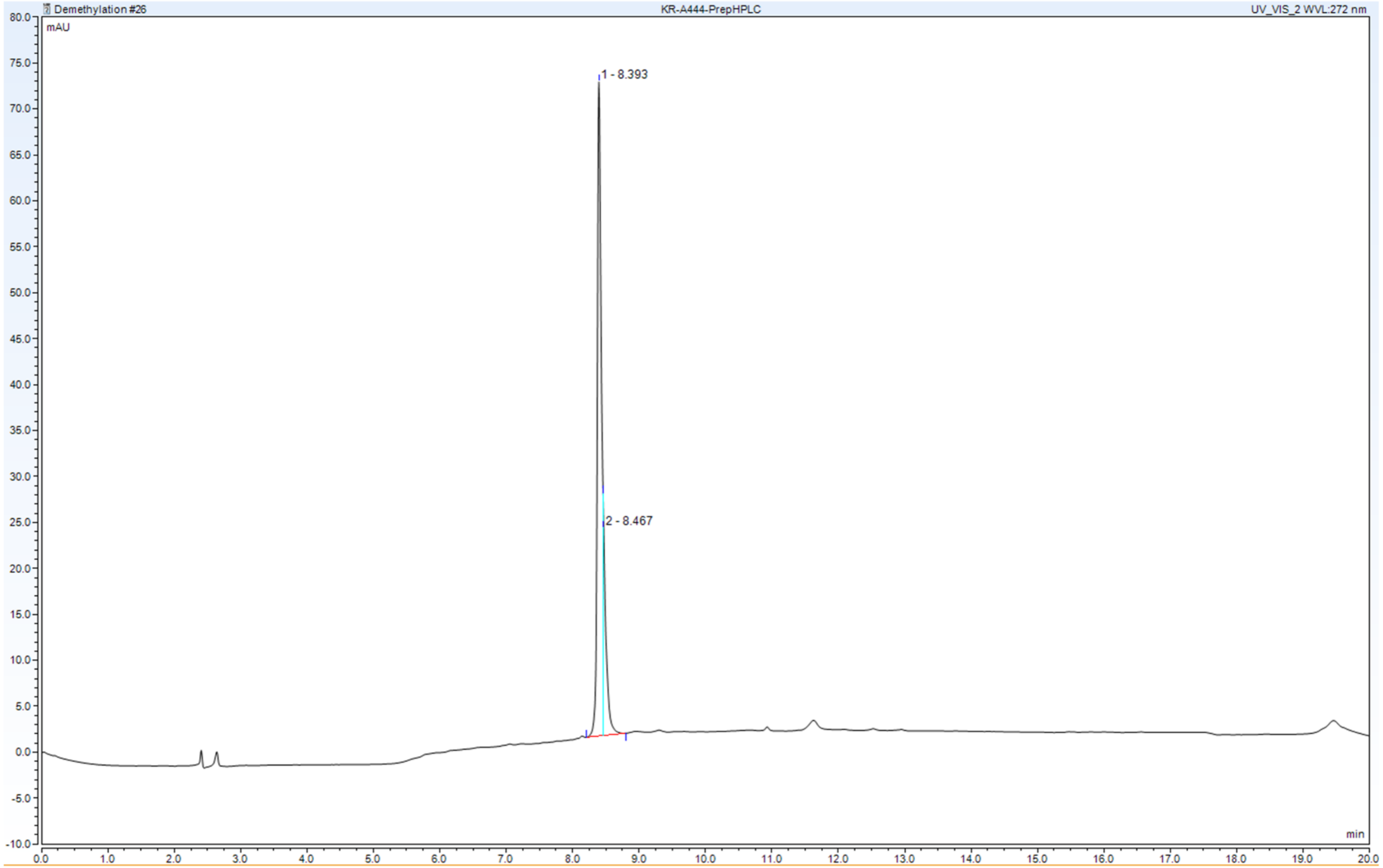

### Compound 21: HRMS (ESI) Analysis

#### Analysis Report

Agilent | Trusted Answers

##### Sample Information

| | | | |
|---|---|---|---|
| **Name** | kr.a383 | **Data File Path** | D:\MassHunter\Data\2023\09\rijea53dis01.d |
| **Sample ID** | lc neg.esi | **Acq. Time (Local)** | 9/20/2023 2:52:40 PM (UTC+02:00) |
| **Instrument** | QTOF-2 | **Method Path (Acq)** | D:\MassHunter\Methods\Christoph\esi-neg90a-100D(8-10)700c20.m |
| **MS Type** | QTOF | **Version (Acq SW)** | 6200 series TOF/6500 series Q-TOF 10.1 (48.0) |
| **Inj. Vol. (ul)** | 0.2 | **IRM Status** | Success |
| **Position** | | **Method Path (DA)** | |
| **Plate Pos.** | | **Target Source Path** | |
| **Operator** | | **Result Summary** | |

##### Sample Chromatograms

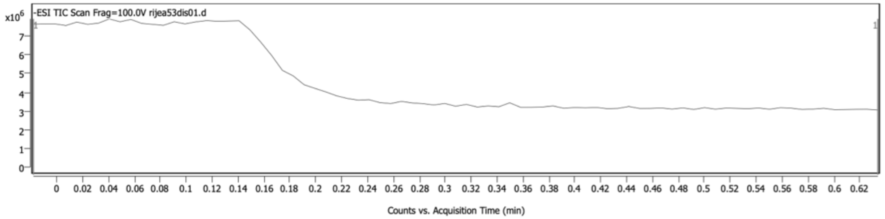


##### Sample Spectra

**- Scan (rt: 0.074-0.116 min) Sub (2)**

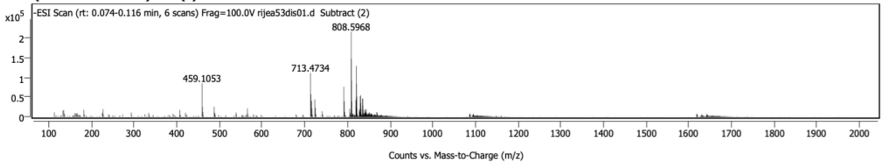


*Spectrum Peaks*

| m/z | Z | Abund | Abund % | m/z (Calc) | Diff (ppm) | Ion Species | Formula | Ion Type |
|---|---|---|---|---|---|---|---|---|
| 808.5968 | 2 | 215185 | 100.00 | 808.5968 | -0.02 | (M-2H)-2 | C45 H80 N9 O35 P7 S3 | |
| 809.0984 | 2 | 146723 | 68.18 | 809.0983 | 0.17 | (M-2H)-2 | C45 H80 N9 O35 P7 S3 | |
| 809.5978 | 2 | 90836 | 42.21 | 809.5977 | 0.04 | (M-2H)-2 | C45 H80 N9 O35 P7 S3 | |
| 810.0980 | 2 | 38882 | 18.07 | 810.0984 | -0.40 | (M-2H)-2 | C45 H80 N9 O35 P7 S3 | |
| 459.1053 | | 84204 | 39.13 | | | | | |
| 713.4734 | 1 | 110827 | 51.50 | | | | | |
| 714.4770 | 1 | 56658 | 26.33 | | | | | |
| 715.4723 | 1 | 39892 | 18.54 | | | | | |
| 723.5018 | | 43490 | 20.21 | | | | | |
| 791.4893 | 1 | 75951 | 35.30 | | | | | |
| 792.4934 | 1 | 39380 | 18.30 | | | | | |
| 819.5858 | | 42182 | 19.60 | | | | | |
| 820.5761 | 2 | 128810 | 59.86 | | | | | |
| 821.0775 | 2 | 83185 | 38.66 | | | | | |
| 821.5769 | 2 | 50201 | 23.33 | | | | | |
| 829.5806 | | 51131 | 23.76 | | | | | |
| 830.5796 | 2 | 54574 | 25.36 | | | | | |
| 831.0805 | 2 | 34453 | 16.01 | | | | | |
| 835.0527 | 2 | 44708 | 20.78 | | | | | |
| 835.5550 | 2 | 46508 | 21.61 | | | | | |

*Spectrum Identification Table*

| Best | ID Source | Name | Formula | Species | m/z | Diff (ppm) | CAS | Score | Score (Lib) | Score (DB) | Score (MFG) | Lib/DB |
|---|---|---|---|---|---|---|---|---|---|---|---|---|
| No | MFG | | C45 H80 N9 O35 P7 S3 | (M-2H)-2 | 808.5968 | 0.03 | | 96.63 | | | 96.63 | |

##### Sample Information



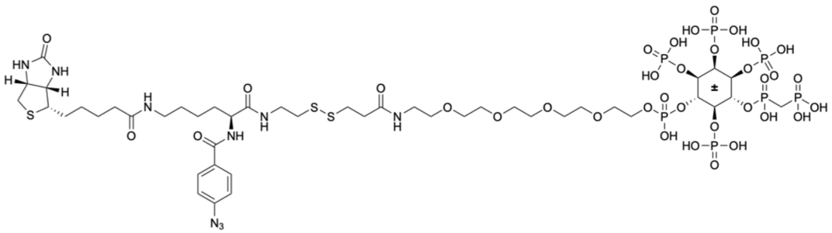

**21**

$C_{45}H_{80}N_9O_{35}P_7S_3$
MW: 1620,1593

### Compound 21: HPLC Analysis

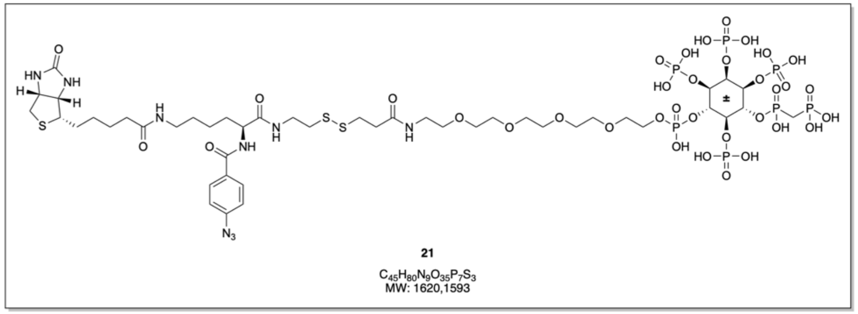

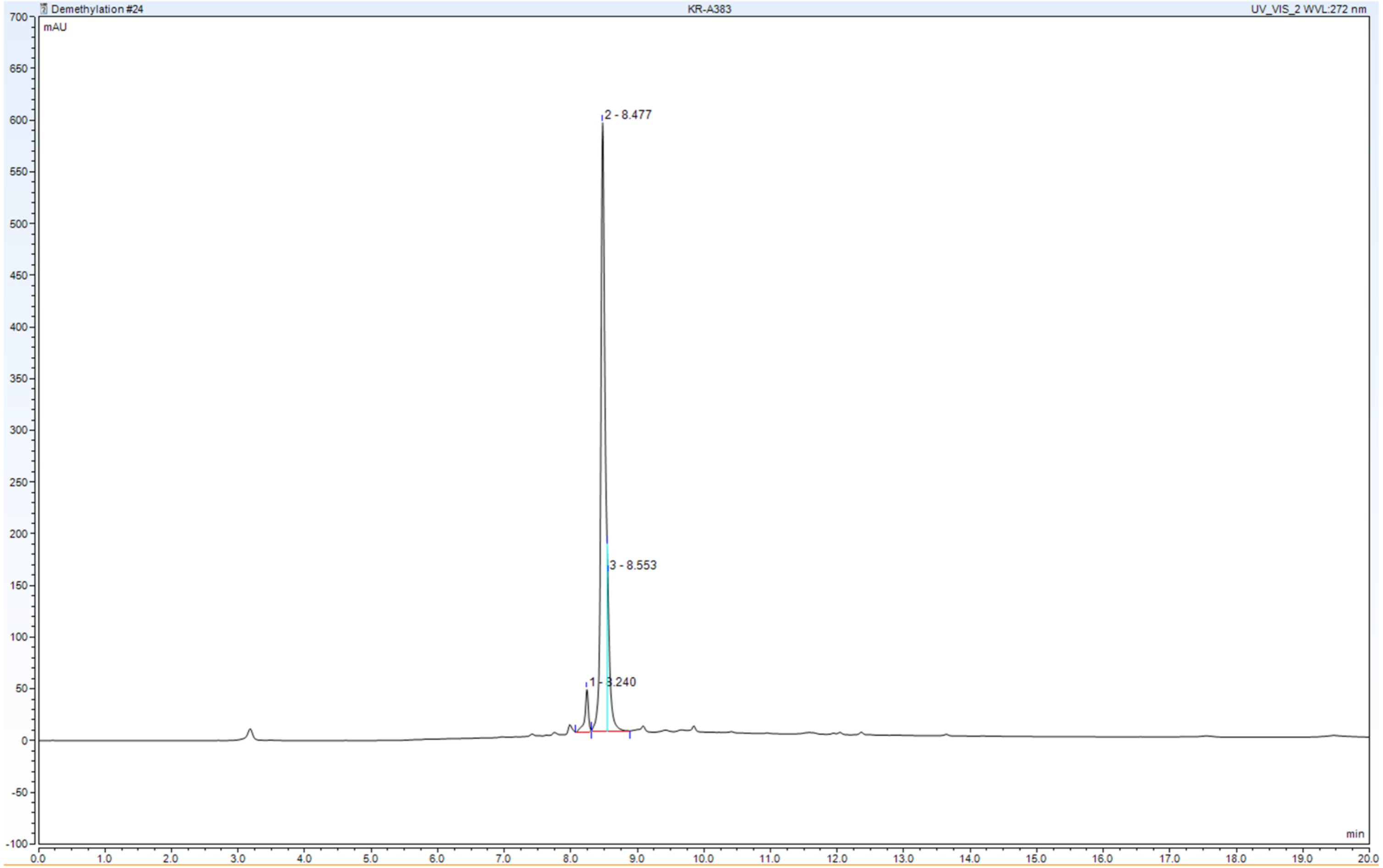
